# Transcriptional profiling of Multiple System Atrophy cerebellar tissue highlights differences between the parkinsonian and cerebellar sub-types of the disease

**DOI:** 10.1101/2020.02.11.944306

**Authors:** Ignazio S. Piras, Christiane Bleul, Isabelle Schrauwen, Joshua Talboom, Lorida Llaci, Matthew D. De Both, Marcus A. Naymik, Glenda Halliday, Conceicao Bettencourt, Janice L. Holton, Geidy E. Serrano, Lucia I. Sue, Thomas G. Beach, Nadia Stefanova, Matthew J. Huentelman

**Affiliations:** Neurogenomics Division, The Translational Genomics Research Institute, Phoenix, AZ, US; The University of Sydney School of Medicine, Sydney, AU; Queen Square Brain Bank for Neurological Disorders, UCL Queen Square Institute of Neurology, London, UK; Civin Laboratory of Neuropathology at Banner Sun Health Research Institute, Sun City, AZ, 85351, US; Department of Neurology, Division of Neurobiology, Medical University of Innsbruck, Innsbruck, AT

**Keywords:** Multiple System Atrophy, RNA sequencing, oligodendrocytes, neurodegeneration

## Abstract

Multiple system atrophy (MSA) is a rare adult-onset neurodegenerative disease of unknown cause, with no effective therapeutic options, and no cure. Limited work to date has attempted to characterize the transcriptional changes associated with the disease, which presents as either predominating parkinsonian (MSA-P) or cerebellar (MSC-C) symptoms. We report here the results of RNA expression profiling of cerebellar white matter (CWM) tissue from two independent cohorts of MSA patients (*n*=66) and healthy controls (HC; *n*=66). RNA samples from bulk brain tissue and from oligodendrocytes obtained by laser capture microdissection (LCM) were sequenced. Differentially expressed genes (DEGs) were obtained and were examined before and after stratifying by MSA clinical sub-type.

We detected the highest number of DEGs in the MSA-C group (*n* = 747) while only one gene was noted in MSA-P, highlighting the larger dysregulation of the transcriptome in the MSA-C CWM. Results from both bulk tissue and LCM analysis of MSA-C showed a downregulation of oligodendrocyte genes and an enrichment for myelination processes with a key role noted for the *QKI* gene. Additionally, we observed a significant upregulation of neuron-specific gene expression in MSA-C and an enrichment for synaptic processes. A third cluster of genes was associated with the upregulation of astrocyte and endothelial genes, two cell types with a key role in inflammation processes. Finally, network analysis in MSA-C showed enrichment for β-amyloid related functional classes, including the known Alzheimer’s disease (AD) genes, *APP* and *PSEN1*.

This is the largest RNA profiling study ever conducted on post-mortem brain tissue from MSA patients. We were able to define specific gene expression signatures for MSA-C highlighting the different stages of the complex neurodegenerative cascade of the disease that included alterations in several cell-specific transcriptional programs. Finally, several results suggest a common transcriptional dysregulation between MSA and AD-related genes despite the clinical and neuropathological distinctions between the two diseases.

## Introduction

Multiple-system atrophy (MSA) is a rare neurodegenerative disorder characterized by autonomic dysfunction, ataxia, and parkinsonism. The prevalence is estimated to be between 1.9 to 4.9 per 100,000 (Bhidayasiri and Ling, 2008; Stefanova et al., 2009). The disease affects both sexes equally with onset typically in the sixth decade of life and with an average survival after diagnosis of less than ten years [53]. There are no effective long-term therapeutic options for the MSA patient, and no cure.

MSA as a unifying diagnostic terminology was developed to encapsulate three neurological entities: striatonigral degeneration, olivopontocerebellar atrophy, and Shy-Drager syndrome (Goedert, 2001; Quinn and Wenning, 1995; Vanacore, 2005; Wakabayashi et al., 1998). Two different clinical subtypes have been described based on the predominating motor features noted during the early stages of the disease: the MSA-P subtype (dominated by parkinsonism) and the MSA-C subtype (dominated by cerebellar ataxia). However, in the later stages of the disease, the phenotypic characteristics of both subtypes are typically noted in the patient [16]. A definitive diagnosis of MSA is obtained through autopsy confirmation of a high density of α-synuclein-containing protein aggregates, known as glial cytoplasmic inclusion (GCI) bodies, in oligodendrocytes along with striatonigral degeneration and/or olivopontocerebellar atrophy (Bhidayasiri and Ling, 2008; Papp et al., 1989; Stefanova et al., 2009).

GCIs are primarily comprised of aggregated α-synuclein, therefore MSA can be classified as an oligodendroglial α-synucleinopathy, which is a point of distinction compared to neuronal α-synucleinopathies like Parkinson’s disease. Interestingly, work investigating the earliest molecular changes associated with MSA has suggested that oligodendrocyte intracellular accumulation of p25α, a protein associated with myelination, may be altered before α-synuclein aggregation is observed [53]. The aggregation of α-synuclein is thought to lead to a disruption of the role of the oligodendrocyte in the process of neuronal myelination leading to microglial activation and subsequent release of mis-folded α-synuclein by the increasingly dysfunctional oligodendrocytes. Neighboring neurons may uptake this extracellular α-synuclein and it could thereby initiate new aggregation inside the neuronal cell. Additionally, the toxic α-synuclein species may spread to neurons in other synaptically-connected brain regions in a prion-like fashion. The lack of effective oligodendrocyte support for the local neurons, and the neuronal effects of the α-synuclein inclusions, eventually results in axonal dysfunction, neuronal cell death, and a reactive astrogliosis [16].

The cause of MSA is not known, however it is generally believed to be sporadic. Several genomic studies have been performed to shed light on the molecular pathogenesis of the disease. Three SNPs located in the α-synuclein gene (*SCNA*) have been associated with the risk of developing MSA [48]. In an independent study conducted by evaluating 32 SNPs in the *SNCA* gene, one SNP associated with MSA and one haplotype associated with the MSA-C subgroup were noted [2]. Whole genome sequencing analysis identified *COQ2* genetic variants associated with both sporadic and familial MSA [37]. However, this finding has not been replicated in other cohorts [49]. In another GWAS, including MSA patients and healthy controls, several SNPs located in different genes (*FBX047*, *ELOVL7*, *EDN1*, and *MAPT*) were found to be potentially associated, but were not significant after multiple test correction [47]. Finally, the presence of an expansion of one allele in *SCA3* (a gene associated with spinocerebellar ataxia) was observed in a patient showing clinical features consistent with MSA-C [38]. Recently, epigenetic modifications, such as DNA methylation changes, have also been identified in neurodegenerative diseases. A recent study reported white matter tissue DNA methylation changes associated with MSA, including changes in *HIP1*, *LMAN2* and *MOBP* [8].

Three different gene expression profiling studies conducted on neuropathologically verified human brain samples have been reported to date. The first study [35] utilized transcriptome profiling by RNA-sequencing of the white and grey matter of the frontal cortex from 6 MSA patients and 6 controls. In the grey matter they detected 5 genes differentially expressed (*HLA-A*, *HLA-B*, *HLA-C*, *TTR* and *LOC389831*). In the white matter they identified 7 genes, including the 3 HLA genes detected in the grey matter. The additional genes were: *HBA1*, *HBA2*, *HBB* and *IL1RL1*.

The *SNCA* gene was detected to be upregulated in both comparisons but it was not statistically significant. They also compared the white matter versus the grey matter in patients, detecting a total of 1,910 differentially expressed genes. A second study was conducted using the same 12 samples, but using strand-specific RNA-sequencing [36], detecting a total of 123 differentially expressed genes. Most detected genes were lincRNAs or un-annotated transcripts. Some of the genes found in the previous study [35] were confirmed; *HBB*, *IL1RL1*, *TTR* and *LOC389831*. Finally, a study determining the differential expression of circular RNA (circRNA) in the MSA frontal cortex was conducted [12], identifying 5 circRNAs produced by backsplicing of the precursor mRNAs from the *IQCK*, *EFCAB11*, *DTNA*, *MAP4K3*, and *MCTP1* genetic loci. No other RNA sequencing studies have been conducted thus far.

In this study we utilized RNA sequencing to characterize the cerebellar white matter transcriptome from neuropathologically verified MSA cases and controls using two independent sample sets and two different profiling technologies.

## Material and methods

Extended methods are reported in the Supplementary Appendix

### Human samples

We analyzed two independent cohorts of postmortem cerebellar white matter (CWM) that included both MSA-P and MSA-C subtypes. Cohort 1 (C1) was obtained from the New South Wales (NSW) brain bank (Sidney, AU) and from the Brain and Body Donation Program (Sun City, AZ) to yield a total of 19 pathologically-proven cases MSA and 10 Healthy Controls (HC) (Table 1A). Cohort 2 (C2) was obtained from the Queen Square Brain Bank for Neurological Disorders (London, UK) and included 48 pathologically proven MSA cases and 47 HC (Table 1B).

**Table 1.**
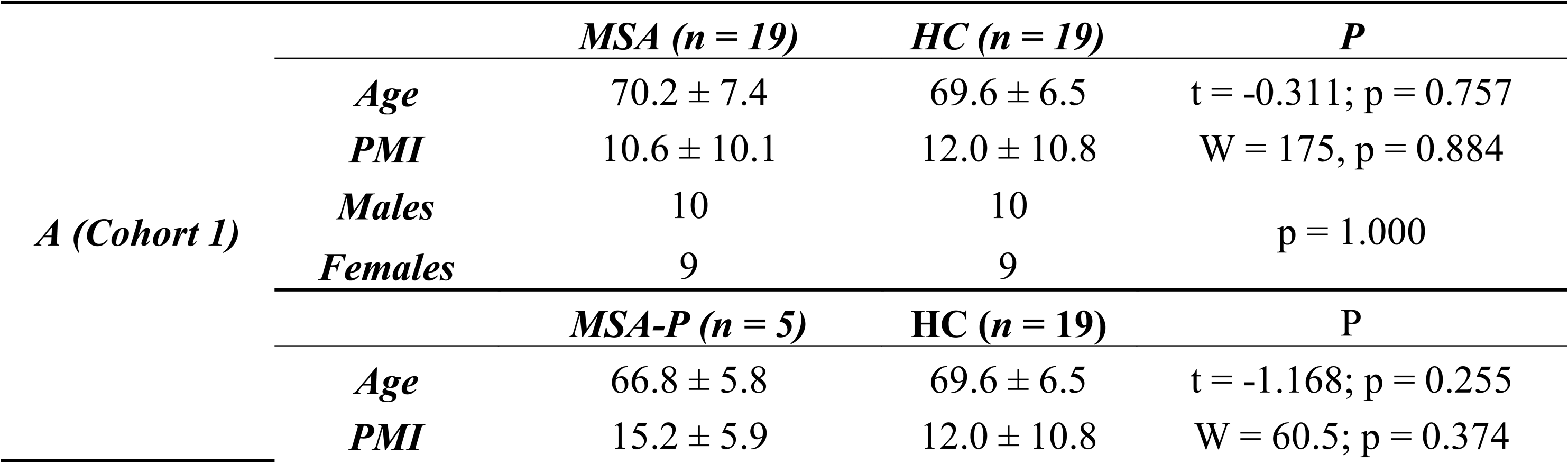
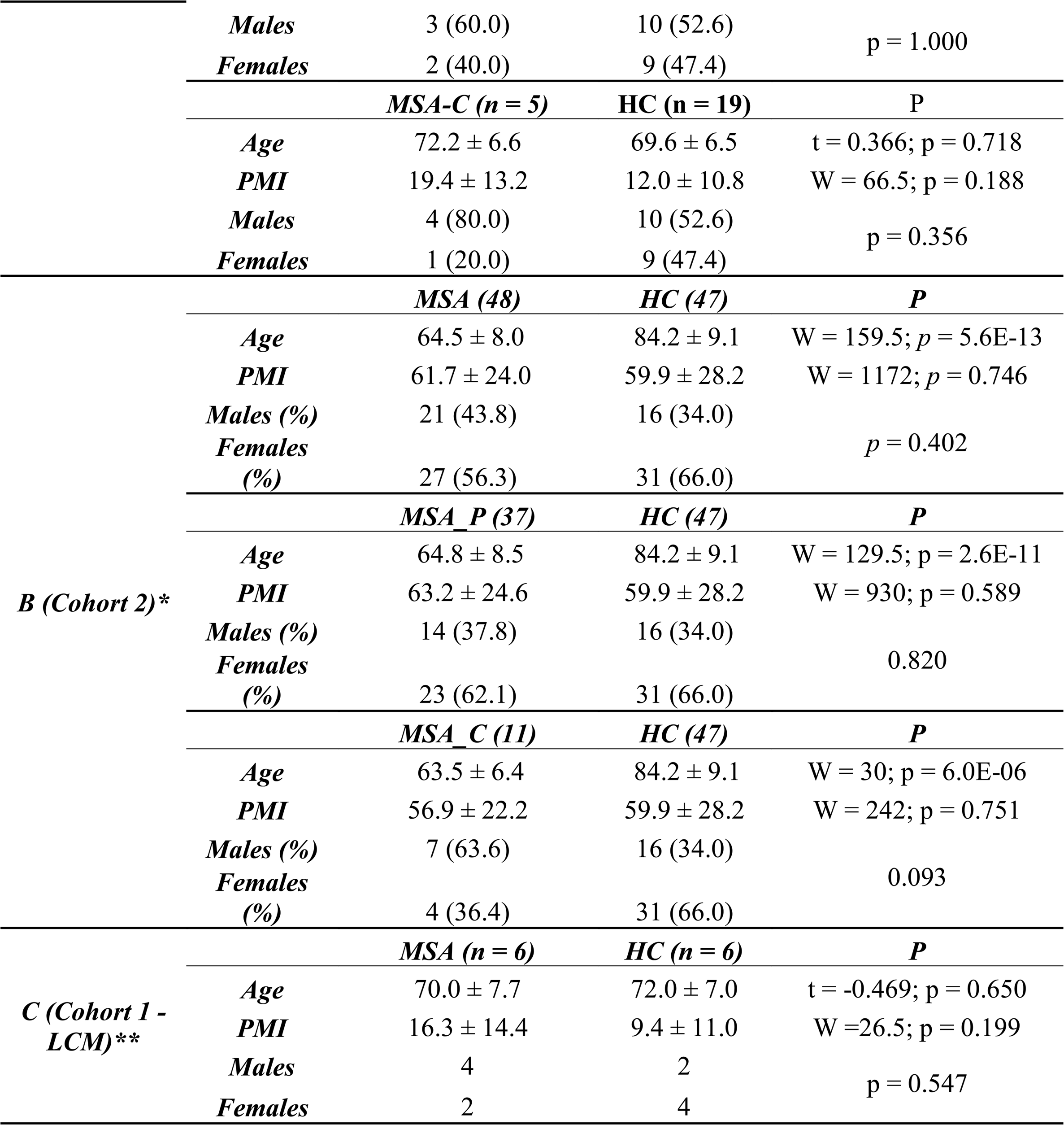
Sample characteristics of the different cohorts analyzed. Differences in age and PMI between cases and controls were assessed using t-test or Wilcoxon test, according to the data distribution. Sex distribution was assessed using the Fisher’s Exact test. * One MSA-P sample was removed after the PCA analysis (Final sample size: MSA = 47; MSA-P = 36; MSA-C = 11; HC = 47) ** Two MSA and one HC samples were removed after PCA analysis (Final sample size: MSA = 4; HC = 5)

### RNA extraction and RNA sequencing

For C1, total DNAse-treated RNA was extracted using the Qiagen RNAeasy kit (Qiagen). Quality was assessed by Bioanalyzer (Agilent). Sequencing libraries were prepared with 250 ng of total RNA using Illumina’s Truseq RNA Sample Preparation Kit v2 (Illumina, Inc.) following the manufacturer’s protocol. The final library was sequenced by 50 bp paired-end sequencing on a HiSeq 2500 (Illumina, Inc.). For C2, total DNAse-treated RNA was extracted in TRI Reagent from ∼5 mg of tissue using Rino Tubes (Next Advance) (TempO-Seq Assay User Guide version 2.0). The final library was sequenced by 50 bp single-end sequencing on a NextSeq 500 (Illumina, Inc.).

### Laser Capture Microdissection (LCM)

Twelve samples (6 MSA, 6 HC) from C1 were used for laser capture microdissection (LCM) of oligodendrocytes from cerebellar white matter (Ordway et al., 2009). Oligodendrocytes were stained by using a modified H&E staining protocol adapted from Ordway et al. [39]. A total of 300 oligodendrocytes per sample were captured using Arcturus CapSure Macro LCM Caps (Applied Biosystems) with the following settings: UV speed at 676 um/s and UV current at 2%. RNA was extracted immediately after cell capture using the Arcturus PicoPure RNA Isolation Kit (Applied Biosystems). For library preparation the SMARTer® Stranded Total RNA-Seq Kit - Pico Input (Clontech/Takara) was used. Samples were sequenced (2 x 75 bp paired-end run) on the Illumina HiSeq2500.

### Data Analysis

Quality controls on FASTQ files were conducted using MultiQC software v0.9 [22]. The reads were aligned to the human reference genome (GRCh37) using the STAR software v2.5 [20] and summarized as gene-level counts using featureCounts 1.4.4 [32]. For both datasets (C1 and C2) PCA analysis was used to assess the presence of outliers and to detect any batch effects. Four samples were deemed to be outliers and were removed (detailed below in Results). Gene expression differential analyses between MSA cases and HC were conducted using the R package DESeq2 v1.14.1 (Love et al., 2014), including age, sex (only C2), PMI and sample source (only C1) as covariates. Sex was not included as a covariate for C2 because the sexes were balanced and sample source was not included as a covariate for C1 because the tissue sources were balanced. The p-values were corrected for multiple testing using the False Discovery Rate (FDR) method (Benjamini and Hochberg, 1995), considering as significant all the genes with adjusted p-value (adj-p) < 0.05.

The results from the two cohorts were combined using a meta-analysis approach based on the weighted-Z method [62] as implemented in the R-package *survcomp* [50]

### Cell specific Expression

We classified the genes detected in the differential expression analysis using an external database of expression values from different types of cells isolated from mouse cerebral cortex [63]. We computed an enrichment score for each cell type and gene, assigning each gene to a specific cell type according to the relative expression in the other cell types. The enrichment of cell specific genes was investigated across DEGs and co-expression modules using a hypergeometric statistic (R function *phyper*).

### Enrichment and functional Network analysis

Lists of DEGs were analyzed for Gene Ontology (GO) enrichment using the R-package *anRichmentMethods*, adjusting the *p*-value with the FDR method. The same gene lists were also analyzed using *HumanBase* (https://hb.flatironinstitute.org/gene), constructing tissue-specific functional networks [25].

The enrichment of Alzheimer’s disease genes in MSA was conducted using the data from the Accelerated Medicine Partnership – Alzheimer’s Disease (AMP-AD) portal. We downloaded the differential expression results from 7 different brain regions from the Mayo, Mount Sinai and ROSMAP cohorts [3, 6, 61]. Specifically, the brain regions included were: temporal cortex (TCX), cerebellum (CBE), dorsolateral pre-frontal cortex (DLPFC), inferior frontal gyrus (IFG), frontal pole (FP), parahippocampal gyrus (PHC), and superior temporal gyrus (STG). The DEGs from these 7 brain regions were used as gene set references for the list of MSA genes ranked by log2 FC. The analysis was conducted using R-package *fgsea* adjusting the p-values with the FDR method.

### Weighted Correlation Network Analysis

We conducted Weighted Correlation Network Analysis (WGCNA) in the MSA-C cohorts with the aim of identifying modules of co-expressed genes associated with the disease and enriched for specific biological processes [31]. We computed the co-expression networks using the data from C1 and then we estimated the module preservation in C2, using only MSA-C and HC. The analysis was conducted using the *WGCNA* R-package [31]. Genes for both C1 and C2 with less than 10 average counts were filtered out due to low expression and data were normalized using the *vst* function of the DESeq2 package [33]. The matrix of expression values was adjusted for age, sex, source and PMI using the function *removeBatchEffect* as implemented in the limma R-package [45]. Finally, we filtered out the 50% of genes having lower Median Absolute Deviation (MAD). We generated a signed co-expression network for C1 using the function *blockwiseModules*, with the option *mergeCutHeight* = 0.25. Then, we computed the module eigengenes and we investigated their relationship with disease status using a linear model as implemented in the *limma* package. We calculated the module membership and gene-trait significance (MSA-C disease status) with the goal of ranking genes in each co-expression modules. Modules associated with disease status were further investigated using GO enrichment analysis. The enrichment for genes expressed in specific cell types was conducted using as reference gene sets the gene specifically expressed in the 5 cell types from Zhang et al. [63] and test sets including all of the genes ranked by module memberships for the module associated with the disease status. Finally, we checked the module preservation in C2 using the *modulePreservation* function with 1,000 permutations. Relevant coexpression networks were exported and visualized using *Cytoscape v3.7.2* [51].

## Results

### Quality controls

For C1 (Illumina), we sequenced a total of 470 Million (M) reads (average: 12.4 M; range: 3.8 – 32.6 M) with a 76.7% average mapping rate. PCA analysis did not show the presence of outliers (Fig. S1). For C2 (TempO-Seq) we sequenced a total of 162 M reads (average: 1.7 M; range: 0.12 – 3.8 M), with a 90.2% average mapping rate. PCA analysis showed the presence of one outlier in the C2 group and it was removed from all subsequent analyses. The final sample size was: MSA = 47 and HC = 47 (Fig. S2A and Fig. S2B). For the LCM sample (a subsample from C1) we sequenced a total of 353 M reads (average: 29.4 M; range: 23.4 – 33.3 M) with an average 64.4% mapping rate. We detected the presence of three outliers that were also removed. The final sample size used for the differential analysis from the LCM dataset was: MSA = 4 and HC = 5 (Fig. S3A and Fig. S3B).

### Differential expression results: bulk tissue human samples (MSA, MSA-P and MSA-C)

Differentially expressed genes (DEGs) were obtained by combining the results from both cohorts using a meta-analysis approach. Details about the specific results for each cohort are reported in Tables S1-S3 and Fig. S4.The comparison of the log2 FC obtained in the differential analyses for the two independent cohorts for MSA, MSA-P and MSA-C was statistically significant (ρ range *=* 0.204 – 0.456; p < 2.2E-16). The largest correlation coefficient (ρ = 0.456) was obtained for the MSA-C subtype probably due to the larger significance and effect size of the genes detected (Fig. S5).

After p-value combination, we obtained a set of DEGs ranging from 1 (MSA-P) to 747 (MSA-C) depending on the MSA sub-type (Fig. 1A – 1C). The complete results are reported in Tables S4-S6.The top 3 DEGs for MSA in general were *ACTN1, EMP1 and NFIL3* (adj *p < 0.01*; all upregulated). In the MSA-P clinical sub-type we detected only one DEG (*GPNMB*), whereas in MSA-C the top genes were: *PGAM2*, *ST5*, *STON1*, *RFTN1*, *ACTN1* and *MMP14* (adj *p* < 1.0E-04; all upregulated) (Table 2; Fig. 2). We explored the differential expression between SND vs HC, and OPCA vs HC, detecting a total of 7 and 58 genes, respectively. *MLPH*, detected in SND, was also detected when analyzing the clinical subtype MSA-P in C2, whereas a total of 47 genes detected in OPCA were also observed in the MSA-C clinical subtype in C2 (Table S7; Fig. S6). Correlation of the log2 FC between the differential analysis for clinical and neuropathological classification criteria were ρ = 0.622 (MSA-P/SND) and ρ = 0.830 (both p < 2.2E-16) (Fig. S7).

**Figure 1.**
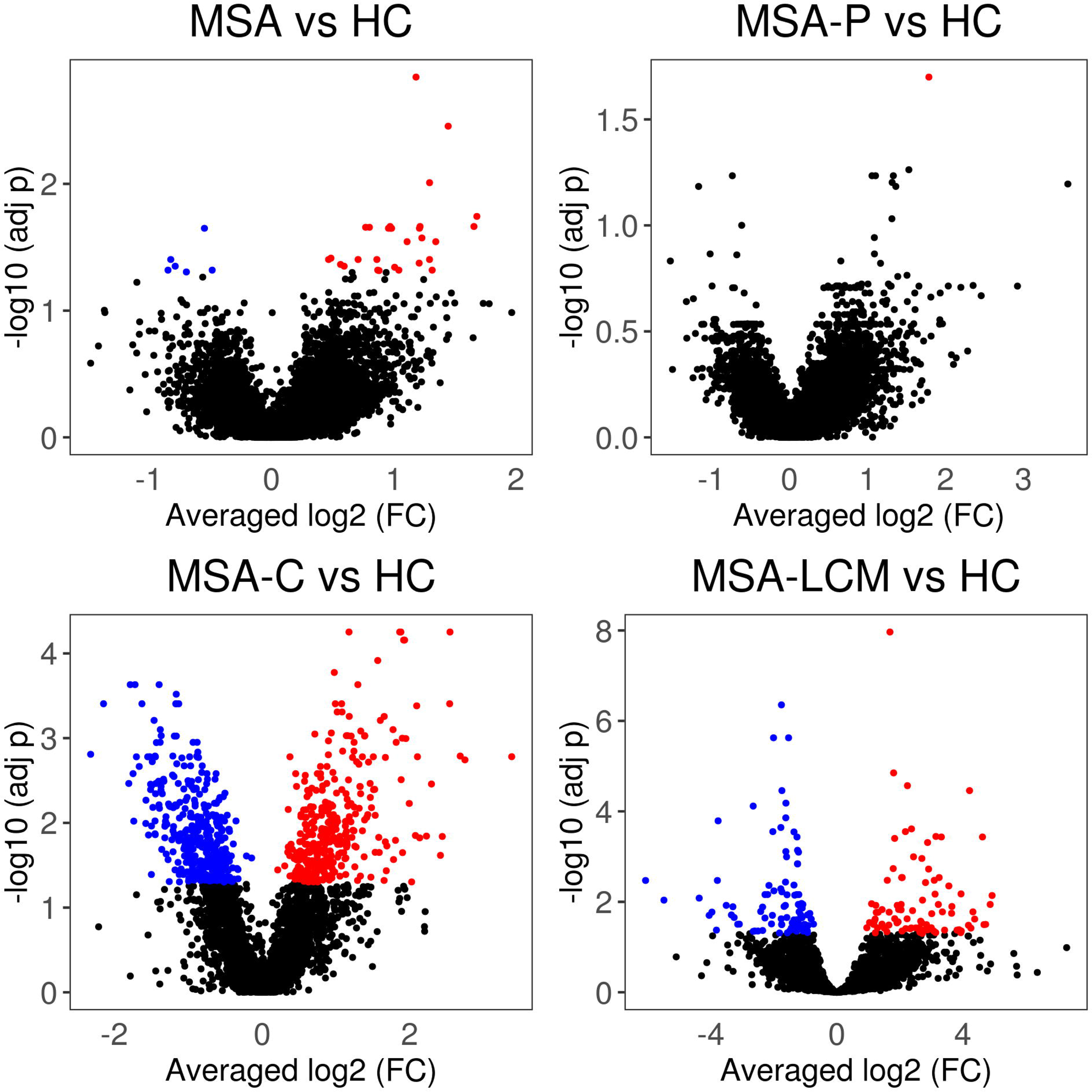
Volcano plots representing the differential expression results after p-value combination (excluding LCM dataset). In red and blue upregulated and downregulated genes, respectively. (A): MSA (B): MSA-P (C): MSA-C (D): Oligodendrocytes

**Figure 2.**
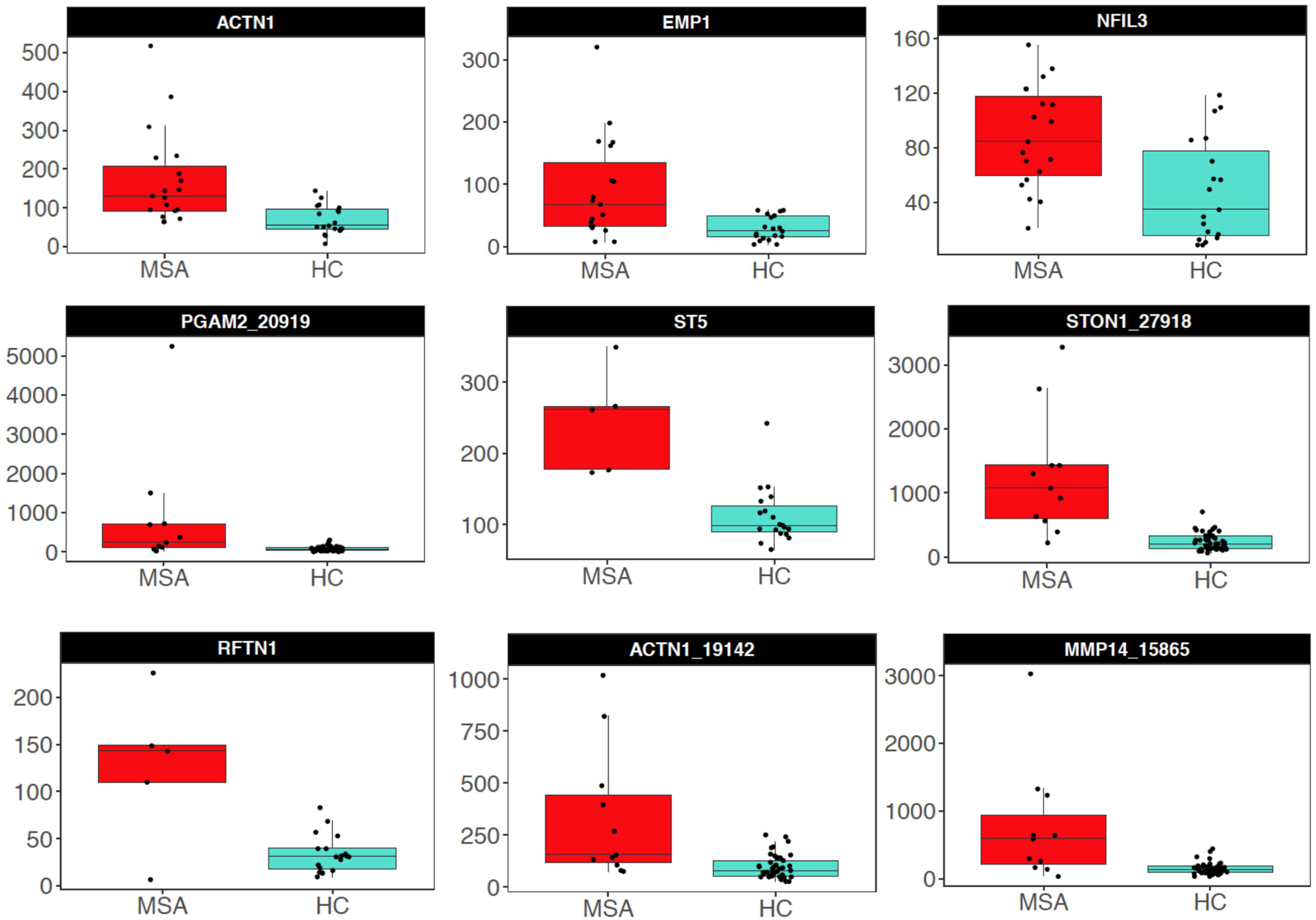
Top differentially expressed genes found in: MSA MSA-C

**Table 2.**
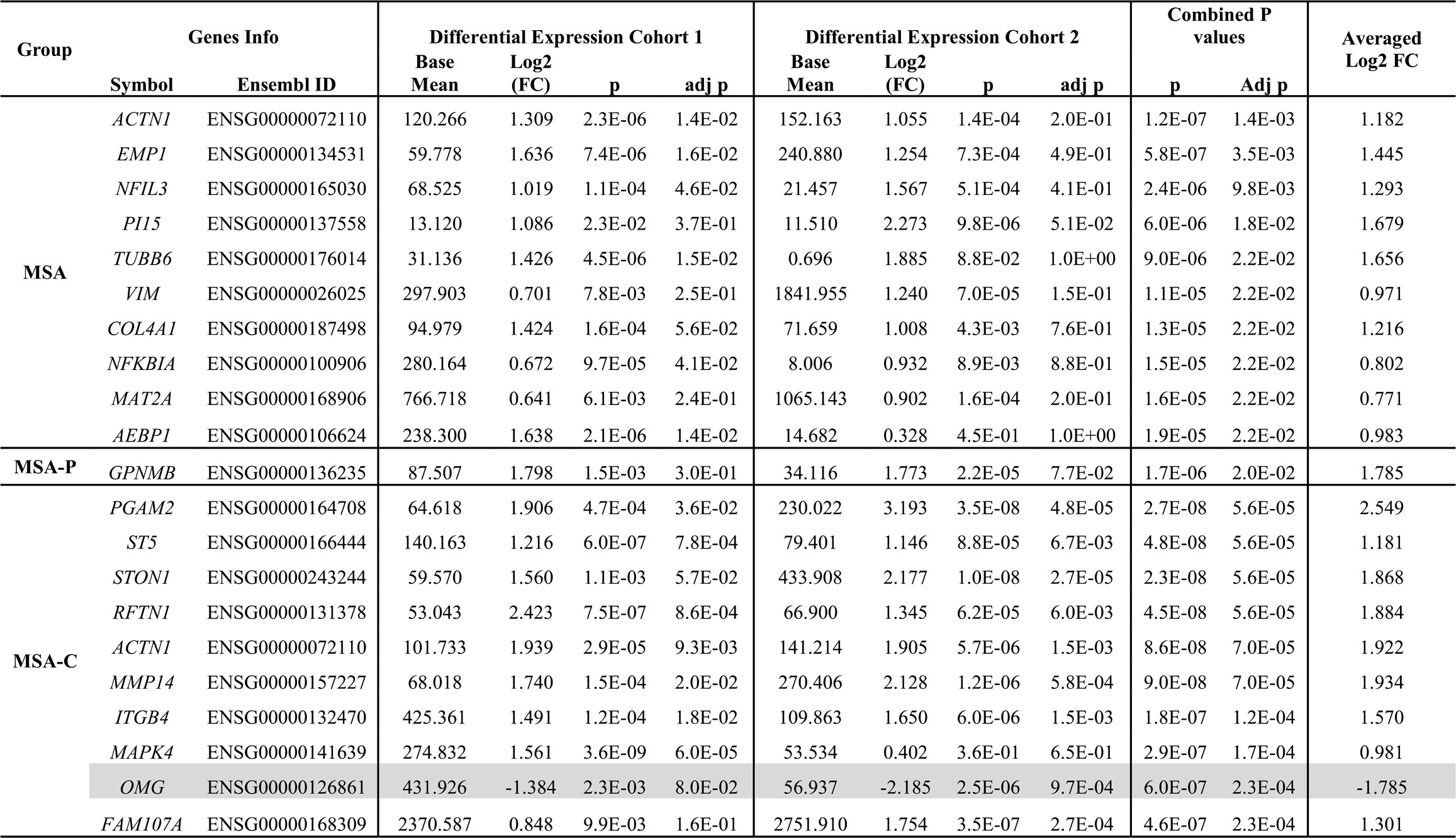
Top genes for the different MSA subtypes after p-value combination. Downregulated genes are reported in grey.

We explored the functional significance of the DEGs by applying a functional network analysis specific for the cerebellum and a GO enrichment analysis. Using the 35 MSA DEGs (Table S4) we detected a small network with 2 modules enriched for “cell-cell adhesion” (*SELL* and *BCL6* genes) and “angiogenesis” (*COL4A1* and *COL4A2* genes) (both *q* < 0.01) (Fig. S8). The GO analysis yielded significant enrichment of the Biological Process “collagen-activated signaling pathway” (adj *p* = 0.030; genes: *COL4A1*, *COL4A2*, *ITGA11).* Using all of the 747 MSA-C DEGs (Table S6) we detected a large network including 9 different modules (Fig. 3). We observed the highest enrichment significance in module 1 (M1) which was amyloid-β metabolism (top GO process: q = 5.3E-05, Table S8), including the Alzheimer’s disease (AD) relevant genes: *APP*, *PSEN1*, *CLU, ROCK2* and *DYRK1*. The central role of *APP* was confirmed by a separate protein-protein interaction analysis showing this gene as the most important hub in a network that included 30% of the DEGs generated using *WEBGESTALT* [60] (Fig. S9). The second highest significance was reached in module 2 (M2) for respiratory chain complex assembly (top GO process: *q* = 8.2E-03) (Table 3, Table S8). With the GO analysis we detected 625 significant functional classes, mostly related to cellular and cytoplasmatic components, neuro and gliogenesis (Fig. S10).

**Figure 3.**
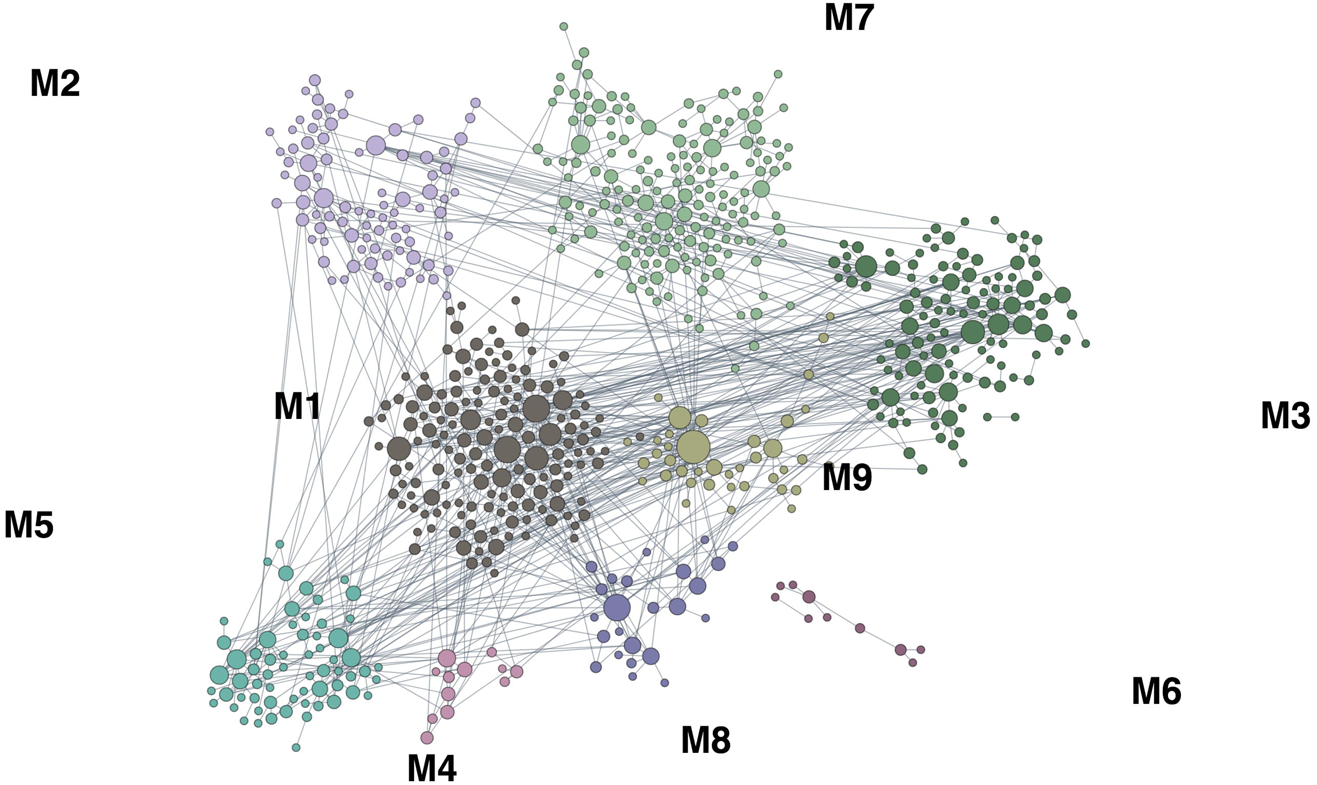
Results of the functional network analysis on MSA-C DEGs. Module 1 was enriched for amyloid-β metabolism (q = 5.3E-05) including key AD genes as: *APP*, *PSEN1*, *CLU*, *ROCK2* and *DYRK1*.

**Table 3.**
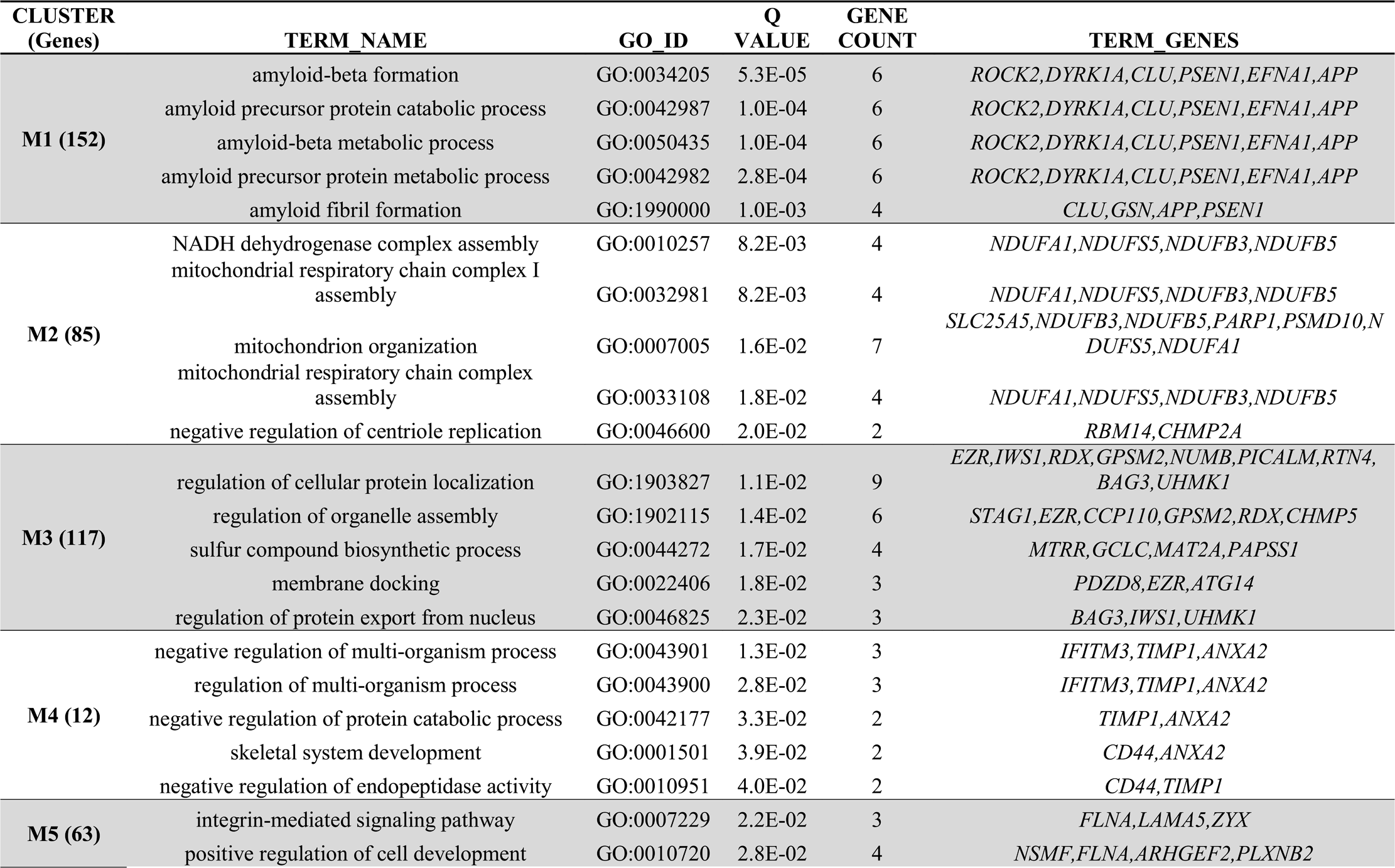
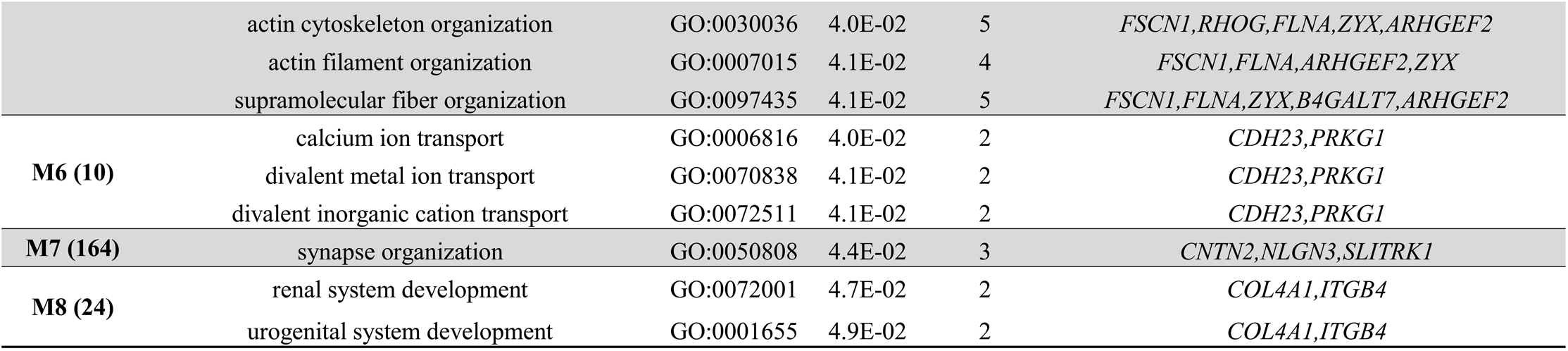
Top results of the functional module discovery analysis using the DEGs identified in MSA-C

| CLUSTER<br>(Genes) | TERM_NAME | GO_ID | Q<br>VALUE | GENE<br>COUNT | TERM_GENES |
| --- | --- | --- | --- | --- | --- |
| **M1 (152)** | amyloid-beta formation | GO:0034205 | 5.3E-05 | 6 | *ROCK2,DYRK1A,CLU,PSEN1,EFNA1,APP* |
|  | amyloid precursor protein catabolic process | GO:0042987 | 1.0E-04 | 6 | *ROCK2,DYRK1A,CLU,PSEN1,EFNA1,APP* |
|  | amyloid-beta metabolic process | GO:0050435 | 1.0E-04 | 6 | *ROCK2,DYRK1A,CLU,PSEN1,EFNA1,APP* |
|  | amyloid precursor protein metabolic process | GO:0042982 | 2.8E-04 | 6 | *ROCK2,DYRK1A,CLU,PSEN1,EFNA1,APP* |
|  | amyloid fibril formation | GO:1990000 | 1.0E-03 | 4 | *CLU,GSN,APP,PSEN1* |
| **M2 (85)** | NADH dehydrogenase complex assembly | GO:0010257 | 8.2E-03 | 4 | *NDUFA1,NDUFS5,NDUFB3,NDUFB5* |
|  | mitochondrial respiratory chain complex I assembly | GO:0032981 | 8.2E-03 | 4 | *NDUFA1,NDUFS5,NDUFB3,NDUFB5* |
|  | mitochondrion organization | GO:0007005 | 1.6E-02 | 7 | *SLC25A5,NDUFB3,NDUFB5,PARP1,PSMD10,NDUFS5,NDUFA1* |
|  | mitochondrial respiratory chain complex assembly | GO:0033108 | 1.8E-02 | 4 | *NDUFA1,NDUFS5,NDUFB3,NDUFB5* |
|  | negative regulation of centriole replication | GO:0046600 | 2.0E-02 | 2 | *RBM14,CHMP2A* |
| **M3 (117)** | regulation of cellular protein localization | GO:1903827 | 1.1E-02 | 9 | *EZR,IWS1,RDX,GPSM2,NUMB,PICALM,RTN4,BAG3,UHMK1* |
|  | regulation of organelle assembly | GO:1902115 | 1.4E-02 | 6 | *STAG1,EZR,CCP110,GPSM2,RDX,CHMP5* |
|  | sulfur compound biosynthetic process | GO:0044272 | 1.7E-02 | 4 | *MTRR,GCLC,MAT2A,PAPSS1* |
|  | membrane docking | GO:0022406 | 1.8E-02 | 3 | *PDZD8,EZR,ATG14* |
|  | regulation of protein export from nucleus | GO:0046825 | 2.3E-02 | 3 | *BAG3,IWS1,UHMK1* |
| **M4 (12)** | negative regulation of multi-organism process | GO:0043901 | 1.3E-02 | 3 | *IFITM3,TIMP1,ANXA2* |
|  | regulation of multi-organism process | GO:0043900 | 2.8E-02 | 3 | *IFITM3,TIMP1,ANXA2* |
|  | negative regulation of protein catabolic process | GO:0042177 | 3.3E-02 | 2 | *TIMP1,ANXA2* |
|  | skeletal system development | GO:0001501 | 3.9E-02 | 2 | *CD44,ANXA2* |
|  | negative regulation of endopeptidase activity | GO:0010951 | 4.0E-02 | 2 | *CD44,TIMP1* |
| **M5 (63)** | integrin-mediated signaling pathway | GO:0007229 | 2.2E-02 | 3 | *FLNA,LAMA5,ZYX* |
|  | positive regulation of cell development | GO:0010720 | 2.8E-02 | 4 | *NSMF,FLNA,ARHGEF2,PLXNB2* |
|  | actin cytoskeleton organization | GO:0030036 | 4.0E-02 | 5 | *FSCN1,RHOG,FLNA,ZYX,ARHGEF2* |
|  | actin filament organization | GO:0007015 | 4.1E-02 | 4 | *FSCN1,FLNA,ARHGEF2,ZYX* |
|  | supramolecular fiber organization | GO:0097435 | 4.1E-02 | 5 | *FSCN1,FLNA,ZYX,B4GALT7,ARHGEF2* |
| **M6 (10)** | calcium ion transport | GO:0006816 | 4.0E-02 | 2 | *CDH23,PRKG1* |
|  | divalent metal ion transport | GO:0070838 | 4.1E-02 | 2 | *CDH23,PRKG1* |
|  | divalent inorganic cation transport | GO:0072511 | 4.1E-02 | 2 | *CDH23,PRKG1* |
| **M7 (164)** | synapse organization | GO:0050808 | 4.4E-02 | 3 | *CNTN2,NLGN3,SLITRK1* |
| **M8 (24)** | renal system development | GO:0072001 | 4.7E-02 | 2 | *COL4A1,ITGB4* |
|  | urogenital system development | GO:0001655 | 4.9E-02 | 2 | *COL4A1,ITGB4* |

### Enrichment of AD genes in MSA-C

After we observed the presence of the AD-related process (amyloid-β metabolism) and genes in MSA-C, we tested the enrichment of AD genes in MSA-C. We used data from AMP-AD, running an enrichment analysis by brain region using as reference sets the DEGs from each brain region. The results showed a significant enrichment of TCX (adj p = 7.4E-05) and PHG (adj *p* = 2.0E-02) AD DEGs among upregulated MSA-C genes (Fig. S11) which were also confirmed when we used more conservative cutoffs to select AD genes (adj *p* < 0.01, < 0.001, and < 0.0001) (Table S9A). As further validation, we used the less variable genes between AD and non-demented controls (ND) (adj *p* > 0.950). As expected, we did not observe any significant enrichment of TCX or PHG AD DEGs genes (Table S9B). We compared the DEGs detected in MSA-C, with the DEGs detected in TCX and PHG, only selecting genes having the same log2 FC direction, considering the comparison: affected vs non-affected. We detected 201 genes in TCX, 152 in PHG and 103 common between both regions TCX, PHG and MSA-C (Table S10).

### Differential expression in LCM oligodendrocytes

We detected a total of 187 differentially expressed genes in oligodendrocytes (90 upregulated and 97 downregulated) (Fig. 1D). Details for the complete list of genes are reported in Table S11. The top 4 significant genes (adj *p* < 1.0E-05) were: *GGCX*, *MOCS1*, *NF1* and *LINC01572* (Table 4). Using the functional module discovery analysis we detected a network including 4 modules (72 genes in total) enriched for telomere maintenance (M1: *q* = 1.9E-03; genes: *PTGES3* and *WRAP53*) and ncRNA processing (M2: *q* = 0.0025; genes: *DIMT1*, *INTS8*, and *MTREX*). Modules 3 and 4 are weakly enriched for immune processes and cell growth (*q* < 0.05) (Fig. S12). Using the GO analysis in the complete gene list, we detected a significant enrichment in the myelination process mostly due to downregulated genes (Fig. S13).

**Table 4.**
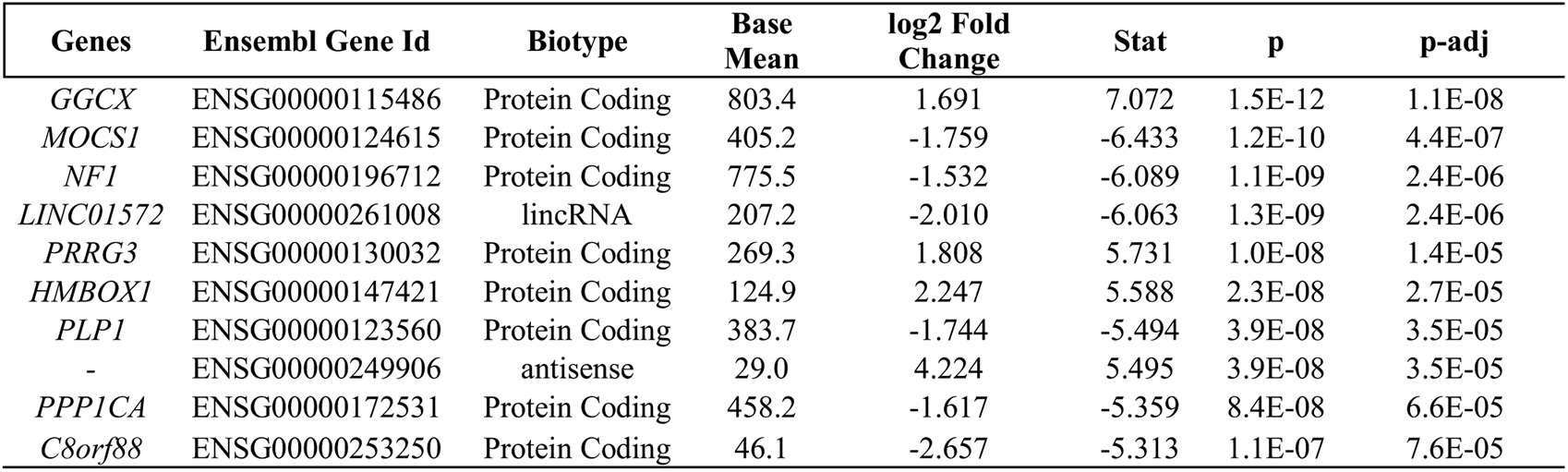
Top genes differentially expressed in oligodendrocytes in MSA vs HC

| **Genes** | **Ensembl Gene Id** | **Biotype** | **Base Mean** | **log2 Fold Change** | **Stat** | **p** | **p-adj** |
| --- | --- | --- | --- | --- | --- | --- | --- |
| *GGCX* | ENSG00000115486 | Protein Coding | 803.4 | 1.691 | 7.072 | 1.5E-12 | 1.1E-08 |
| *MOCS1* | ENSG00000124615 | Protein Coding | 405.2 | -1.759 | -6.433 | 1.2E-10 | 4.4E-07 |
| *NF1* | ENSG00000196712 | Protein Coding | 775.5 | -1.532 | -6.089 | 1.1E-09 | 2.4E-06 |
| *LINC01572* | ENSG00000261008 | lincRNA | 207.2 | -2.010 | -6.063 | 1.3E-09 | 2.4E-06 |
| *PRRG3* | ENSG00000130032 | Protein Coding | 269.3 | 1.808 | 5.731 | 1.0E-08 | 1.4E-05 |
| *HMBOX1* | ENSG00000147421 | Protein Coding | 124.9 | 2.247 | 5.588 | 2.3E-08 | 2.7E-05 |
| *PLP1* | ENSG00000123560 | Protein Coding | 383.7 | -1.744 | -5.494 | 3.9E-08 | 3.5E-05 |
| - | ENSG00000249906 | antisense | 29.0 | 4.224 | 5.495 | 3.9E-08 | 3.5E-05 |
| *PPP1CA* | ENSG00000172531 | Protein Coding | 458.2 | -1.617 | -5.359 | 8.4E-08 | 6.6E-05 |
| *C8orf88* | ENSG00000253250 | Protein Coding | 46.1 | -2.657 | -5.313 | 1.1E-07 | 7.6E-05 |

### Bioinformatic-based cell specific expression profiling

We classified the DEGs obtained in the MSA-C group according to their expression in five brain cell types [63]. We selected only the MSA-C results because the large number of DEGs makes it possible to identify robust cell-specific upregulation/downregulation trends. Most of the DEGs were not cell specific (“mixed”: 74.7% of the total DEGs), whereas the remaining genes were: astrocyte (6.6%), oligodendrocyte (5.8%), endothelial cell (5.1%), neuron (4.1%) and microglia (3.7%) specific. We found a significant overrepresentation of astrocyte and oligodendrocyte genes (both: adj *p* = 2.9E-04). (Fig. S14). We observed a strong downregulation of oligodendrocyte genes and upregulation of microglia, neuron and astrocyte genes (Fig. 4A). To investigate if these patterns are disease specific, we compared the log2 FC of genes differentially expressed (adj *p* < 0.05) with those non-differentially expressed (adj *p* ≥ 0.05) for each cell type. We observed the highest significance for oligodendrocyte (downregulated in MSA) and neuronal genes (upregulated in MSA) (*p* < 0.001). Similar results were obtained when we relaxed the gene inclusion cutoff to adj *p* < 0.10 (Fig. S15). We conducted GO enrichment analysis on the cell-specific DEGs. The highest significance was reached for oligodendrocyte genes, enriched for myelination and oligodendrocyte development processes. Astrocytes were enriched for transport of ion across the membrane, plasma membrane components, and ATPase complex (*FDR* < 0.01). Endothelial cell genes were enriched for cell migration and angiogenesis. Neuronal genes were enriched for neurogenesis and post-synapse organization (Fig. 5; Table S12).

**Figure 4.**
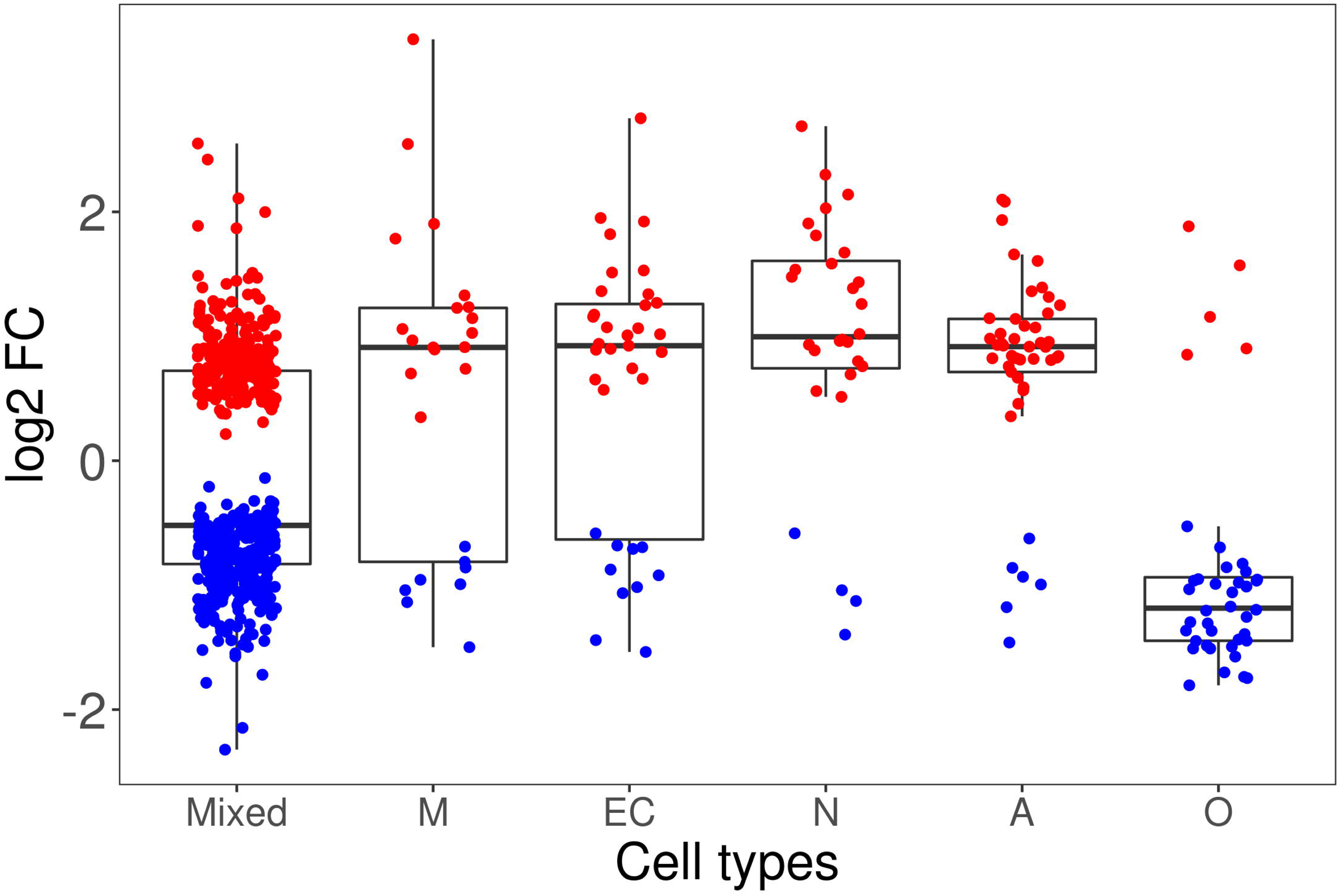
DEGs Log2 FC distribution across the cell-specific genes classes. Upregulated and downregulated genes in MSA-C are in red and blue, respectively.

**Figure 5.**
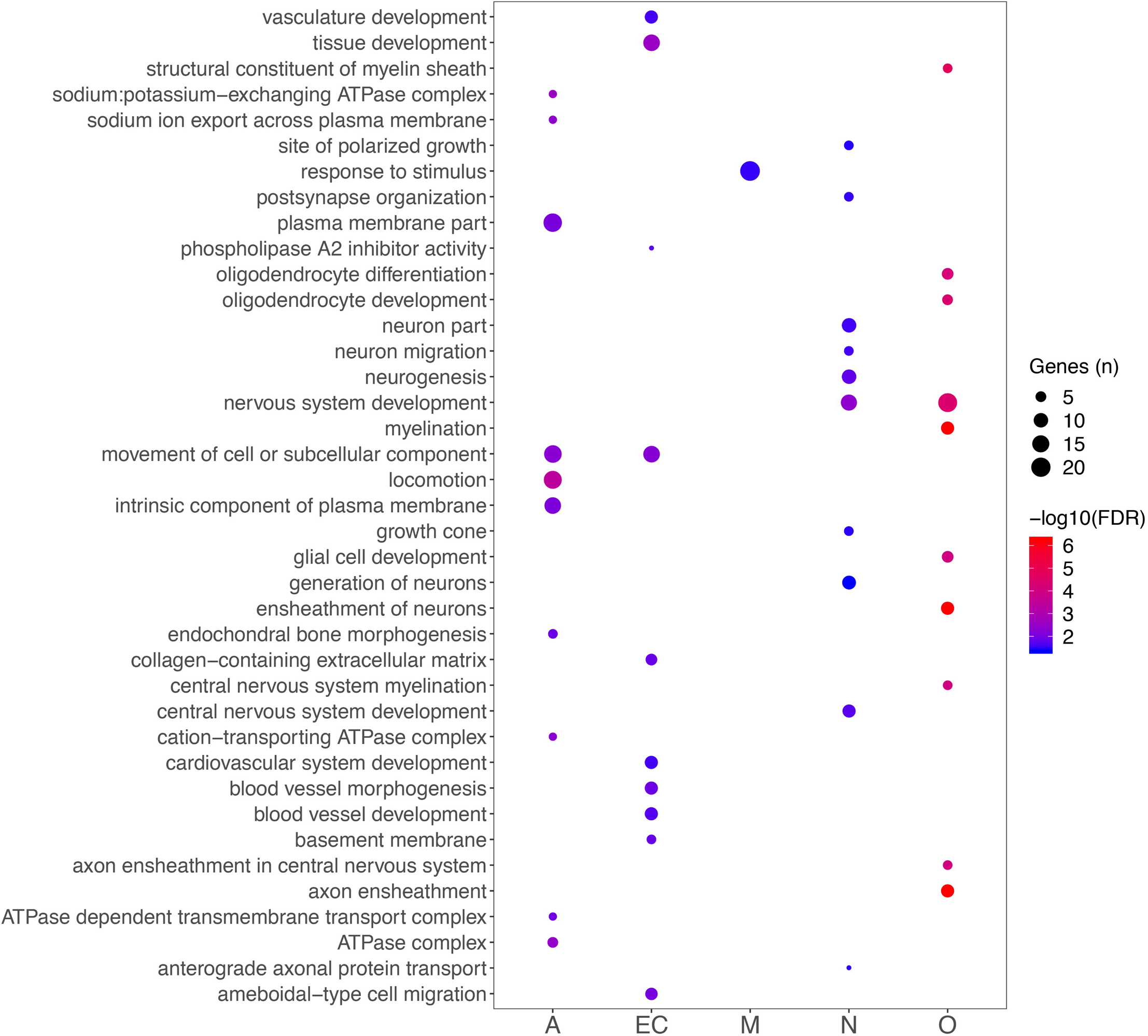
Dot plot reporting the top 10 GO functional classes enriched in each cell-specific gene class.

### WGCNA analysis

Considering the large number of DEGs for the MSA-C subtype, we further investigated this group using WGCNA analysis. We computed a coexpression network using the data from C1 and validated the results in C2 by means of the module preservation analysis. After filtering (see Methods), a network was generated using the 7,650 genes with larger MAD using “9” as threshold power (Fig. S16). We obtained nine co-expression modules in total including 2,675 genes (35.0% of the genes), whereas the remaining were not significantly co-expressed and then were included in the grey module (Fig. 6A). The number of the genes in each module ranged from 78 (magenta) to 917 (turquoise). In Fig. S17 we show the heatmap and the dendrogram indicating the correlation between modules. A total of 4 modules (yellow, green, brown and blue) were associated with disease status, all showing an increase in MSA-C with the exception of the blue module (Fig. 6B). The number of genes in these 4 modules ranged from 160 (green) to 485 (blue). Two of the significant modules (brown and green) were highly correlated with each other (Fig. S17). We computed the module membership (correlation of each gene with the module eigengenes), and the gene-trait significance (correlation with disease status). As expected, the gene-trait significance was highly correlated with the log2 FC (*r* = 0.846; *p* < 2.2E-16). We represented the correlation of the module membership with gene-trait significance in the scatterplots in Fig. S18 and Fig. S19. As expected, we detected a significant positive correlation for the 4 modules associated with MSA (yellow, green, brown, and blue in the Figures) but not for the others (not shown). The genes for these 4 significant modules are reported in Tables S13 ranked by module membership p-value. The most important hubs for the 4 modules were: *TGFB2* (yellow), *SYNGAP1* (green), *TIAM1* (brown) and *QKI* (blue). These networks are represented in Fig. 7.

**Figure 6.**
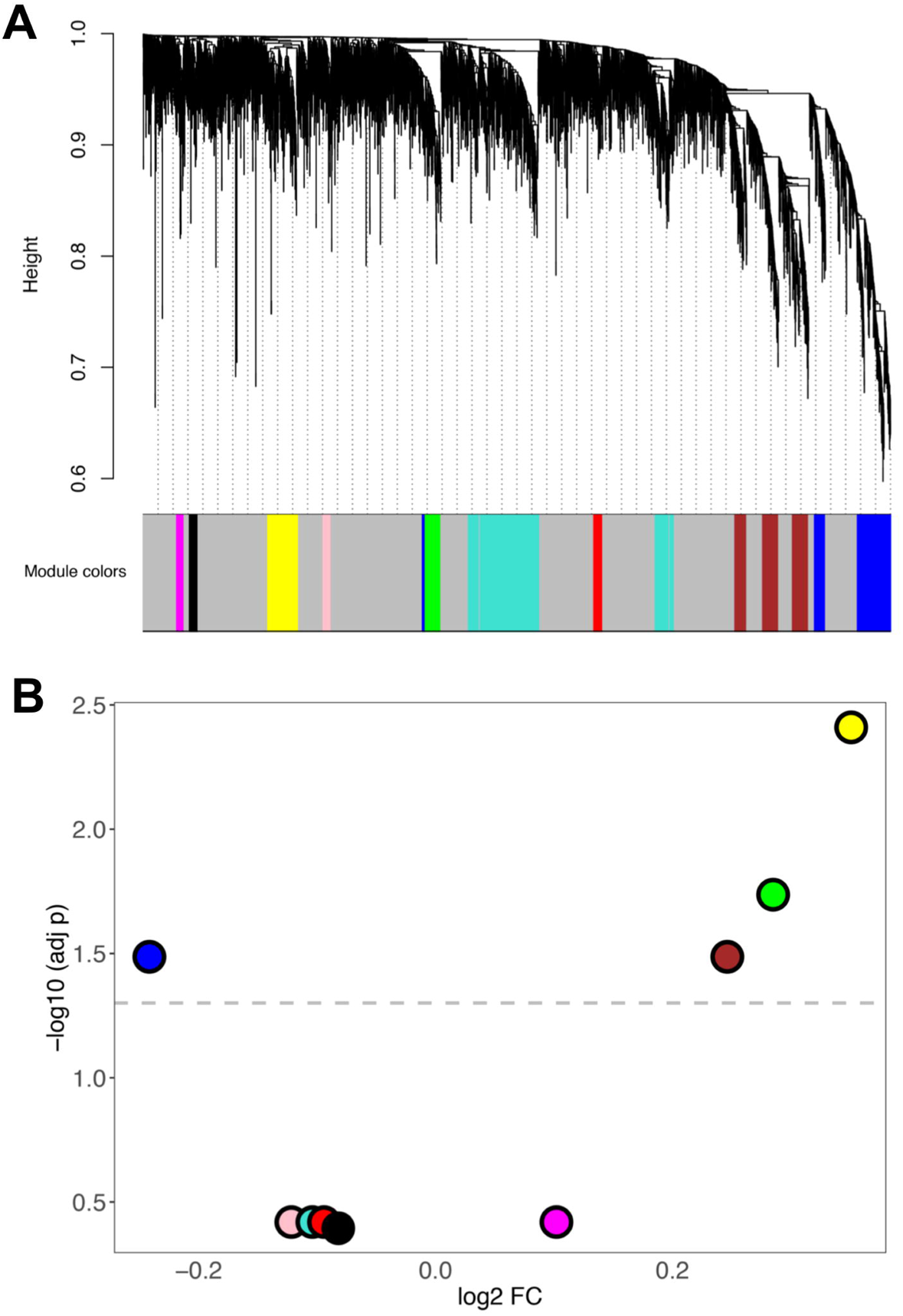
WGCNA analysis (A) Cluster dendrogram showing the 9 coexpression modules detected in MSA-C (C1) (B) Volcano plot representing the results of the differential expression of the eigengene modules between MSA-C vs HC.

**Figure 7.**
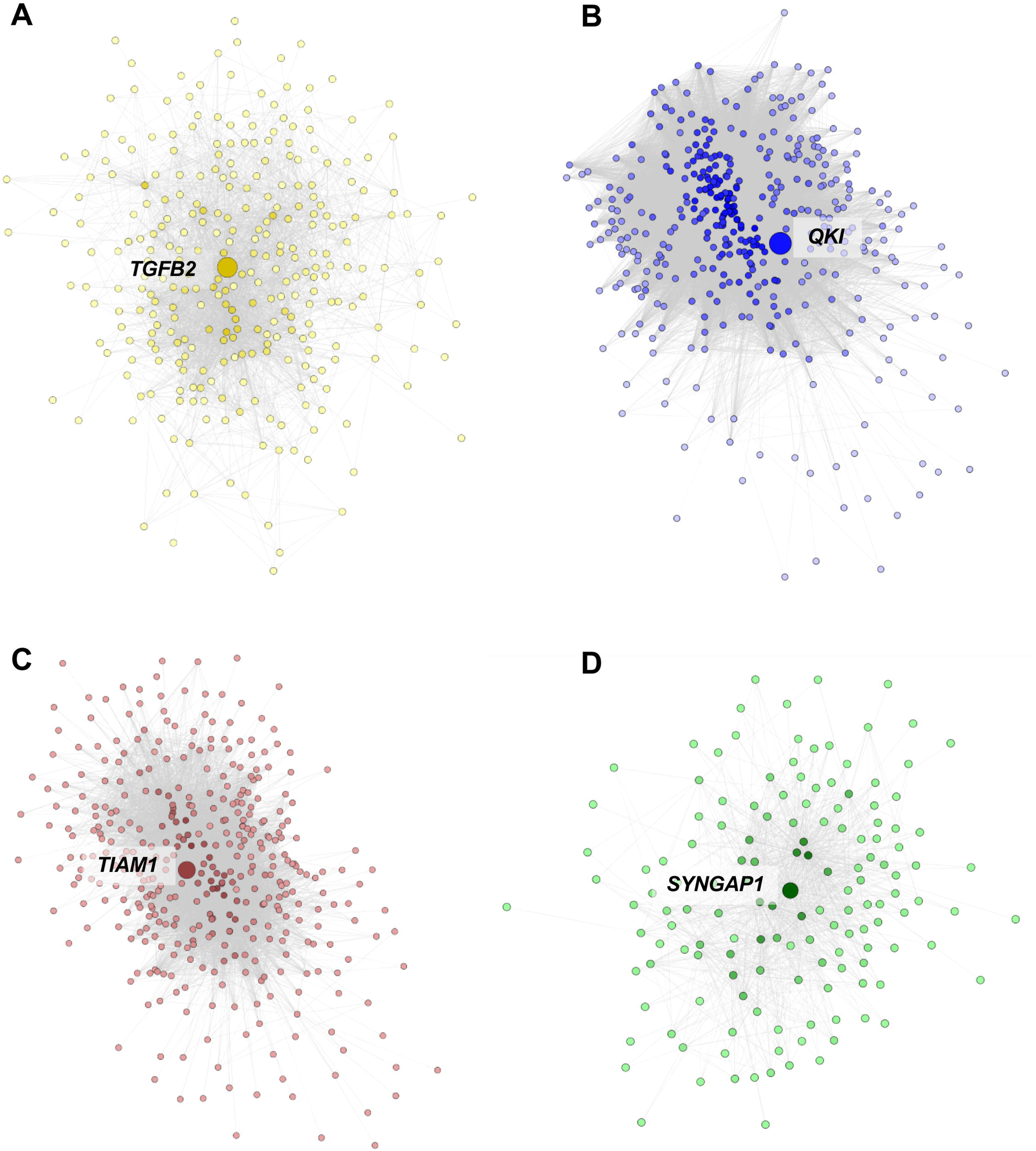
Coexpression network generated from the significant coexpression modules visualized with Cytoscape. The hub genes are the larger nodes. The color intensity of each node is proportional to the number of connections. (A) Network for the yellow module, upregulated in MSA-C. We represented edges with weight ≥ 0.05. (B) Newtork for the blue module, downregulated in MSA-C. We exported edges with weight ≥ 0.20. (C) Network for the brown module, upregulated in MSA-C. We exported edges with weight ≥ 0.20. (D) Network for the green module, upregulated in MSA-C. We exported edges with weight ≥ 0.01

We conducted GO enrichment analysis and observed the most significant and specific enrichment in the brown and blue modules (Fig. S20 and Table S14). The yellow module (upregulated) showed a heterogeneous enrichment, including immune response but also tissue and organ development and response to stress. The green module (upregulated) was enriched for membrane proteins, ribosome and translation. The brown module (upregulated) was enriched for synaptic functional classes (top class: FDR = 1.2E-33), and the blue module (downregulated) was enriched for myelination and oligodendrocyte classes (top class: FDR = 3.1E-09).

We explored the enrichment for specific brain cell types gene expression using the data from [63]. Accordingly with the GO enrichment we observed, we found a significant enrichment of astrocyte (adj *p* = 3.3E-19) and endothelial genes (adj *p* = 2.8E-04) in the yellow module (upregulated, enriched for immunity and organ development), and a significant enrichment of neuronal genes (adj *p* = 2.5E-60) in the brown module (upregulated, enriched for synaptic processes). Furthermore, we detected a significant enrichment of oligodendrocyte genes (adj *p* = 7.7E-33) in the blue module (downregulated, enriched for myelination) (Fig. S21).

We validated the results conducting module preservation analysis, using C2 as the test sample. We observed strong evidence of preservation for the blue (myelination) and brown (synapse) modules, and moderate evidence of preservation in the green module (translation). No evidence of preservation was detected for the yellow module (Fig. S22).

## Discussion

### Overview

We conducted a genome-wide expression profiling study using cerebellar white matter (CWM) homogenates and LCM purified oligodendrocytes from MSA patients and healthy controls (HC). Two independent cohorts were analyzed using different expression profiling approaches and the differentially expressed genes were prioritized using meta-analysis techniques. WGCNA was applied to find clusters of genes functionally related and associated with the disease. This is the largest RNA profiling study conducted on post-mortem brain samples from MSA patients to date.

### Differential dysregulation in MSA subtypes demonstrates more CWM transcriptional changes in MSA-C than in MSA-P

After p-value combination, we obtained the largest number of DEGs in the MSA-C subgroup comparison (*n* = 747). Only one gene was noted to be differentially expressed in the MSA-P sub-type analysis. Of note, the majority of the MSA-P patients had demonstrable GCIs in the CWM and 35 DEGs were identified when the MSA cohort was utilized as a single group in a case/control analysis (MSA-C plus MSA-P vs. HC). Of note, the ratio of MSA-P:MSA-C was 2.6:1 therefore the decreased number of DEGs noted in the combined MSA analysis is likely due to the higher number of MSA-P patients in our study. These results agree with the larger involvement of CWM alterations in MSA-C compared to MSA-P during the early stages of the disease [18, 46]. It is possible that due to the early involvement of CWM in MSA-C there is a longer time for the disease-related transcriptional changes to develop in the CWM in these patients. [46] found more cerebellar and pontine involvement in MSA-C compared to MSA-P. Dash *et al*. used voxel-based morphometry (VBM) and diffusion tensor imaging (DTI) to assess the WM and GM changes in the two MSA subtypes and healthy controls. In comparison to controls, MSA-C showed widespread WM changes in supratentorial and infratentorial regions, whereas MSA-P only showed the involvement of association tracts. Their comparison between MSA-C and MSA-P confirmed a greater prevalence of cerebellar changes in MSA-C patients.

### Oligodendrocyte genes are downregulated and enriched for myelination processes in MSA-C CWM

Results from multiple analyses in our study converge on strong evidence of the dysregulation of oligodendrocyte genes in MSA-C. WGCNA analysis showed the presence of a coexpression module (blue, *n* = 485 genes) negatively associated with disease status in MSA-C, enriched for myelination processes, and showing a very large prevalence of oligodendrocyte genes in comparison to the other modules. Additionally, this module showed a strong preservation in the independent C2 dataset. The top hub gene in this blue coexpression network was *QKI.* This gene (downregulated in MSA-C) encodes for an RNA-binding protein involved in myelination and oligodentrocyte differentiation [1]. Darbelli et al., (2017) conducted a transcriptomic analysis of oligodendrocyte-specific QKI conditional knock-out mouse brain and found approximately 1,800 genes differentially expressed and the underlying functional annotation of these genes were enriched for axon ensheathment and myelination. Moreover, they detected 810 alternatively spliced genes in the conditional knock-out animals. These results suggest a potential key role of *QKI* as a regulator of RNA metabolism and alternative splicing in oligodendrocytes.

Interestingly, key myelin genes, including *MBP*, *MAG*, *MOBP*, and *PLP1* were found significantly downregulated in MSA-C patients in our study. The study by Bettencourt et al., (2019) reported MSA-associated DNA methylation changes in *MOBP*, suggesting that the observed downregulation of this gene in MSA might be regulated by changes in DNA methylation levels.

As mentioned in the Introduction, the relocalization of p25α from the myelin sheath to the oligodendrocyte soma is one of the earliest molecular events that may trigger α-synuclein aggregation. This process may also slow oligodendrocyte precursor cell maturation by the α-synuclein mediated downregulation of myelin-gene regulatory factor and myelin basic protein [34]. Interestingly, the gene coding for p25α (*TPPP*), which is expressed in oligodendrocytes, was detected to be significantly downregulated in MSA-C patients in our study (adj *p* < 0.05). Additionally, *SNCA* was significantly downregulated after p-value combination in MSA-C (adj *p* < 0.05). The same result was found in another study [30], but not confirmed in other work [28, 42]. Other studies based on oligodendrocyte isolation and qPCA analysis described a basal expression and a trend of an increased expression in MSA patients [4, 19].

Other relevant genes from the LCM study also associated with the myelination process were *NF1*, *PLP1* and *ERMN*. *NF1* (*Neurofibromin 1*) was downregulated in MSA and it encodes for a protein specialized in the formation of myelin sheaths. Mutations in this gene causes Neurofibromatosis type 1, which is characterized by the growth of tumors along nerves in various parts of the body including the brain. *PLP1* (Proteolipid Protein 1) is specifically expressed in oligodendrocytes and it was downregulated in MSA-C patients in our sample. The protein product is a predominant component of myelin, and it also has a role in the maintenance of the myelin sheath as well as in oligodendrocyte development and axonal maintenance. Groh et al. [26] showed that mice with a loss of function *PLP1* mutation exhibit neuroinflammation that leads to axonal degeneration and neuronal cell loss. Finally, *ERMN* (Ermin), downregulated in MSA-C, is involved in myelinogenesis and in maintenance and stability of the myelin sheath.

It is worth mentioning other genes highly differentially expressed in the LCM study even if not directly functionally associated with myelination: *GGCX*, and *MOCS*. *GGCX* (Gamma-Glutamyl Carboxylase) was upregulated in MSA patients, and it is essential for activating vitamin K-dependent proteins [55]. Mutations in this gene cause the “GGCX Syndrome” (OMIM: 137167). It has been observed *in vitro* that Vitamin K delays α-synuclein fibrillization through its interaction at specific sites at the N-terminus of α-synuclein [52]. *MOCS1* (Molybdenum Cofactor Synthesis 1) was downregulated and it is involved in the biological activation of molybdenum. Mutations in *MOCS1* causes molybdenum cofactor deficiency which is characterized by neurodegeneration and seizures [5].

### Neuron cell-specific genes are upregulated in MSA CWM and are enriched for biological pathways related to synaptic processes

Two different analytical approaches suggested significant upregulation of neuronal cell-specific genes in MSA-C and these genes were enriched for biological roles in synaptic and neurogenesis processes. When we classified the DEGs from MSA-C according to our cell-specific gene analysis approach, we detected an upregulation of neuronal genes and an enrichment for synaptic and neuronal processes. Using WGCNA analysis we detected a module of 451 co-expressed genes (brown) significantly upregulated in MSA tissue and enriched for synaptic processes. The genes in this “brown” module demonstrated a higher prevalence of neuronal-specific genes in comparison to the other significant modules. As was the case with the blue module (discussed above), the brown module showed strong model preservation in our independent C2 dataset. The hub gene in the brown module co-expression network was *TIAM1* (T Cell Lymphoma Invasion And Metastasis 1). This gene (upregulated in MSA-C CWM) encodes a RAC1-specific guanine nucleotide exchange factor that is involved in the control of excitatory synapse development [57]. Interestingly, the green module (significantly upregulated in MSA-C CWM) was correlated with the brown module and showed an enrichment in protein transport and translation. The hub gene in this module was *SYNGAP1* (Synaptic Ras GTPase Activating Protein 1, upregulated in MSA-C) which, like TIAM1, is also involved in synaptogenesis [7, 15]. The upregulation of neuron-specific genes and the enrichment for synaptogenesis is surprising in the context of a neurodegenerative disease like MSA. Monomeric α-synuclein is normally located in the presynaptic nerve terminals and is involved in synaptogenesis [64, 65]. Perhaps, the enrichment of the synaptogenesis process in MSA-C CWM in our study might be a consequence of an abnormal accumulation of α-synuclein in the synapse of MSA patients. This elevated synaptic accumulation was previously described to precede the re-localization of α-synuclein from neurons to oligodendrocytes and may represent one of the earliest and ongoing molecular events associated with the disease [54]. Alternatively, this upregulation of synaptogenesis in the context of neurodegeneration in the MSA-C brain may represent a transcriptional attempt by the remaining neurons to compensate for the overall synaptic losses within the CWM.

### The importance of neuroinflammation in MSA-C

The combined relocalization of p25α and the ectopic presence of α-synuclein in oligodendrocytes are thought to trigger the formation of α-synuclein and p25α inclusions. These inclusions and resulting oligodendrocyte dysfunction, activate microglia and astrocytes contributing to the neurodegenerative process through neuroinflammation (Fanciulli and Wenning, 2015). These phenomena may explain our finding of the upregulated yellow module (314 genes). This module includes a large prevalence of astrocyte and microglia genes compared to the other significant modules and it is enriched for inflammatory and tissue/organ developmental processes. We found that astrocyte and endothelial specific genes were significantly upregulated in the DEGs from bulk tissue. The top hub gene in the yellow module was *TGFB2* (Transforming Growth Factor Beta 2). This gene encodes a secreted ligand of the TGF-beta (transforming growth factor-beta) superfamily of proteins that are involved in the recruitment and activation of SMAD family transcription factors. Interestingly, the levels of TGFβ-2 were previously found to be increased in the neocortex of AD and dementia with Lewy bodies and were positively correlated with neuropathological markers of disease severity [14]. This finding may suggest that TGF-beta is a key regulator of the inflammatory processes that may be more generalizable to neurodegenerative diseases regardless of the underlying causes and resulting neuropathologies. In the yellow module we found also *MASP1 (log2 FC = 0.944; adj p 0.380),* whose mRNA expression was found upregulated in a separate study conducted using frontal lobe post-mortem brains from MSA patients and controls [29].

### Collagen genes are upregulated in MSA

In the combined MSA group after p-value combination we detected 35 genes, most of them upregulated in patients. In both enrichment analyses we detected a key role of collagen genes: *COL4A1*, *COL4A2*, and *ITGA11;* all upregulated. *COL4A1* (collagen type IV alpha 1 chain) and *COL4A2* (collagen type IV alpha 1 chain) encode respectively for the alpha 1 and alpha 2 chains of type IV collagen which are important components of the basement membrane in all tissues, especially blood vessels. *ITGA11* (Integrin Subunit Alpha 11) is functionally related as it is a collagen receptor. Mutations in *COL4A1* and *COL4A2* have been associated with sporadic brain small vessel disease [44] and porencephaly [10]. Recently Paiva et al., (2018) found *COL4A2* upregulated in both A30P aSyn mice and dopaminergic neurons expressing A30P aSyn, suggesting a key role of collagen-related genes in α-synuclein induced toxicity. In the same study, they demonstrated a regulation of *COL4A2* expression by miR-29a-3p, known to target *COL4A4* mRNA. In a separate study the loss of miR-29a was correlated with increased levels of BACE1 and amyloid-β in sporadic Alzheimer’s Disease [27]. Finally, lack of collagen VI has been related to neurodegeneration in mice models [11], and its presence has been related to a neuroprotective role against β-Amyloid toxicity [13].

Beside the collagen related pathway, the top genes detected in the differential expression analysis were: *ACTN1* (Actinin Alpha 1), *EMP1* (Epithelial Membrane Protein 1), and *NFIL3* (Nuclear Factor, Interleukin 3 Regulated). Expression changes of *ACTN1* were associated with AD in hippocampus [24], whereas *NFIL3* was associated with neuroprotection in models of Amyotrophic Lateral Sclerosis [56]. EMP1 protein was also found upregulated in 5xFAD AD model [21].

### MSA-C shows a common transcriptional background with Alzheimer’s Disease

We detected a large functional network in MSA-C patients that included *APP* and other AD-related genes that included *PSEN1*, *CLU*, *ROCK2, EFNA1* and *DYRK1*. The module (M1) including these genes was enriched for amyloid-β metabolism. The strongest enrichment between MSA-C and AD DEGs was found in the temporal cortex and parahippocampal gyrus (AMP-AD data).

### Study Limitations

We note some limitations of our study. First, MSA is a rare disease and although our cohort is the largest that has been expression profiled to date it is still likely that we are underpowered to detect small effect sizes that could be functionally important. Secondly, we acknowledge that the findings would be improved by the inclusion of additional brain regions that may be altered by the disease. For example, it isn’t particularly surprising that we noted the most significant cerebellar transcriptional changes in MSA-C, a clinical subtype of MSA with predominating cerebellar symptoms. It would be interesting to compare the transcriptomic changes in the striatum, olivary nuclei, and pontine nuclei as well. Thirdly, we assessed C1 and C2 using different profiling approaches. This could be considered a positive aspect of our work as the identified transcriptional changes that cross-validate are likely not specific to a particular gene expression measurement approach and therefore may have higher reproducibility, however, this could also be considered a limitation as some true associations may be unreported due to their failure in one of the profiling chemistries and not due to the underlying biology. Lastly, layering additional genomic information – like DNA sequence information – would also enhance the study as it could facilitate more detailed analyses such as allele specific expression or epigenetic regulation of transcription.

### Conclusions

This is the largest study ever conducted on the MSA brain transcriptome. We utilized two different cohorts that were each assessed by different gene expression analysis chemistries that we propose increases the robustness of DEG and co-expression network detection.

The main findings of this study are the multiple evidence of oligodendrocyte gene downregulation associated with the loss of myelination. We detected the *QKI* gene as a master regulator of this associated gene network. Additionally, we showed an upregulation of neuronal-specific gene expression possibly as a consequence of the initial accumulation of monomeric α-synuclein in neurons, with *TIAM1* and *SYNGAP1* as top hubs in the two networks. An additional coexpression network highlighted the later stages of the neurodegenerative cascade with activation of microglia and astrocytes. Finally, our results suggest a common transcriptional background between MSA and AD, potentially through *APP*-mediated mechanisms.

## Supporting information

Supplementary Materials an Results

## Acknowledgements

The authors acknowledge funding support Austrian Science Fund (FWF). The MSA laser capture microdissection work was supported by NIH-NINDS grant R21-NS093222 to Matthew Huentelman. All of the Alzheimer’s Disease RNA sequencing data used for this study were downloaded from the AMP-AD portal (Accession number: Syn14237651).

Mayo Study: Study data were provided by the following sources: The Mayo Clinic Alzheimers Disease Genetic Studies, led by Dr. Nilufer Taner and Dr. Steven G. Younkin, Mayo Clinic, Jacksonville, FL using samples from the Mayo Clinic Study of Aging, the Mayo Clinic Alzheimers Disease Research Center, and the Mayo Clinic Brain Bank. Data collection was supported through funding by NIA grants P50 AG016574, R01 AG032990, U01 AG046139, R01 AG018023, U01 AG006576, U01 AG006786, R01 AG025711, R01 AG017216, R01 AG003949, NINDS grant R01 NS080820, CurePSP Foundation, and support from Mayo Foundation. Study data includes samples collected through the Sun Health Research Institute Brain and Body Donation Program of Sun City, Arizona, and the New South Wales (NSW) Brain Bank (Sydney, AU). The Brain and Body Donation Program is supported by the National Institute of Neurological Disorders and Stroke (U24 NS072026 National Brain and Tissue Resource for Parkinsons Disease and Related Disorders), the National Institute on Aging (P30 AG19610 Arizona Alzheimers Disease Core Center), the Arizona Department of Health Services (contract 211002, Arizona Alzheimers Research Center), the Arizona Biomedical Research Commission (contracts 4001, 0011, 05-901 and 1001 to the Arizona Parkinson’s Disease Consortium) and the Michael J. Fox Foundation for Parkinsons Research.

ROSMAP study: Gene Expression data from this study was funded by grant U01AG046152.

Mount Sinai Study: Gene Expression data for this study was funded through grant U01AG046170.

## REFERENCES

1. Aberg K, Saetre P, Jareborg N, Jazin E (2006) Human QKI, a potential regulator of mRNA expression of human oligodendrocyte-related genes involved in schizophrenia. Proc Natl Acad Sci. doi: 10.1073/pnas.0601213103

2. Al-Chalabi A, Dürr A, Wood NW, Parkinson MH, Camuzat A, Hulot J-S, Morrison KE, Renton A, Sussmuth SD, Landwehrmeyer BG, Ludolph A, Agid Y, Brice A, Leigh PN, Bensimon G (2009) Genetic variants of the alpha-synuclein gene SNCA are associated with multiple system atrophy. PLoS One 4:e7114. doi: 10.1371/journal.pone.0007114

3. Allen M, Carrasquillo MM, Funk C, Heavner BD, Zou F, Younkin CS, Burgess JD, Chai HS, Crook J, Eddy JA, Li H, Logsdon B, Peters MA, Dang KK, Wang X, Serie D, Wang C, Nguyen T, Lincoln S, Malphrus K, Bisceglio G, Li M, Golde TE, Mangravite LM, Asmann Y, Price ND, Petersen RC, Graff-Radford NR, Dickson DW, Younkin SG, Ertekin-Taner N (2016) Human whole genome genotype and transcriptome data for Alzheimer’s and other neurodegenerative diseases. Sci Data 3. doi: 10.1038/sdata.2016.89

4. Asi YT, Simpson JE, Heath PR, Wharton SB, Lees AJ, Revesz T, Houlden H, Holton JL (2014) Alpha-synuclein mRNA expression in oligodendrocytes in MSA. Glia 62:964–970. doi: 10.1002/glia.22653

5. Atwal PS, Scaglia F (2016) Molybdenum cofactor deficiency. Mol. Genet. Metab. 117:1–4

6. Bennett DA, Schneider JA, Arvanitakis Z, Wilson RS (2012) Overview and findings from the religious orders study. Curr Alzheimer Res 9:628–645. doi: 10.2174/156720512801322573

7. Berryer MH, Chattopadhyaya B, Xing P, Riebe I, Bosoi C, Sanon N, Antoine-Bertrand J, Lévesque M, Avoli M, Hamdan FF, Carmant L, Lamarche-Vane N, Lacaille JC, Michaud JL, Di Cristo G (2016) Decrease of SYNGAP1 in GABAergic cells impairs inhibitory synapse connectivity, synaptic inhibition and cognitive function. Nat Commun. doi: 10.1038/ncomms13340

8. Bettencourt C, Foti SC, Miki Y, Botia J, Chatterjee A, Warner TT, Revesz T, Lashley T, Balazs R, Viré E, Holton JL (2019) White matter DNA methylation profiling reveals deregulation of HIP1, LMAN2, MOBP, and other loci in multiple system atrophy. Acta Neuropathol. doi: 10.1007/s00401-019-02074-0

9. Bhidayasiri R, Ling H (2008) Multiple System Atrophy. Neurologist 14:224–237. doi: 10.1097/NRL.0b013e318167b93f

10. Breedveld G, De Coo IF, Lequin MH, Arts WFM, Heutink P, Gould DB, John SWM, Oostra B, Mancini GMS (2006) Novel mutations in three families confirm a major role of COL4A1 in hereditary porencephaly. J Med Genet 43:490–495. doi: 10.1136/jmg.2005.035584

11. Cescon M, Chen P, Castagnaro S, Gregorio I, Bonaldo P (2016) Lack of collagen VI promotes neurodegeneration by impairing autophagy and inducing apoptosis during aging. Aging (Albany NY) 8:1083–1101. doi: 10.18632/aging.100924

12. Chen BJ, Mills JD, Takenaka K, Bliim N, Halliday GM, Janitz M (2016) Characterization of circular RNAs landscape in multiple system atrophy brain. J Neurochem 139:485–496. doi: 10.1111/jnc.13752

13. Cheng JS, Dubal DB, Kim DH, Legleiter J, Cheng IH, Yu G-Q, Tesseur I, Wyss-Coray T, Bonaldo P, Mucke L (2009) Collagen VI protects neurons against Abeta toxicity. Nat Neurosci 12:119–21. doi: 10.1038/nn.2240

14. Chong JR, Chai YL, Lee JH, Howlett D, Attems J, Ballard CG, Aarsland D, Francis PT, Chen CP, Lai MKP (2017) Increased transforming growth factor β in the neocortex of Alzheimer’s disease and dementia with lewy bodies is correlated with disease severity and soluble Aβ 42 load. J Alzheimer’s Dis. doi: 10.3233/JAD-160781

15. Clement JP, Ozkan ED, Aceti M, Miller CA, Rumbaugh G (2013) SYNGAP1 Links the Maturation Rate of Excitatory Synapses to the Duration of Critical-Period Synaptic Plasticity. J Neurosci. doi: 10.1523/jneurosci.0765-13.2013

16. Curry-Hyde A, Chen BJ, Ueberham U, Arendt T, Janitz M (2017) Multiple System Atrophy: Many Lessons from the Transcriptome. Neurosci 107385841772391. doi: 10.1177/1073858417723915

17. Darbelli L, Choquet K, Richard S, Kleinman CL (2017) Transcriptome profiling of mouse brains with qkI-deficient oligodendrocytes reveals major alternative splicing defects including self-splicing. Sci Rep. doi: 10.1038/s41598-017-06211-1

18. Dash SK, Stezin A, Takalkar T, George L, Kamble NL, Netravathi M, Yadav R, Kumar KJ, Ingalhalikar M, Saini J, Pal PK (2018) Abnormalities of white and grey matter in early multiple system atrophy: comparison of parkinsonian and cerebellar variants. Eur. Radiol. 1–9

19. Djelloul M, Holmqvist S, Boza-Serrano A, Azevedo C, Yeung MS, Goldwurm S, Frisén J, Deierborg T, Roybon L (2015) Alpha-Synuclein Expression in the Oligodendrocyte Lineage: An in Vitro and in Vivo Study Using Rodent and Human Models. Stem Cell Reports. doi: 10.1016/j.stemcr.2015.07.002

20. Dobin A, Davis CA, Schlesinger F, Drenkow J, Zaleski C, Jha S, Batut P, Chaisson M, Gingeras TR (2013) STAR: Ultrafast universal RNA-seq aligner. Bioinformatics 29:15–21. doi: 10.1093/bioinformatics/bts635

21. Duran RCD, Wang CY, Zheng H, Deneen B, Wu JQ (2019) Brain region-specific gene signatures revealed by distinct astrocyte subpopulations unveil links to glioma and neurodegenerative diseases. eNeuro. doi: 10.1523/ENEURO.0288-18.2019

22. Ewels P, Magnusson M, Lundin S, K??ller M (2016) MultiQC: Summarize analysis results for multiple tools and samples in a single report. Bioinformatics 32:3047–3048. doi: 10.1093/bioinformatics/btw354

23. Goedert M (2001) Alpha-synuclein and neurodegenerative diseases. Nat. Rev. Neurosci. 2:492–501

24. Gómez Ravetti M, Rosso OA, Berretta R, Moscato P (2010) Uncovering molecular biomarkers that correlate cognitive decline with the changes of hippocampus’ gene expression profiles in Alzheimer’s disease. PLoS One 5. doi: 10.1371/journal.pone.0010153

25. Greene CS, Krishnan A, Wong AK, Ricciotti E, Zelaya RA, Himmelstein DS, Zhang R, Hartmann BM, Zaslavsky E, Sealfon SC, Chasman DI, Fitzgerald GA, Dolinski K, Grosser T, Troyanskaya OG (2015) Understanding multicellular function and disease with human tissue-specific networks. Nat Genet. doi: 10.1038/ng.3259

26. Groh J, Friedman HC, Orel N, Ip CW, Fischer S, Spahn I, Schäffner E, Hörner M, Stadler D, Buttmann M, Varallyay C, Solymosi L, Sendtner M, Peterson AC, Martini R (2016) Pathogenic inflammation in the CNS of mice carrying human PLP1 mutations. Hum Mol Genet 25:4686–4702. doi: 10.1093/hmg/ddw296

27. Hebert SS, Horre K, Nicolai L, Papadopoulou AS, Mandemakers W, Silahtaroglu AN, Kauppinen S, Delacourte A, De Strooper B (2008) Loss of microRNA cluster miR-29a/b-1 in sporadic Alzheimer’s disease correlates with increased BACE1/ - secretase expression. Proc Natl Acad Sci 105:6415–6420. doi: 10.1073/pnas.0710263105

28. Jin H, Ishikawa K, Tsunemi T, Ishiguro T, Amino T, Mizusawa H (2008) Analyses of copy number and mRNA expression level of the α-synuclein gene in multiple system atrophy. J Med Dent Sci. doi: 10.11480/jmds.550117

29. Kiely AP, Murray CE, Foti SC, Benson BC, Courtney R, Strand C, Lashley T, Holton JL (2018) Immunohistochemical and molecular investigations show alteration in the inflammatory profile of multiple system atrophy brain. J Neuropathol Exp Neurol. doi: 10.1093/jnen/nly035

30. Langerveld AJ, Mihalko D, DeLong C, Walburn J, Ide CF (2007) Gene expression changes in postmortem tissue from the rostral pons of multiple system atrophy patients. Mov Disord. doi: 10.1002/mds.21259

31. Langfelder P, Horvath S (2008) WGCNA: an R package for weighted correlation network analysis. BMC Bioinformatics 9:559. doi: 10.1186/1471-2105-9-559

32. Liao Y, Smyth GK, Shi W (2014) FeatureCounts: An efficient general purpose program for assigning sequence reads to genomic features. Bioinformatics 30:923–930. doi: 10.1093/bioinformatics/btt656

33. Love MI, Huber W, Anders S (2014) Moderated estimation of fold change and dispersion for RNA-seq data with DESeq2. Genome Biol 15:550. doi: 10.1186/s13059-014-0550-8

34. May VEL, Ettle B, Poehler AM, Nuber S, Ubhi K, Rockenstein E, Winner B, Wegner M, Masliah E, Winkler J (2014) α-Synuclein impairs oligodendrocyte progenitor maturation in multiple system atrophy. Neurobiol Aging 35:2357–2368. doi: 10.1016/j.neurobiolaging.2014.02.028

35. Mills JD, Kim WS, Halliday GM, Janitz M (2015) Transcriptome analysis of grey and white matter cortical tissue in multiple system atrophy. Neurogenetics 16:107–122. doi: 10.1007/s10048-014-0430-0

36. Mills JD, Ward M, Kim WS, Halliday GM, Janitz M (2016) Strand-specific RNA-sequencing analysis of multiple system atrophy brain transcriptome. Neuroscience 322:234–250. doi: 10.1016/j.neuroscience.2016.02.042

37. Mitsui J, Matsukawa T, Ishiura H, Fukuda Y, Ichikawa Y, Date H, Ahsan B, Nakahara Y, Momose Y, Takahashi Y, Iwata A, Goto J, Yamamoto Y, Komata M, Shirahige K, Hara K, Kakita A, Yamada M, Takahashi H, Onodera O, Nishizawa M, Takashima H, Kuwano R, Watanabe H, Ito M, Sobue G, Soma H, Yabe I, Sasaki H, Aoki M, Ishikawa K, Mizusawa H, Kanai K, Hattori T, Kuwabara S, Arai K, Koyano S, Kuroiwa Y, Hasegawa K, Yuasa T, Yasui K, Nakashima K, Ito H, Izumi Y, Kaji R, Kato T, Kusunoki S, Osaki Y, Horiuchi M, Kondo T, Murayama S, Hattori N, Yamamoto M, Murata M, Satake W, Toda T, Dürr A, Brice A, Filla A, Klockgether T, Wallner U, Nicholson G, Gilman S, Shults CW, Tanner CM, Kukull WA, Lee VMY, Masliah E, Low PA, Sandroni P, Trojanowski JQ, Ozelius L, Foroud T, Tsuji S (2013) Mutations in COQ2 in familial and sporadic multiple-system atrophy the multiple-system atrophy research collaboration. N Engl J Med. doi: 10.1056/NEJMoa1212115

38. Nirenberg MJ, Libien J, Vonsattel J-P, Fahn S (2007) Multiple system atrophy in a patient with the spinocerebellar ataxia 3 gene mutation. Mov Disord 22:251–254. doi: 10.1002/mds.21231

39. Ordway GA, Szebeni A, Duffourc MM, Dessus-Babus S, Szebeni K (2009) Gene expression analyses of neurons, astrocytes, and oligodendrocytes isolated by laser capture microdissection from human brain: Detrimental effects of laboratory humidity. J Neurosci Res 87:2430–2438. doi: 10.1002/jnr.22078

40. Paiva I, Jain G, Lazaro DF, Jercic KG, Hentrich T, Kerimoglu C, Pinho R, Szego EM, Burkhardt S, Capece V, Halder R, Islam R, Xylaki M, Caldi Gomes LA, Roser A-E, Lingor P, Schulze-Hentrich JM, Borovecki F, Fischer A, Outeiro TF (2018) Alpha-synuclein deregulates the expression of COL4A2 and impairs ER-Golgi function. Neurobiol Dis 119:121–135. doi: 10.1016/j.nbd.2018.08.001

41. Papp MI, Kahn JE, Lantos PL (1989) Glial cytoplasmic inclusions in the CNS of patients with multiple system atrophy (striatonigral degeneration, olivopontocerebellar atrophy and Shy-Drager syndrome). J. Neurol. Sci. 94:79–100

42. Piper DA, Sastre D, Schüle B (2018) Advancing stem cell models of alpha-synuclein gene regulation in neurodegenerative disease. Front. Neurosci.

43. Quinn N, Wenning G (1995) Multiple system atrophy. Curr Opin Neurol 8:323–326

44. Rannikmäe K, Davies G, Thomson PA, Bevan S, Devan WJ, Falcone GJ, Traylor M, Anderson CD, Battey TWK, Radmanesh F, Deka R, Woo JG, Martin LJ, Jimenez-Conde J, Selim M, Brown DL, Silliman SL, Kidwell CS, Montaner J, Langefeld CD, Slowik A, Hansen BM, Lindgren AG, Meschia JF, Fornage M, Bis JC, Debette S, Ikram MA, Longstreth WT, Schmidt R, Zhang CR, Yang Q, Sharma P, Kittner SJ, Mitchell BD, Holliday EG, Levi CR, Attia J, Rothwell PM, Poole DL, Boncoraglio GB, Psaty BM, Malik R, Rost N, Worrall BB, Dichgans M, Van Agtmael T, Woo D, Markus HS, Seshadri S, Rosand J, Sudlow CLM (2015) Common variation in COL4A1/COL4A2 is associated with sporadic cerebral small vessel disease. Neurology 84:918–926. doi: 10.1212/WNL.0000000000001309

45. Ritchie ME, Phipson B, Wu D, Hu Y, Law CW, Shi W, Smyth GK (2015) limma powers differential expression analyses for RNA-sequencing and microarray studies. Nucleic Acids Res 43:e47. doi: 10.1093/nar/gkv007

46. Roncevic D, Palma JA, Martinez J, Goulding N, Norcliffe-Kaufmann L, Kaufmann H (2014) Cerebellar and parkinsonian phenotypes in multiple system atrophy: Similarities, differences and survival. J Neural Transm 121:507–512. doi: 10.1007/s00702-013-1133-7

47. Sailer A, Scholz SW, Nalls MA, Schulte C, Federoff M, Price TR, Lees A, Ross OA, Dickson DW, Mok K, Mencacci NE, Schottlaender L, Chelban V, Ling H, O’Sullivan SS, Wood NW, Traynor BJ, Ferrucci L, Federoff HJ, Mhyre TR, Morris HR, Deuschl G, Quinn N, Widner H, Albanese A, Infante J, Bhatia KP, Poewe W, Oertel W, Höglinger GU, Wüllner U, Goldwurm S, Pellecchia MT, Ferreira J, Tolosa E, Bloem BR, Rascol O, Meissner WG, Hardy JA, Revesz T, Holton JL, Gasser T, Wenning GK, Singleton AB, Houlden H (2016) A genome-wide association study in multiple system atrophy. Neurology 87:1591–1598. doi: 10.1212/WNL.0000000000003221

48. Scholz SW, Houlden H, Schulte C, Sharma M, Li A, Berg D, Melchers A, Paudel R, Gibbs JR, Simon-Sanchez J, Paisan-Ruiz C, Bras J, Ding J, Chen H, Traynor BJ, Arepalli S, Zonozi RR, Revesz T, Holton J, Wood N, Lees A, Oertel W, Wüllner U, Goldwurm S, Pellecchia MT, Illig T, Riess O, Fernandez HH, Rodriguez RL, Okun MS, Poewe W, Wenning GK, Hardy JA, Singleton AB, Gasser T (2009) SNCA variants are associated with increased risk for multiple system atrophy. Ann Neurol 65:610–614. doi: 10.1002/ana.21685

49. Schottlaender L V., Houlden H (2014) Mutant COQ2 in Multiple-System Atrophy [5]. N. Engl. J. Med.

50. Schroder MS, Culhane AC, Quackenbush J, Haibe-Kains B (2011) survcomp: an R/Bioconductor package for performance assessment and comparison of survival models. Bioinformatics 27:3206–3208. doi: 10.1093/bioinformatics/btr511

51. Shannon P, Markiel A, Ozier O, Baliga NS, Wang JT, Ramage D, Amin N, Schwikowski B, Ideker T (2003) Cytoscape: A software Environment for integrated models of biomolecular interaction networks. Genome Res. doi: 10.1101/gr.1239303

52. Da Silva FL, Coelho Cerqueira E, De Freitas MS, Gonçalves DL, Costa LT, Follmer C (2013) Vitamins K interact with N-terminus α-synuclein and modulate the protein fibrillization in vitro. Exploring the interaction between quinones and α-synuclein. Neurochem Int 62:103–112. doi: 10.1016/j.neuint.2012.10.001

53. Stefanova N, Bücke P, Duerr S, Wenning GK (2009) Multiple system atrophy: an update. Lancet Neurol. 8:1172–1178

54. Stefanova N, Wenning GK (2016) Multiple system atrophy: Emerging targets for interventional therapies. Neuropathol. Appl. Neurobiol.

55. Suleiman L, Négrier C, Boukerche H (2013) Protein S: A multifunctional anticoagulant vitamin K-dependent protein at the crossroads of coagulation, inflammation, angiogenesis, and cancer. Crit. Rev. Oncol. Hematol. 88:637–654

56. Tamai S, Imaizumi K, Kurabayashi N, Nguyen MD, Abe T, Inoue M, Fukada Y, Sanada K (2014) Neuroprotective role of the basic leucine zipper transcription factor NFIL3 in models of amyotrophic lateral sclerosis. J Biol Chem 289:1629–38. doi: 10.1074/jbc.M113.524389

57. Um K, Niu S, Duman JG, Cheng JX, Tu YK, Schwechter B, Liu F, Hiles L, Narayanan AS, Ash RT, Mulherkar S, Alpadi K, Smirnakis SM, Tolias KF (2014) Dynamic Control of Excitatory Synapse Development by a Rac1 GEF/GAP Regulatory Complex. Dev Cell. doi: 10.1016/j.devcel.2014.05.011

58. Vanacore N (2005) Epidemiological evidence on multiple system atrophy. J Neural Transm 112:1605–12. doi: 10.1007/s00702-005-0380-7

59. Wakabayashi K, Yoshimoto M, Tsuji S, Takahashi H (1998) α-synuclein immunoreactivity in glial cytoplasmic inclusions in multiple system atrophy. Neurosci Lett 249:180–182. doi: 10.1016/S0304-3940(98)00407-8

60. Wang J, Duncan D, Shi Z, Zhang B (2013) WEB-based GEne SeT AnaLysis Toolkit (WebGestalt): update 2013. Nucleic Acids Res 41. doi: 10.1093/nar/gkt439

61. Wang M, Beckmann ND, Roussos P, Wang E, Zhou X, Wang Q, Ming C, Neff R, Ma W, Fullard JF, Hauberg ME, Bendl J, Peters MA, Logsdon B, Wang P, Mahajan M, Mangravite LM, Dammer EB, Duong DM, Lah JJ, Seyfried NT, Levey AI, Buxbaum JD, Ehrlich M, Gandy S, Katsel P, Haroutunian V, Schadt E, Zhang B (2018) The Mount Sinai cohort of large-scale genomic, transcriptomic and proteomic data in Alzheimer’s disease. Sci data 5:180185. doi: 10.1038/sdata.2018.185

62. Zaykin D V. (2011) Optimally weighted Z-test is a powerful method for combining probabilities in meta-analysis. J Evol Biol 24:1836–1841. doi: 10.1111/j.1420-9101.2011.02297.x

63. Zhang Y, Chen K, Sloan SA, Bennett ML, Scholze AR, O’Keeffe S, Phatnani HP, Guarnieri P, Caneda C, Ruderisch N, Deng S, Liddelow SA, Zhang C, Daneman R, Maniatis T, Barres BA, Wu JQ (2014) An RNA-Sequencing Transcriptome and Splicing Database of Glia, Neurons, and Vascular Cells of the Cerebral Cortex. J Neurosci 34:11929–11947. doi: 10.1523/JNEUROSCI.1860-14.2014

64. Zhong H, May MJ, Jimi E, Ghosh S (2002) The phosphorylation status of nuclear NF-κB determines its association with CBP/p300 or HDAC-1. Mol Cell 9:625–636. doi: 10.1016/S1097-2765(02)00477-X

65. Zhong SC, Luo X, Chen XS, Cai QY, Liu J, Chen XH, Yao ZX (2010) Expression and subcellular location of alpha-synuclein during mouse-embryonic development. Cell Mol Neurobiol. doi: 10.1007/s10571-009-9473-4

