## Supplementary Materials an Results for "Transcriptional profiling of Multiple System Atrophy cerebellar tissue highlights differences between the parkinsonian and cerebellar sub-types of the disease"

**Supplementary Appendix**

1. **Supplementary Methods**
2. **Supplementary Results**

**1. Supplementary Methods**

**Human samples description**

We analyzed two independent cohorts of post mortem brains (cerebellum white matter). The cohort 1 (C1) was obtained from both the New South Wales (NSW) brain brank (Sydney, AU; *n* = 20) and from the Brain and Body Donation Program (BBPD) (Sun City, AZ; *n* = 18), including 19 MSA pathologically proven cases and 19 Healthy Controls (HC). The number of samples for each brain bank was matched between MSA and HC (NSW = 10; BBPD = 9 in each group). All the sample characteristics and the comparison between groups are reported in ***Table 1A***. The samples were matched for age (10 males and 9 females in each group), and there were not significant differences for postmortem interval PMI (*t* = -0.311; *p* = 0.757) and age (*W* = 175, *p* = 0.884). Among the patients, five were clinically diagnosed as MSA-P and other five were clinically diagnosed as MSA-C. The remaining patients were not attributed to a specific clinical subtype. The MSA-P and MSA-C subgroups compared with HC did not show any significant difference in age, sex and PMI distribution. The C2 was obtained from the Queen Square Brain Bank for Neurological Disorders, (London, UK), including 48 patients and 47 HC. All the sample characteristics and the comparisons between groups are reported in ***Table 1B***. There were not significant difference in the gender distribution (*p* = 0.402) and PMI (*W* = 1172; *p* = 0.746), but the age in HC was significantly higher than in MSA cases (*W* = 159.5; *p* = 5.6E-13). Among the patients, 37 were MSA-P, whereas the remaining (*n* = 11) were clinically diagnosed as MSA-C. Also the subgroups compared with HC showed a significant difference for age, but not for PMI and gender distributions. The same patients donors from C2 were classified by neuropathological examination for: striato-nigral degeneration (SND; n = 16), olivo ponto-cerebellar atrophy (OPCA; n = 15) and mixed (SND/OPCA; n = 17). Finally, a subsample of the C1 was selected to conduct Laser Capture Microdissection (LCM) on Oligodendrocytes (6 MSA and 6 HC). There were not significant differences in age (*t* = -0.469; *p* = 0.650), PMI (*W* =26.5; *p* = 0.199) or biological sex (*p* = 0.547). Three MSA cases were diagnosed as MSA-C, one as MSA-P, and the remaining could not be diagnosed to the subtypes MSA-P or MSA-C (***Table 1C***).

**RNA extraction and RNA sequencing**

RNA extraction and sequencing were conducted at the Translational Genomics Research Institute (Phoenix, AZ). For the C1, total RNA was extracted using the Qiagen miRNAeasy kit (#217004), and DNAse-treated (Qiagen). Quality was assessed by Bioanalyzer (Agilent). Sequencing libraries were prepared with 250 ng of total RNA using Illumina’s Truseq RNA Sample Preparation Kit v2 (Illumina, Inc) following the manufacturer’s protocol. In brief, poly-A containing mRNA molecules were purified using poly-T oligo attached magnetic beads. The mRNA was then thermally fragmented and converted to double-stranded cDNA. The cDNA fragments were end-repaired, a single “A” nucleotide was incorporated, sequencing adapters were ligated, and fragments were enriched with 15 cycles of PCR. Final PCR-enriched fragments were validated on a 2200 TapeStation (Agilent Technologies) and quantitated via qPCR using Kapa’s Library Quantification Kit (Kapa Biosystems) on the QuantStudio 6 Flex Instrument (ThermoFisher). The final library was sequenced by 50 bp paired-end sequencing on a HiSeq 2500.

For the C2 as well, RNA extraction and sequencing were conducted at the Translational Genomics Research Institute (Phoenix, AZ). Total DNAse-treated RNA was extracted in TRI Reagent (Zymo, #R2050) from ~ 5 mg of tissue using Rino Tubes (Next Advance , #13503) with one 0.1 mL scoop of Stainless Steel Beads 0.9 – 2.0 mm blend RNase Free (Next Advance, #SSB14B), Bullet Blender Gold (Next Advance, model BB24-AU), and Zymo Direct-zol-96 RNA (#R2054) utilizing the kit’s DNase I digestion step. Extracted RNA was then assayed on an Agilent TapeStation 4200 and Agilent RNA ScreenTape (#5067-5576). The mean RNA integrity number equivalent (RINe) and concentration for MSA cases were 3.42 and 77.6ng/μl, and Control cases were 3.59 and 64.76 ng/μl, respectively. Following TempO-Seq Assay User Guide (version 2.0), 1μl of the extracted RNA from each of the 96 samples processes separately into the TempO-Seq Human Whole Transcriptome Assay (BioSypder, index set C, #200615). A total of sixteen cycles of PCR, on a QuantStudio 6 Flex Instrument (ThermoFisher), were used to amplify the 96 ligated samples. The 96 samples were pooled into one library and then purified using NucleoSpin Gel and PCR clean-up kit (Macherey-Nagel, #740609). The final library was sequenced according to the TempO-Seq Assay User Guide (version 2.0, 50 bp read1 x 9 bp index1 x 9 bp index2) on the NextSeq 500/550 High Output v2 kit (Illumina, 75 cycles, #FC-404-2005) on a NextSeq 500 (Illumina), at a final concentration of 1.6 pM with 2% PhiX. For sequencing, the BioSpyder TempO-Seq Custom Index 1 Sequencing Primer was used at 0.3 μM final concentration diluted in HT1 buffer (illumina, see nextseq-custom-primers-guide-15057456-01). Final library concentration was determined via the mean of three replicates using the Qubit dsDNA BR Assay (invitrogen, #Q32853) on a Qubit 2 Fluorometer (Invitrogen).

**Laser Capture Microdissection (LCM)**

Twelve samples (6 MSA, 6 HC) from the C1 were used for Laser Capture Microdissection of Oligodendrocytes in the cerebellar white matter. Fresh frozen brains were sectioned on a cryostat at 10µm thickness, mounted on PEN membrane glass slides (Applied Biosystems) and stored immediately at -80°C until use. Before sectioning and between each tissue block, the knife holder and antiroll plate were wiped carefully with 100% ethanol to avoid cross-contamination. Oligodendrocytes were stained by using a modified H&E staining protocol adapted from [16]. A total of 300 oligodendrocytes per sample were captured using Arcturus CapSure Macro LCM Caps (Applied Biosystems) with the following settings: UV speed at 676 um/s and UV current at 2%. RNA was extracted immediately after cell capture using the Arcturus PicoPure RNA Isolation Kit (Applied Biosystems). RNA quality was tested with the Agilent BioAnalyzer and RIN’s averaged at 5.5 after extraction. For library preparation the SMARTer® Stranded Total RNA-Seq Kit - Pico Input (Clontech/Takara) was used. Libraries were validated on a 4200 TapeStation (Agilent Technologies) and quantitated via qPCR using Kapa’s Library Quantification Kit (Kapa Biosystems) on the QuantStudio 6 Flex Instrument (ThermoFisher). Samples were sequenced (2 x 75 bp paired-end run) on the Illumina HiSeq2500.

**Differential expression analysis**

The normal distribution of age, and PMI in MSA cases and HC was assessed using the Shapiro-Wilk test. The means between groups were compared using the t-test or Wilcoxon rank-sum test, depending by the data distribution. The distribution of age was compared using the Fisher’s Exact test. All these analysis were conducted using the R v3.3.1 software [17].

Raw sequencing data were demultiplexed and converted to FASTQ files using bcl2fastq Conversion Software v2.17 (Illumina, San Diego, CA). Quality controls on FASTQ files were conducted using MultiQC software v0.9 [7]. The reads were aligned to the Human reference genome (GRCh37) using the Spliced Transcripts Alignment to a Reference (STAR) software v2.5 [5]. Aligned reads were summarized as gene-level counts using featureCounts 1.4.4 [13]. Outliers and batch effects detection were conducted through Principal Component Analysis (PCA), using R software v3.3.1 [17], and quality reports were generated using FastQC 0.11. and Qualimap 2.1.3 [15]. Gene expression differential analyses between MSA cases and controls were conducted using the R package DESeq2 v1.14.1[14], including age, gender, PMI as covariates in all the comparison (except for gender in the C1 differential analysis including all samples because completely matched). The sample origin (Brain Bank) was included as covariate in the C1 analysis (including the LCM data), with the exception of the total sample analyzed because the samples were matched between cases and controls. The method implemented in DESeq2 fits a generalized linear model (GLM) for each gene, modeling reads counts following a negative binomial distribution. The Logarithmic Fold Change (FC), expressed as Log2 Fold Change, is estimated with an Empirical Bayes procedure, whereas the significance is assessed with a Wald Test [14]. Additionally, genes including extreme outliers counts are detected and excluded from the results using Cook’s distance, considering as cutoff the 99% quantile of the F(p,m-p) distribution (with p the number of parameters including the intercept and m number of samples) [4]. The p-values were corrected for multiple testing using the False Discovery Rate (FDR) method (Benjamini and Hochberg, 1995), and genes were annotated using the R-package BioMart v2.30.0 [6]. We considered as significant all the genes with adjusted p-value (adj-p) < 0.05.

**P-value combination**

The results from the two cohorts were combined using a meta-analytic approach based on the weighted-Z method [22]. The input consisted of the *p*-values obtained from the differential expression analysis. Since the Z-Weighted test assumes 1-tailed *p*-values, we converted the 2-tailed nominal *p*-value to1-tailed *p*-value using the following formula when the Log2 Fold Change was > 0: *p*_1Tailed_= *p*_2-Tailed_/2. Otherwise, we used the following formula: *p*_1Tailed_ = 1-(*_P_*_2Tailed/2_). The uncorrected *p*-values were weighted using the sample sizes of the two datasets, and combined using the *combine.test* function with the “*logit*” option included in the R-package *survcomp* [19]. Finally, the combined 1-tailed *p*-values were converted in 2-tailed *p*-values and adjusted for multiple testing with the FDR method.

**Cell specific Expression**

We classified the genes detected in the differential expression analysis using an external database of expression values from different types of cells isolated from mouse cerebral cortex [24]. We computed an enrichment score for each cell type and gene, dividing the *fpkm* expression value in a given cell by the average in the other cell types. The gene was assigned to a specific cell type when the enrichment score for one or multiple cell types was greater than 0.75, and in the other cell types was less than 0.25. Otherwise, the cell was classified as “mixed”. The enrichment of cell specific genes was investigated across DEGs and co-expression modules using a hypergeometric test as implemented in the R function *phyper*. Results were adjusted with the FDR method.

**Enrichment and functional network analysis**

List of DEGs were analyzed for Gene Ontology enrichment using the R-package *anRichmentMethods*, adjusting the *p*-value with the FDR method. The same gene lists were also analyzed using *HumanBase* (<https://hb.flatironinstitute.org/gene>)*,* constructing tissue-specific functional networks [8].

We conducted an additional analysis looking for enrichment of Alzheimer’s Disease (AD) genes in our MSA-C results using as reference gene set the data public available from the Accelerating Medicines Partnership – Alzheimer’s Disease (AMP-AD; accession number syn14237651). We downloaded the differential expression results from the Mount Sinai [21], Mayo [2] and ROSMAP studies [1], including data from 7 brain regions: Temporal Cortex (TCX), Superior Temporal Gyrus (STG), Dorso-Lateral Prefrontal Cortex (DLPFC), Frontal Pole (FP), Inferior Frontal Gyrus (IFG), Parahippocampal Gyrus (PHG), and Cerebellum (CBE). We selected the “Diagnosis” model, where the AD and control definitions were harmonized by defining cognitive scores, Braak staging, and tau pathology. Further details can be found at <https://www.synapse.org/#!Synapse:syn14237651>. We filtered the DEGs for each brain region at different cutoffs: adj *p* < 0.05, adj *p* < 0.01, adj *p* < 0.001, and adj *p* < 0.0001, and they were used as reference AD gene sets. As validation, we also tested the less variable genes between AD and controls in the AMP-AD dataset using *p* > 0.500 as cutoff. As test set we used all the genes detected in MSA-C cohort ranked by log2 FC, without any p-value filtering. The analysis was conducted with the R-package *fgsea using* 100,000 permutations and adjusting the p-values with the False Discovery Rate Method [3].

**WGCNA analysis**

An alternate way of obtaining meaning from data is provided by the WGCNA algorithm [23]. This algorithm extracts additional information by finding clusters of highly correlated genes that have been demonstrated to be functionally related [12]. The analysis was conducted using the *WGCNA* R package [10]. Genes with low counts from both C1 and C2 (MSA-C subgroup) were filtered out (< 10 average counts across all samples) and the expression data matrix was normalized using the *vst* function included in the DESeq2 package [14]. To remove confounding factors, the matrix of expression data was adjusted form age, sex, PMI and sample source using the function *removeBatchEffect* as implemented in the limma R-package [18]. We computed the Median Absolute Deviation (MAD) for each gene in both datasets separately, and we removed the 50% of genes with lowest MAD, aiming to exclude low variable genes usually representing noise.

The coexpression network was computed for the C1. First, we computed soft-thresholding power (*β*)*,* using the *pickSoftThreshold* function. Then, we plotted the values against the scale-free fit index, and selected the lowest power for which the scale-free topology fit index curve flattens out upon reaching a *r^2^* = 0.900 [23]. We generated the coexpression network and identified the resulting clusters using the function *blockwiseModules* with the following parameters: *TOMtype*: “signed”, *minimum module size* = 30, *mergeCutHeight* = 0.25, *deepSplit* = 2; *reassign threshold* = 1.0E-06*,* and *pamRespectsDendro =* “FALSE”*.*

We computed the eigengenes values for each individual and module in our dataset by singular value decomposition (SVD) [9]. The eigengenes were compared by module between MSA-C and HC using a linear model as implemented in *limma*, adjusting the *p*-values for multiple testing accounting for the number of modules using the FDR method [3]. Covariates were not included in the model since we used the adjusted expression matrix, minimizing the confounding factors. To rank the genes, we computed the gene-module membership using the WGCNA package with the *Pearson’s* correlation, as well as the gene-trait significance, in this case the disease status. The gene-module membership is proportional to the correlation of a specific gene with the eigengene module.

We conducted Gene Ontology (GO) enrichment analysis on the modules associated with the disease status by the means of the R-package *anRichment*, using as background the intersection of given genes and genes present in GO. P values of the GO enrichment analysis were adjusted using the FDR method [3]. The enrichment for genes expressed in specific cell types was conducted using as reference gene sets the gene specifically expressed in the 5 cell types from Zhang et al. (Zhang et al., 2014) and test set all the genes ranked by module memberships for the module associated with the disease status.

The top hubs in the coexpression modules were identified using the function *chooseTopHubInEachModule* in the WGCNA package. Relevant co-expression networks were exported and then visualized using *Cytoscape v3.7.1*. Finally, we checked the module preservation in C2 using the *modulePreservation* function with 1,000 permutations, representing the results in a plot including the number of genes for each module (X-axis) and the Zsummary statistics (Y-axis). The Zsummary statistics summarize evidence that a module is preserved more significantly than a random sample of all network genes. Values of Zsummary greater than 10 correspond to strong evidence for module preservation, between 2 and 10 moderate evidence, and Zsummary < 2 indicate weak module preservation [11].

**2. Supplementary results**

**Figure S1.** PCA analysis conducted on cohort 1 after transformation of the raw counts using regularized log2 transformation. Samples were labeled by diagnostic status. No significant outliers were detected. The final sample size was: MSA = 19, HC = 19.

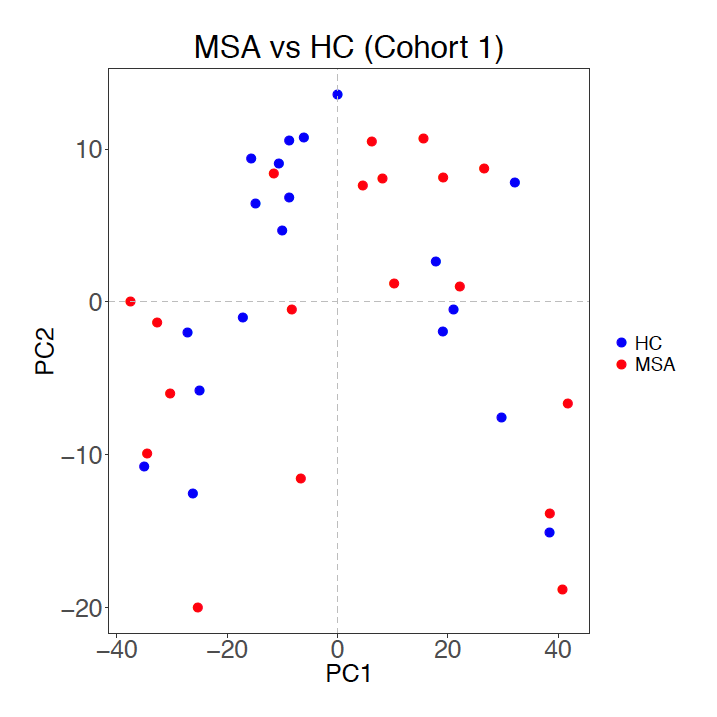

**Figure S2.** PCA analysis conducted on cohort 2 after transformation of the raw counts using regularized log2 transformation. Samples were labeled by diagnostic status. One outlier was identified (A) and removed (B). The final sample size was: MSA = 47; HC = 47.

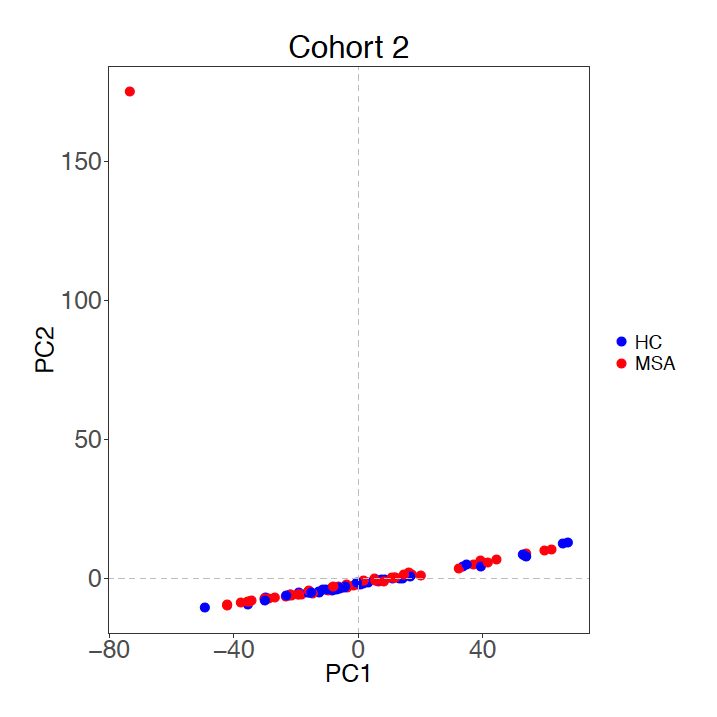

**A**

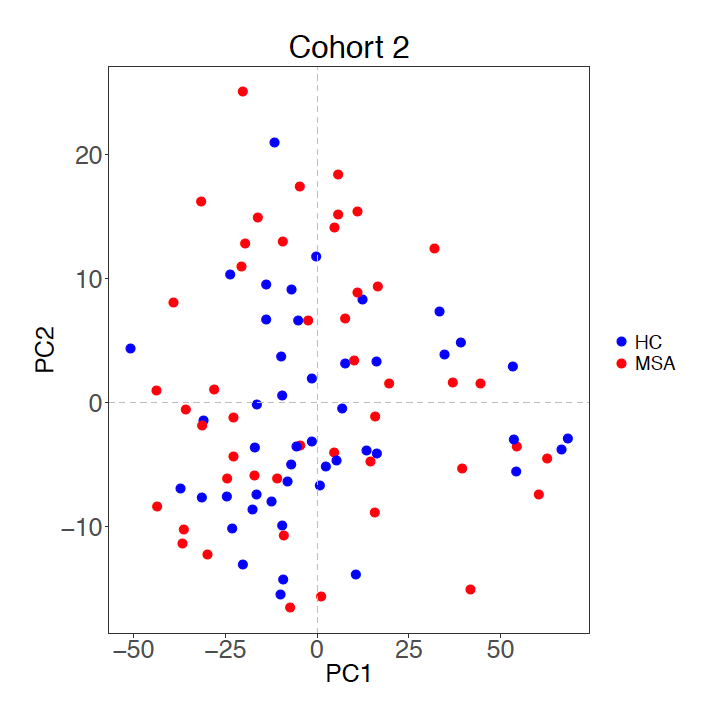

**B**

**Figure S3.** PCA analysis conducted on oligodendrocytes LCM data from cohort 1 after transformation of the raw counts using regularized log2 transformation. Samples were labeled by diagnostic status. Three extreme outliers were identified (A) and removed (B). The final sample size was: MSA = 4; HC = 5.

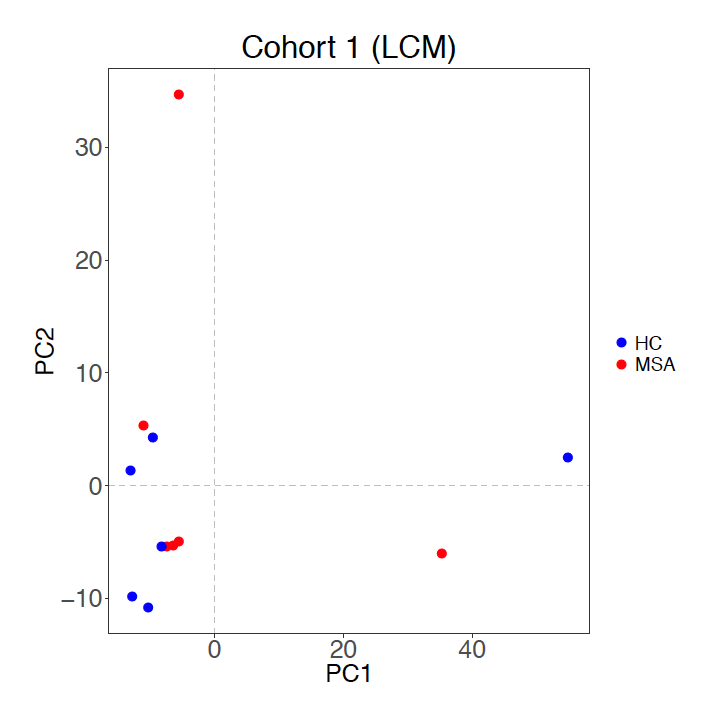

**A**

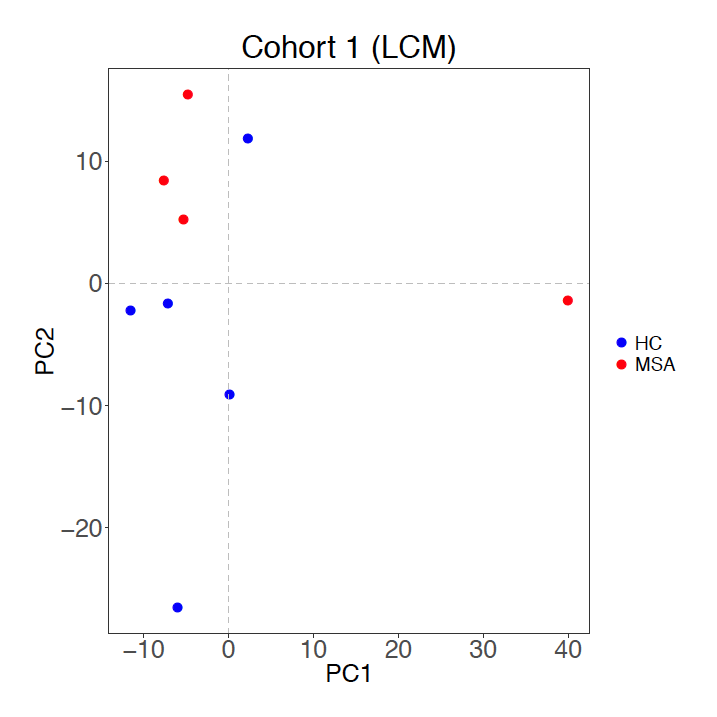

**B**

**Figure S4.** Number of differentially expressed genes (adj p < 0.05) detected in cohort 1 and cohort 2 for each MSA subtype (and in the analysis MSA-C + MSA-P, indicated as “MSA”, as well as in the p-value combination analysis (CB).

The barplots are colored in red and blue for the number of upregulated and downregulated genes, respectively. We observed the larger number of DEGs in the MSA-C subtype in both cohort specific and combined analyses.

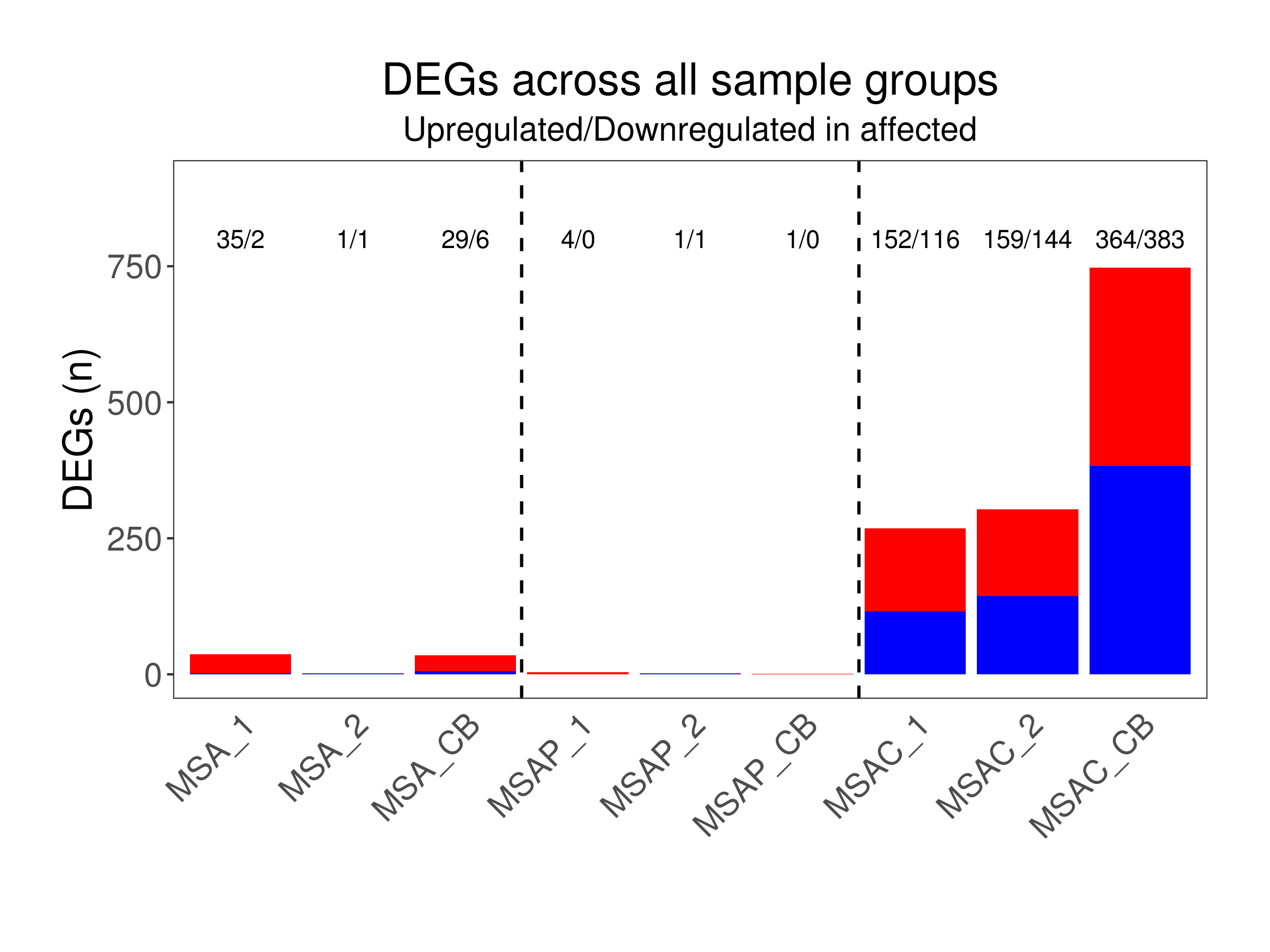

**Figure S5**. The log2 FC between cohort 1 and 2 was compared using the Spearman correlation. In all the comparisons (MSA, MSA-P and MSA-C) we obtained a positive and statistically significant correlation (ρ > 0.204; p < 2.2E-16). In red genes with the same log2 Fold Change trend.

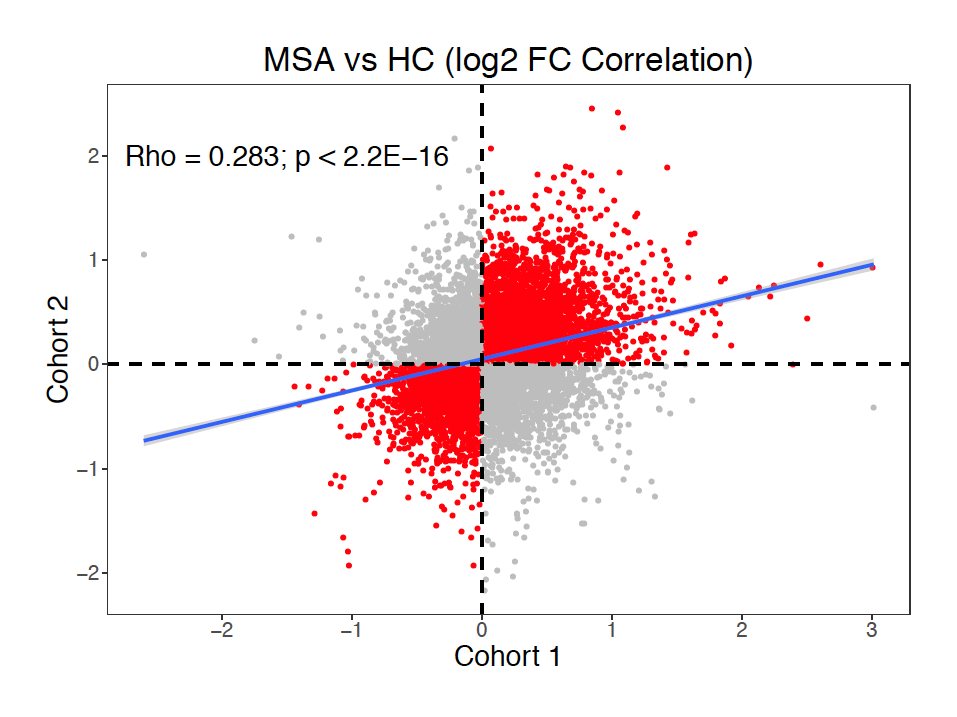

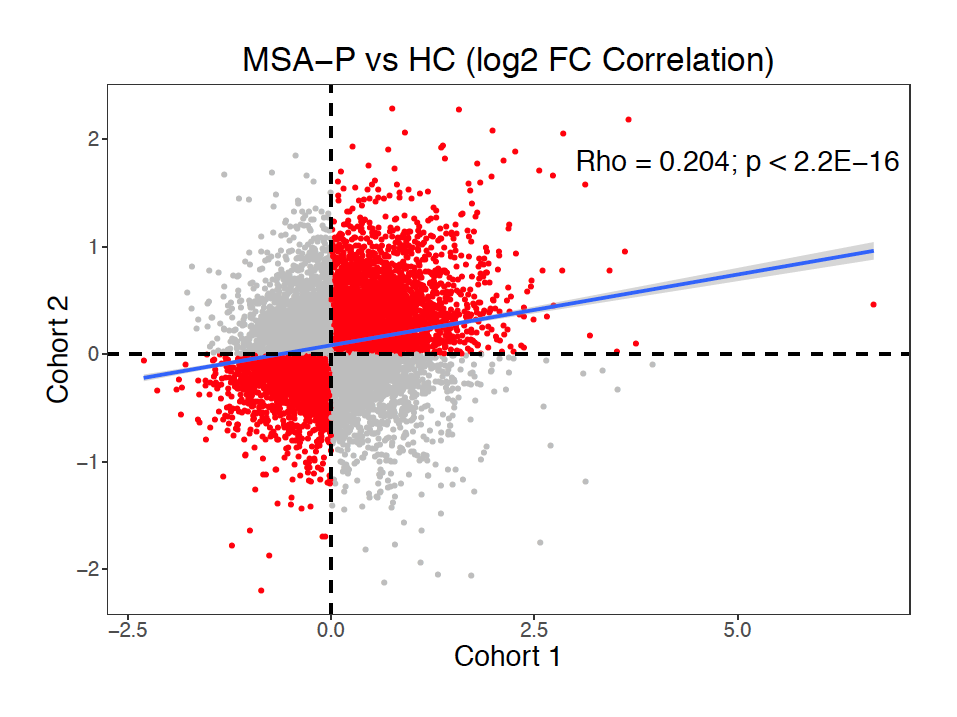

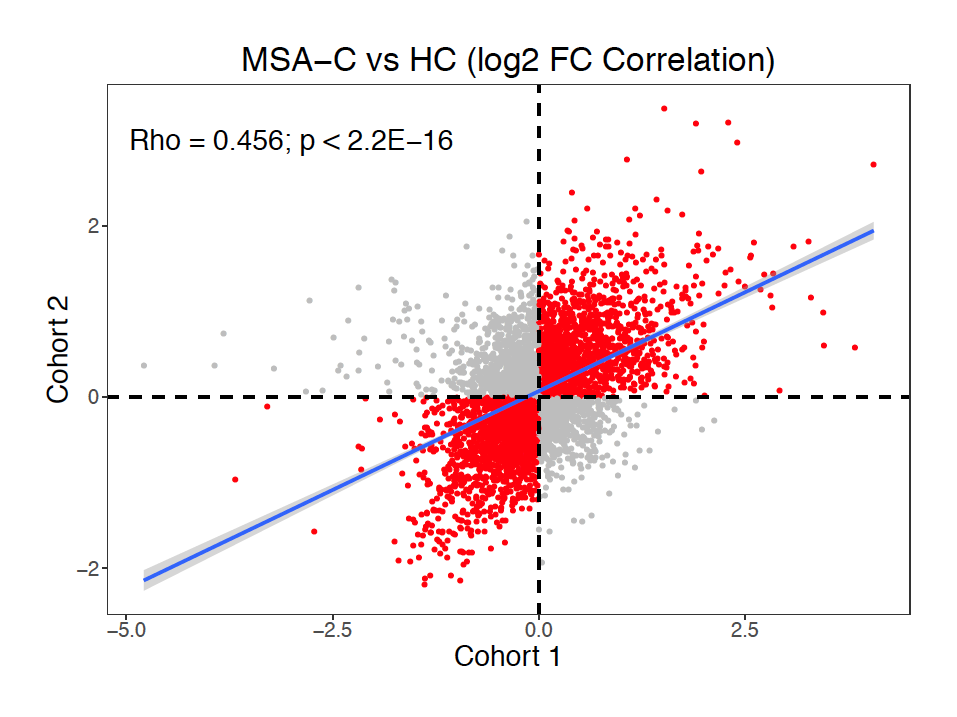

**Figure S6.** Volcano plot reporting the differential analysis between SND (top) and OPCA (bottom) vs HC in C2. We detected a total of 7 and 58 genes, respectively.

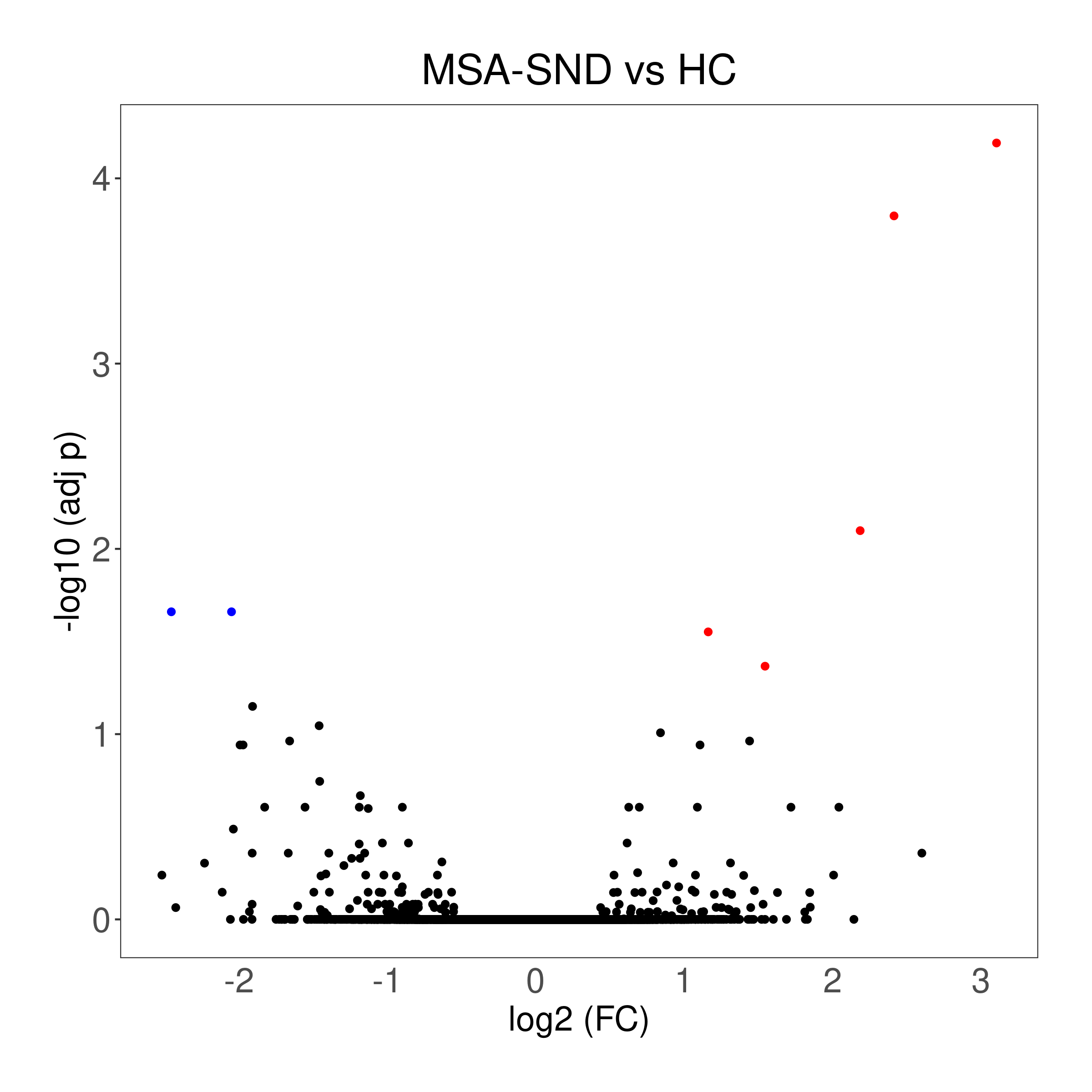

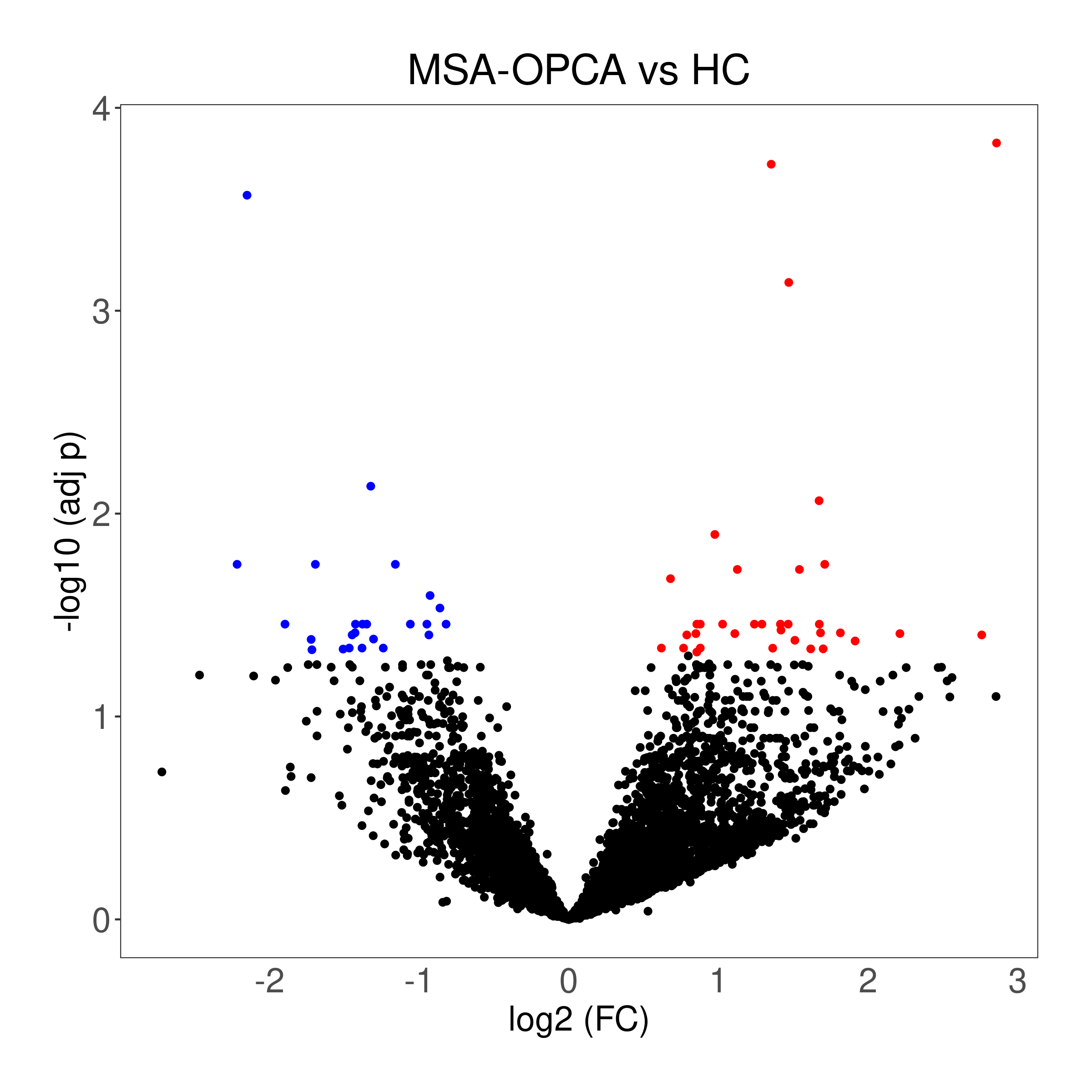

**Figure S7.** Log2 FC Correlation plots between MSA-P and SND patients (C2) (top) and between MSA-C and OPCA patients (bottom). We detected only one gene significant in both MSA-P/SND and 47 in both MSA-C/OPCA analyses (red dots).

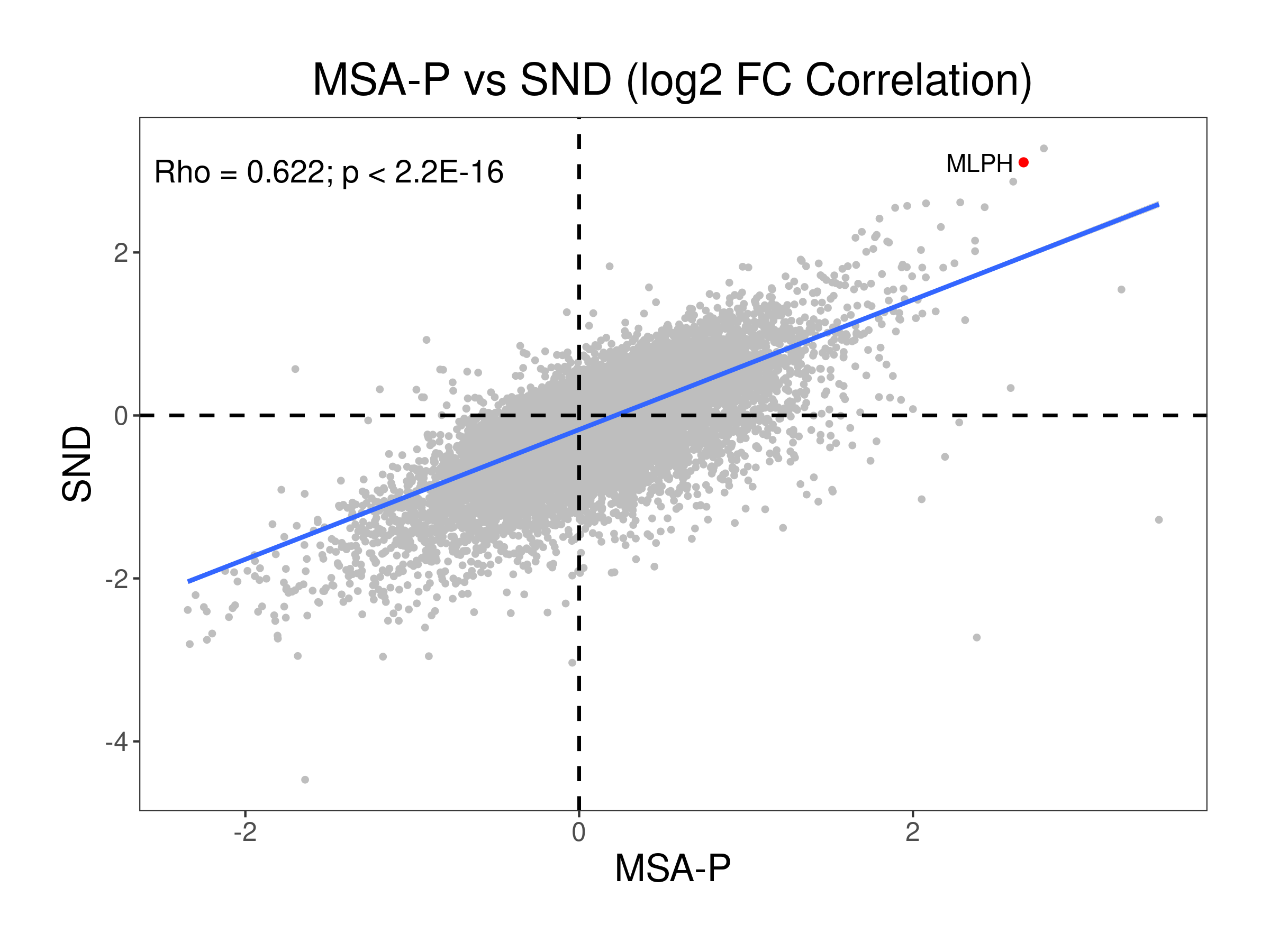

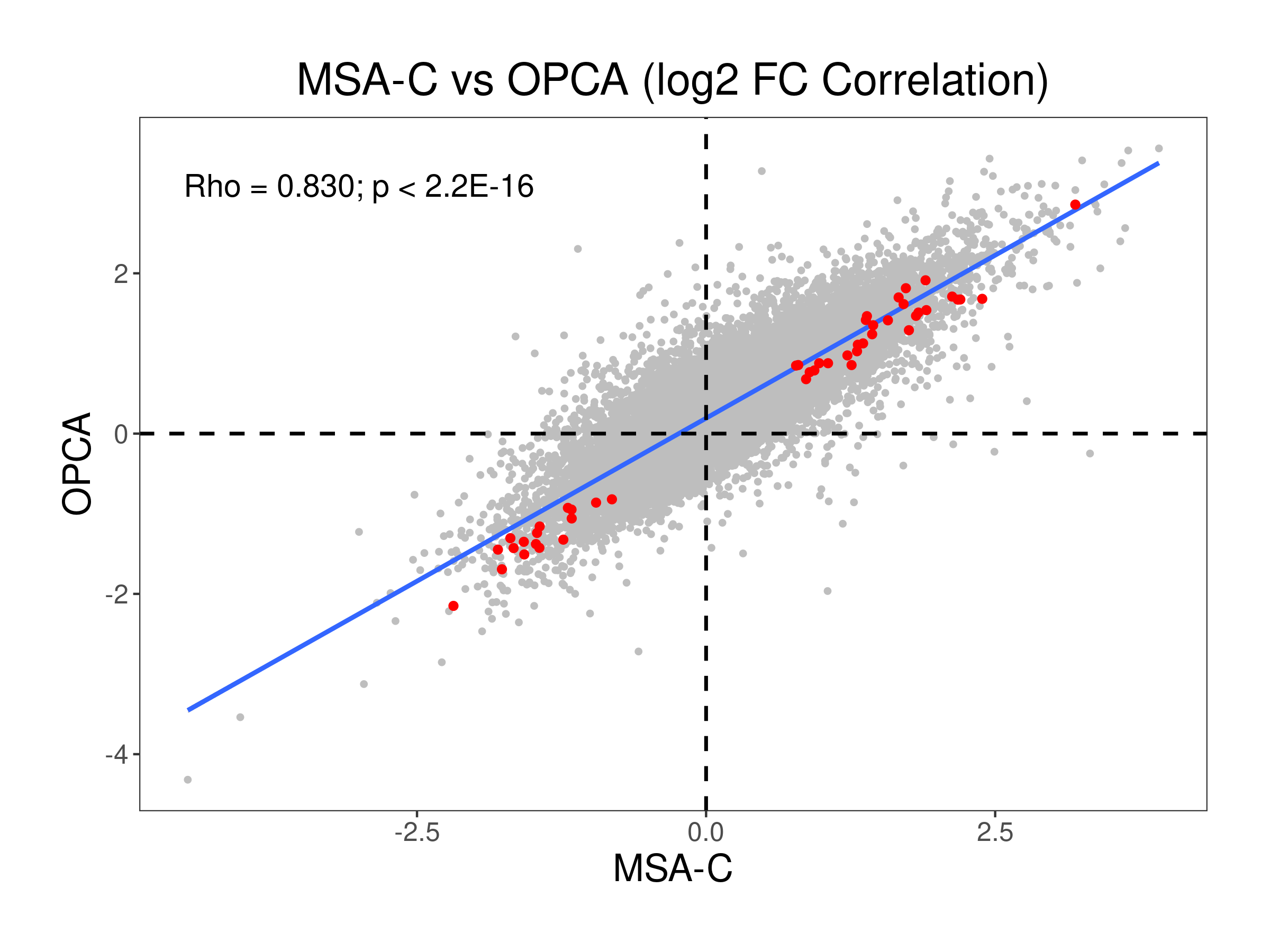

**Figure S8.** Results of the functional module discovery analysis using the 35 DEGs identified in MSA (MSA-P + MSA-C). The module M1 is enriched for cell-cell adhesion processes, and the module 2 is enriched for angiogenesis (*q* < 0.01).

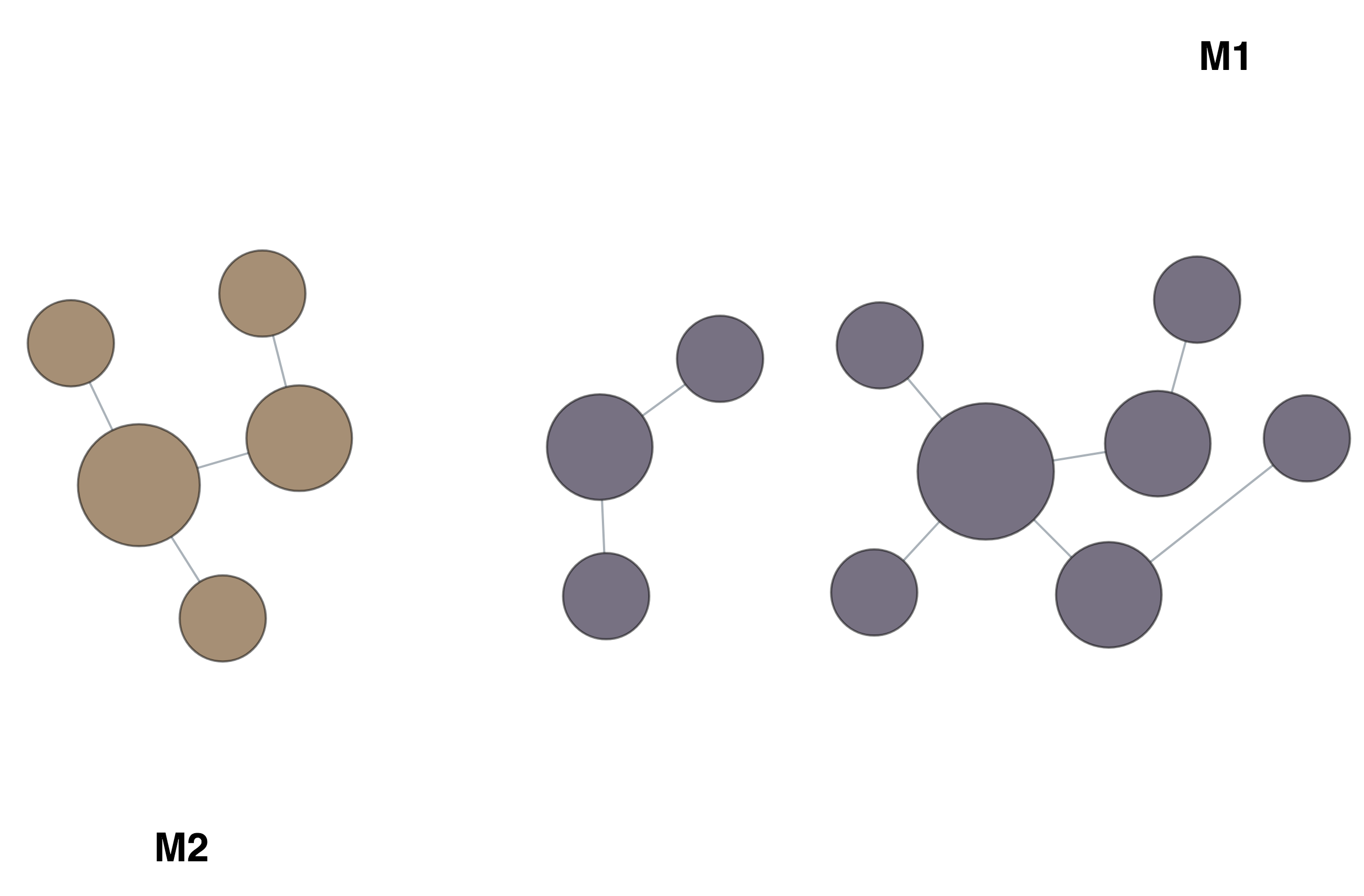

| **CLUSTER** | **TERM_NAME** | **GO_ID** | **Q-VALUE** | **GENES** |
| --- | --- | --- | --- | --- |
| M1 | *cell-cell adhesion* | GO:0098609 | 1.0E-02 | *SELL,BCL6* |
| M1 | *leukocyte cell-cell adhesion* | GO:0007159 | 3.4E-03 | *SELL,BCL6* |
| M1 | *negative regulation of cell differentiation* | GO:0045596 | 5.3E-03 | *NFKBIA,LDLRAD4* |
| M2 | *angiogenesis* | GO:0001525 | 3.4E-03 | *COL4A1,COL4A2* |
| M2 | *blood vessel morphogenesis* | GO:0048514 | 3.4E-03 | *COL4A1,COL4A2* |
| M2 | *blood vessel development* | GO:0001568 | 3.4E-03 | *COL4A1,COL4A2* |
| M2 | *vasculature development* | GO:0001944 | 3.4E-03 | *COL4A1,COL4A2* |
| M2 | *cardiovascular system development* | GO:0072358 | 3.4E-03 | *COL4A1,COL4A2* |
| M2 | *tube morphogenesis* | GO:0035239 | 3.4E-03 | *COL4A1,COL4A2* |
| M2 | *tube development* | GO:0035295 | 3.4E-03 | *COL4A1,COL4A2* |

**Figure S9.** Protein-protein network generated using the prioritization method using as input all the DEGs detected in MSA-C after p-value combination. The analysis was conducted using *WEBGESTALT* [20]. The hub gene *APP* is the large circle.

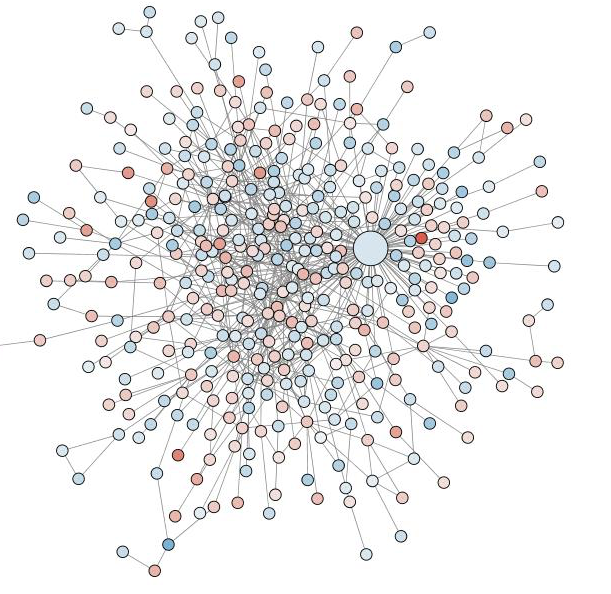

**Figure S10.** Top 20 GO enrichment functional class detected using all the differentially expressed genes detected in MSA-C. We detected a total of 625 significant functional classes (adj *p* < 0.05).

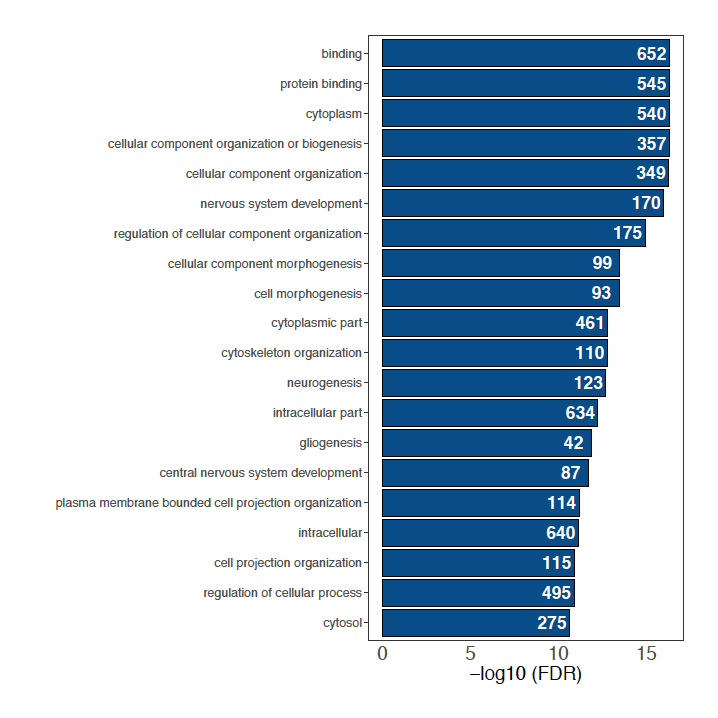

**Figure S11. Enrichment of AD genes in MSA-C dataset.**

(A) Enrichment plot for genes differentially expressed in the Temporal Cortex in AD (adj *p* < 0.05). The enrichment was confirmed using more conservative thresholds (adj p > 0.01; adj p < 0.001 and adj p < 0.0001).
(B) Enrichment plot for genes differentially expressed in the Parahippocampal Gyrus in AD (adj *p* < 0.05). The enrichment was confirmed using more conservative DEGs thresholds (adj *p* > 0.01; adj *p* < 0.001 and adj *p* < 0.0001).

**A**

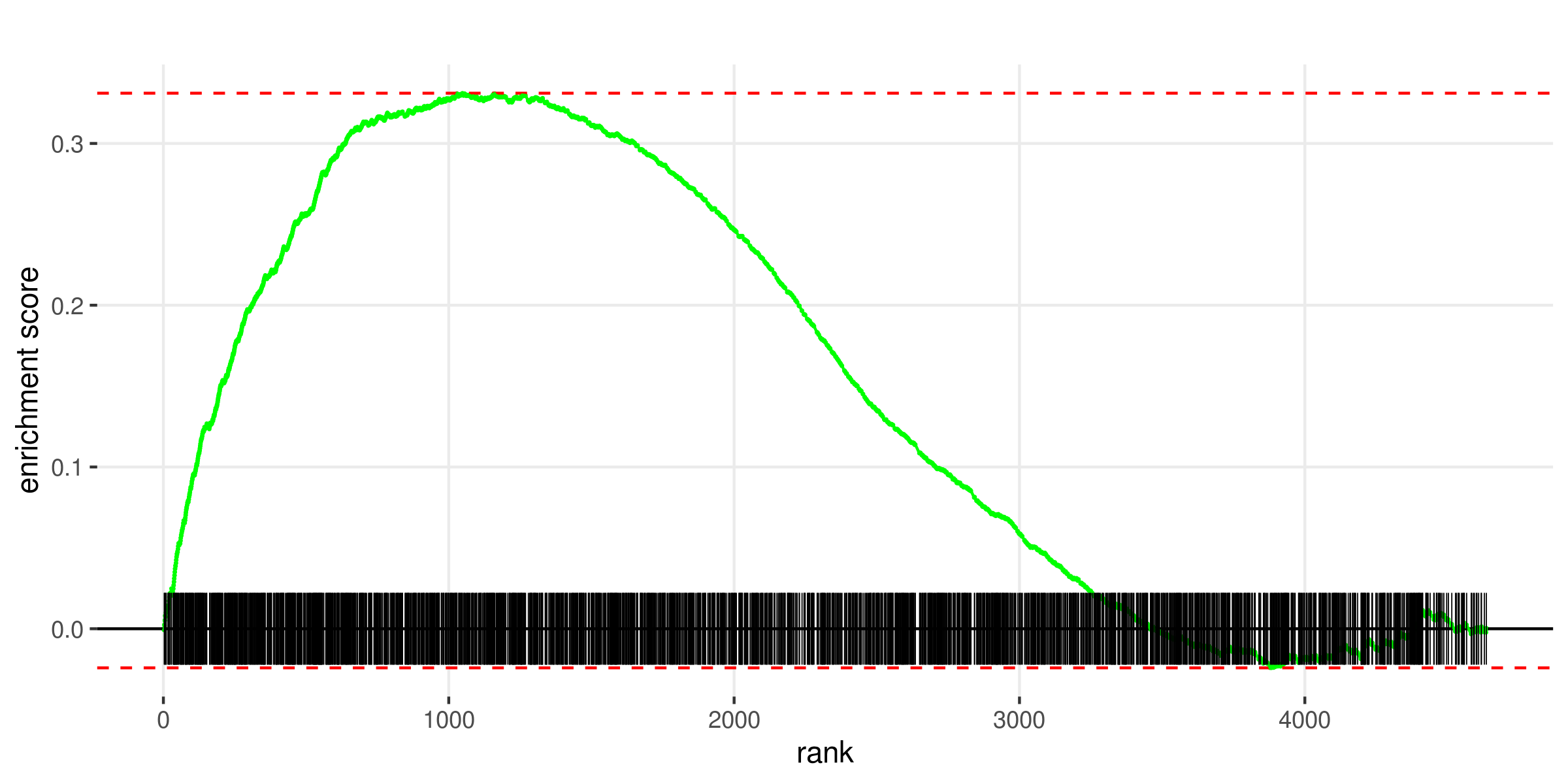

**B**

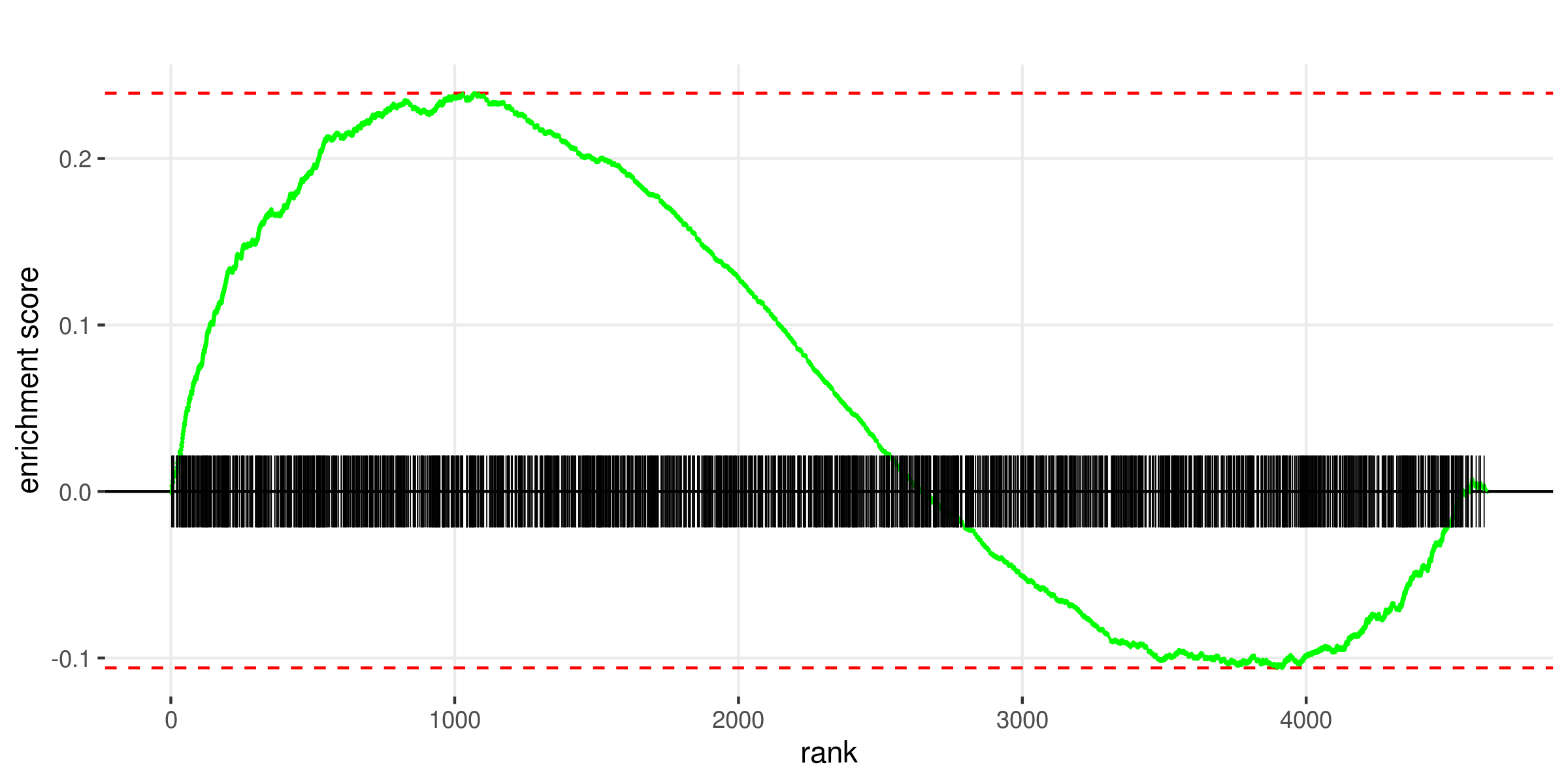

**Figure S12.** Results of the functional module discovery analysis using the 187 DEGs identified in LCM oligodendrocytes (MSA vs HC). The table showed below reports the GO top 3 significant enriched processes for each module.

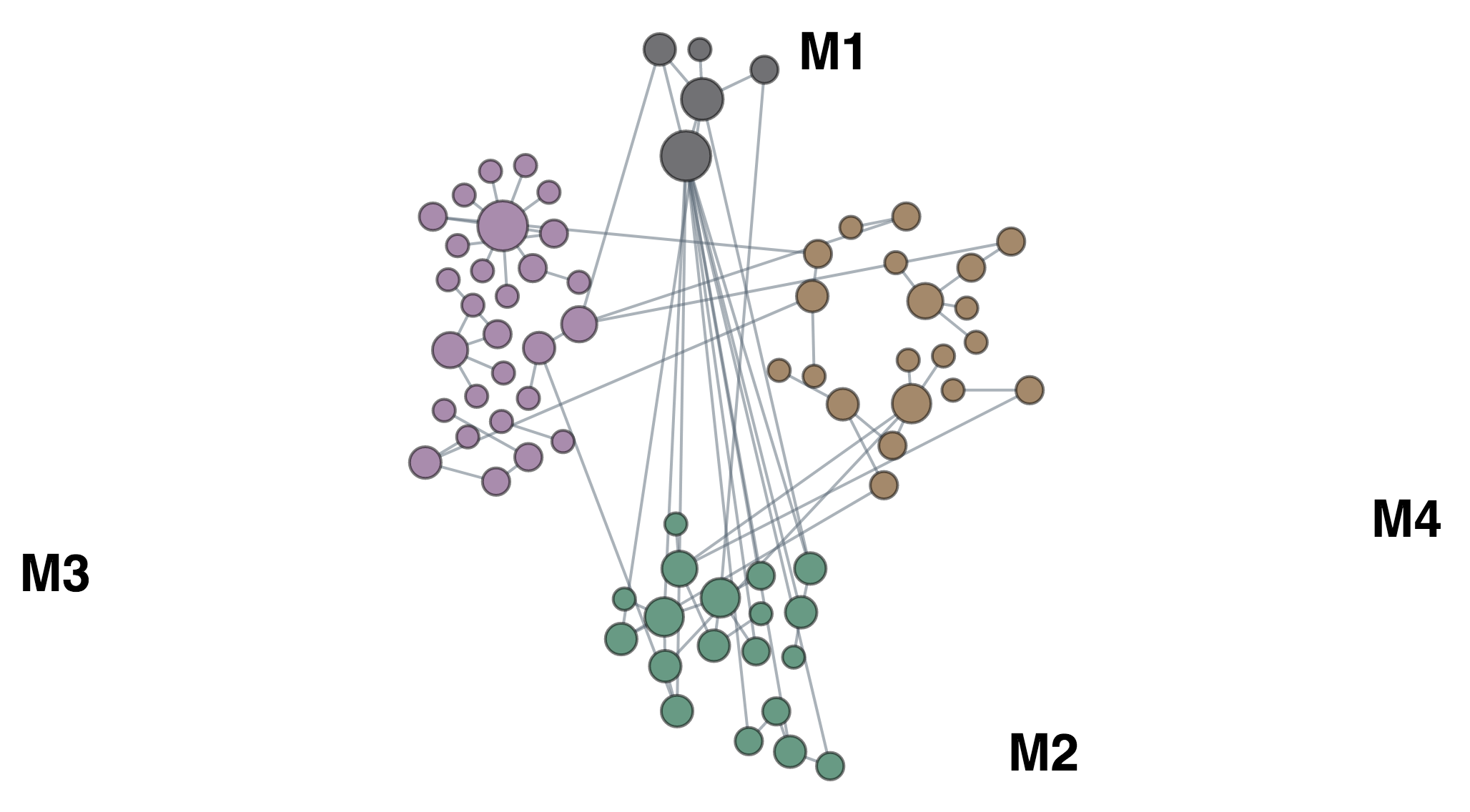

| **CLUSTER (GENES)** | **TERM_NAME** | **GO_ID** | **Q_VALUE** | **GENES** |
| --- | --- | --- | --- | --- |
| M1 (5) | RNA-dependent DNA biosynthetic process | GO:0006278 | 1.9E-03 | *PTGES3,WRAP53* |
|  | telomere maintenance via telomerase | GO:0007004 | 1.9E-03 | *PTGES3,WRAP53* |
|  | telomere maintenance via telomere lengthening | GO:0010833 | 1.9E-03 | *PTGES3,WRAP53* |
| M2 (19) | ncRNA processing | GO:0034470 | 2.5E-03 | *INTS8,DIMT1,MTREX* |
|  | ncRNA metabolic process | GO:0034660 | 4.9E-03 | *INTS8,DIMT1,MTREX* |
|  | rRNA processing | GO:0006364 | 5.7E-03 | *DIMT1,MTREX* |
| M3 (28) | leukocyte chemotaxis | GO:0030595 | 2.3E-02 | *PADI2,CHGA* |
|  | defense response to bacterium | GO:0042742 | 2.4E-02 | *CHGA,SIRT2* |
|  | negative regulation of response to external stimulus | GO:0032102 | 2.5E-02 | *PADI2,SIRT2* |
| M4 (20) | regulation of cell growth | GO:0001558 | 2.8E-02 | *NME6,BDNF* |
|  | cell growth | GO:0016049 | 2.8E-02 | *NME6,BDNF* |
|  | regulation of growth | GO:0040008 | 3.2E-02 | *NME6,BDNF* |

**Figure S13.** GO enrichment analysis results obtained using the LCM DEGs. We detected an enrichment for myelination processes, especially due to downregulated genes in MSA.

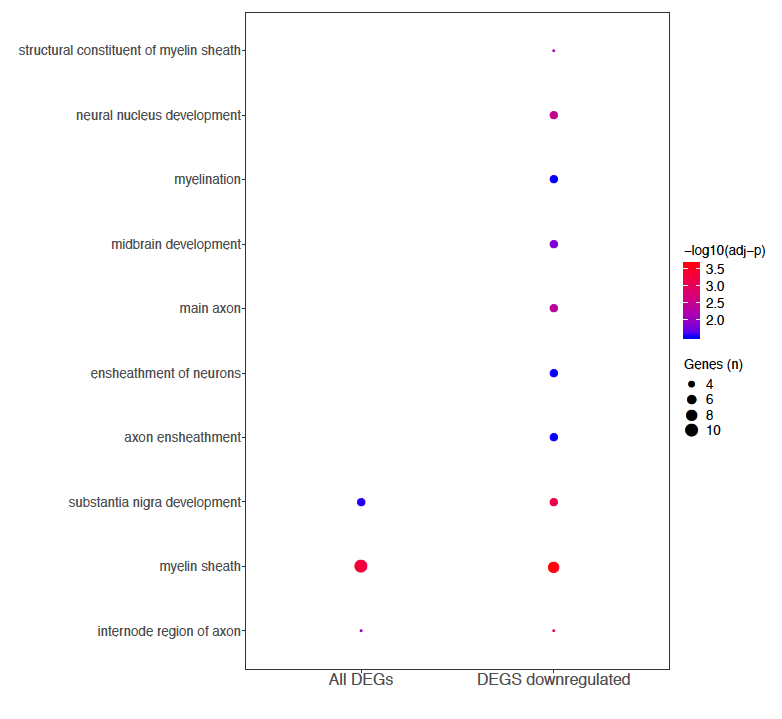

**Figure S14.** Hypergeometric test results of the cell specific genes classification. The number at the top indicates total number of gene for each class. Dashed line indicates statistical significance (adj *p* < 0.05). We detected a significant enrichment for astrocyte (A) and oligodendrocyte (O) genes.

EC = endothelial; N = neuron; M = microglia.

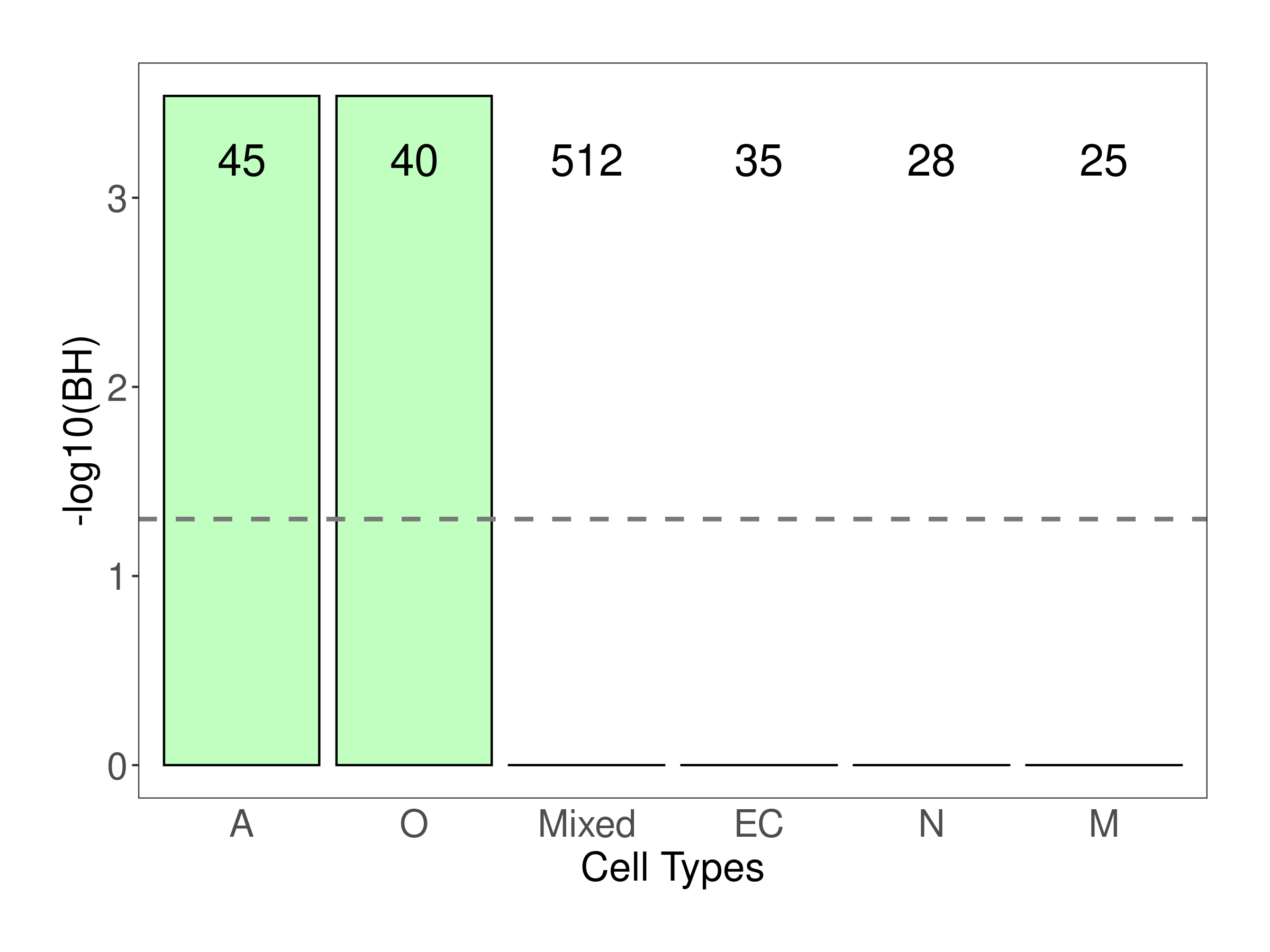

**Figure S15.** Results of the gene classification by cell type using our deconvolution method, including DEGs with adj *p* < 0.10 (A), and DEGs with adj p < 0.025 (B). The results confirm what we obtained using a more restrictive cutoff (adj *p* < 0.05).

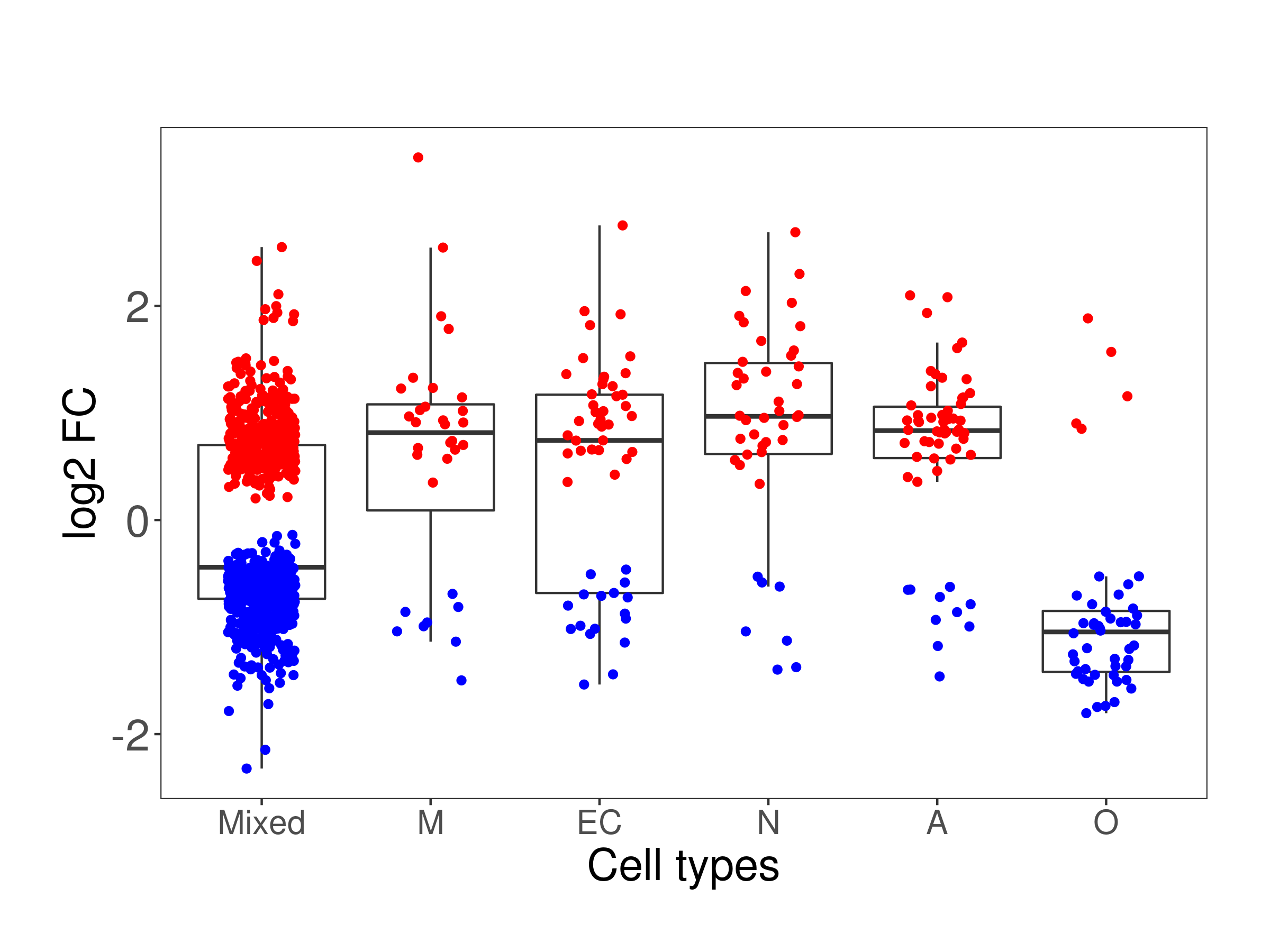

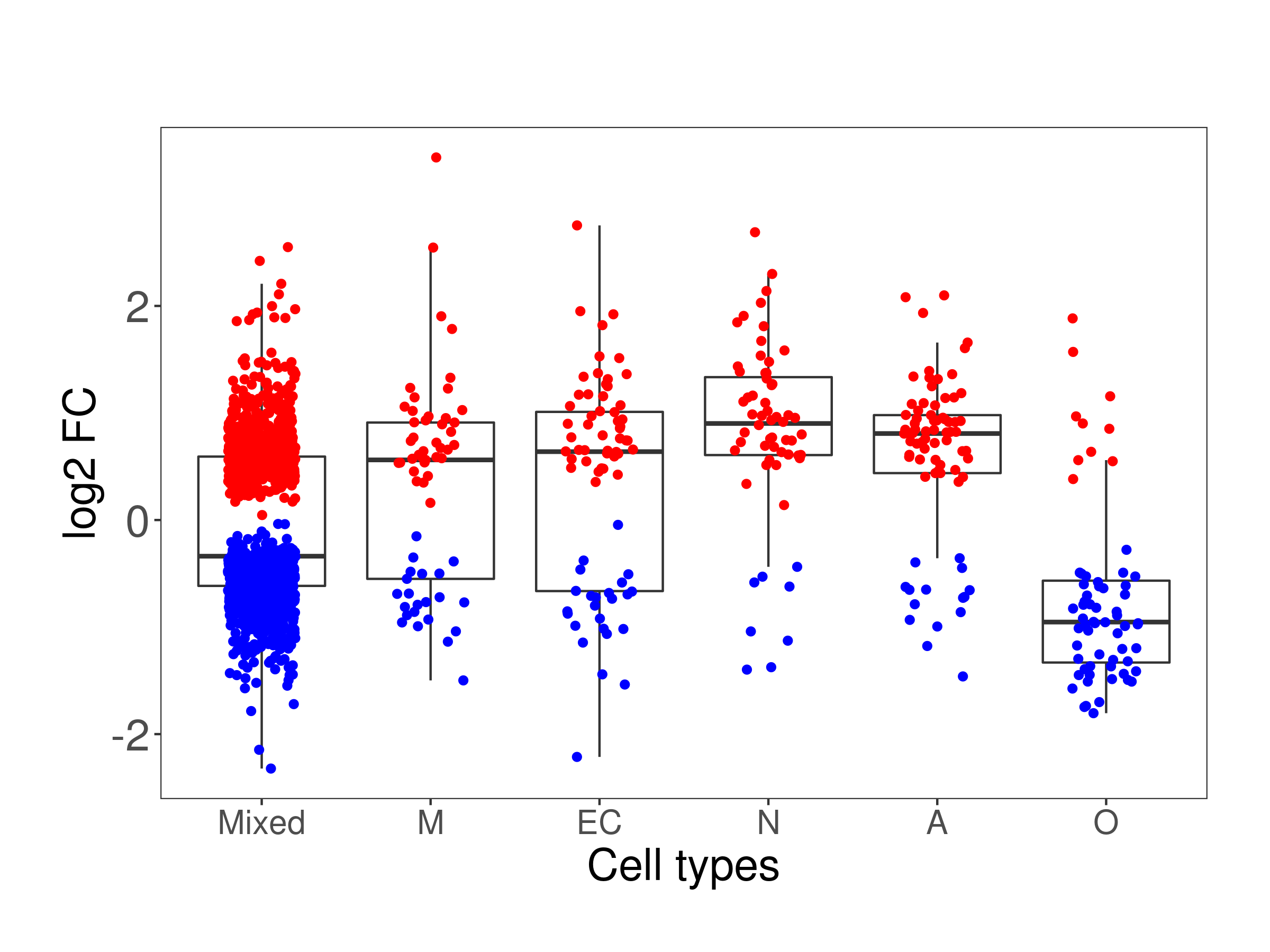

**Figure S16.** WGCNA analysis: Scale-free fit index (y-axis) as a function of the soft-thresholding power (x-axis). We selected 9 as soft threshold power for the WGCNA analysis.

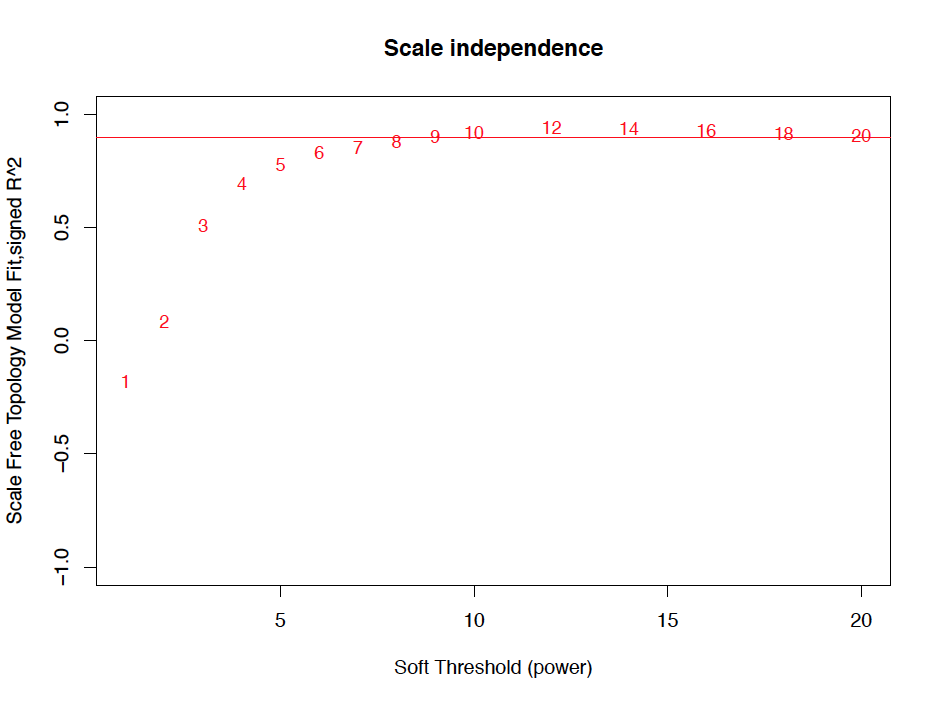

**Figure S17.** Dendrogram and heatmap showing the correlation between coexpression modules.

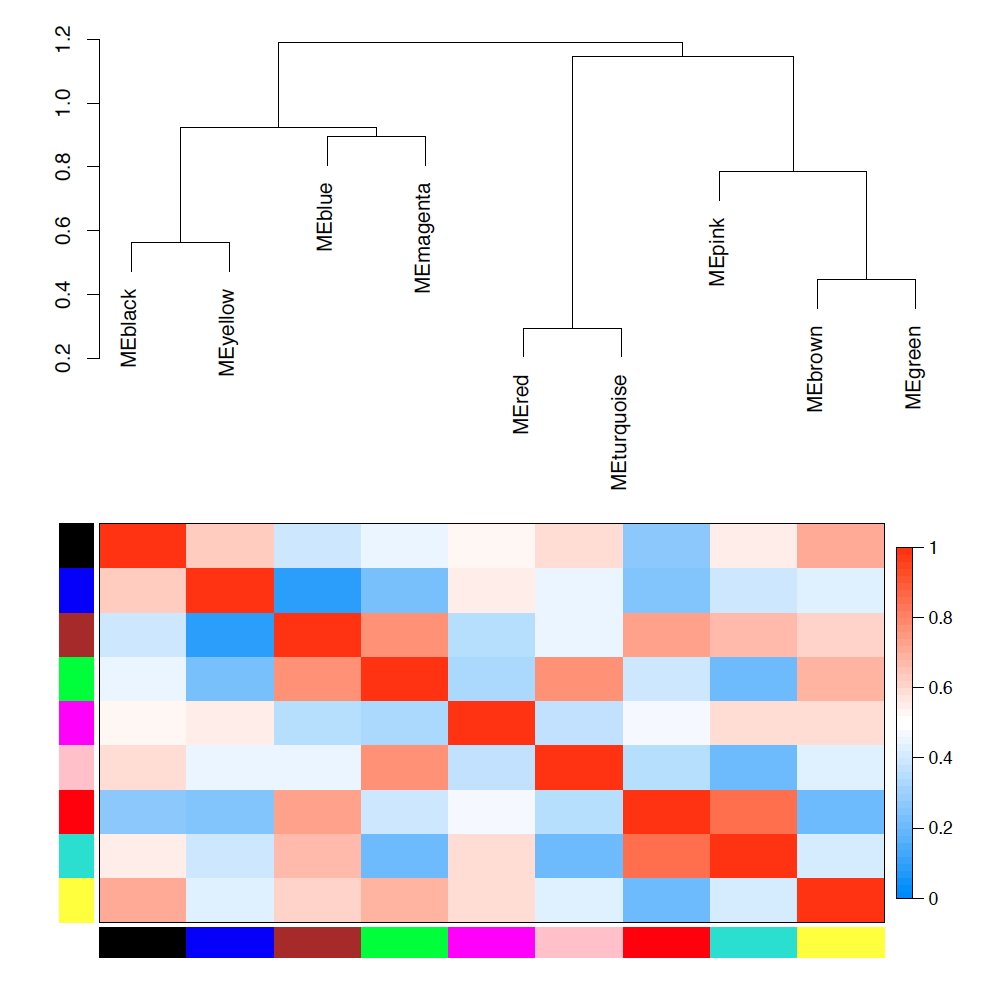

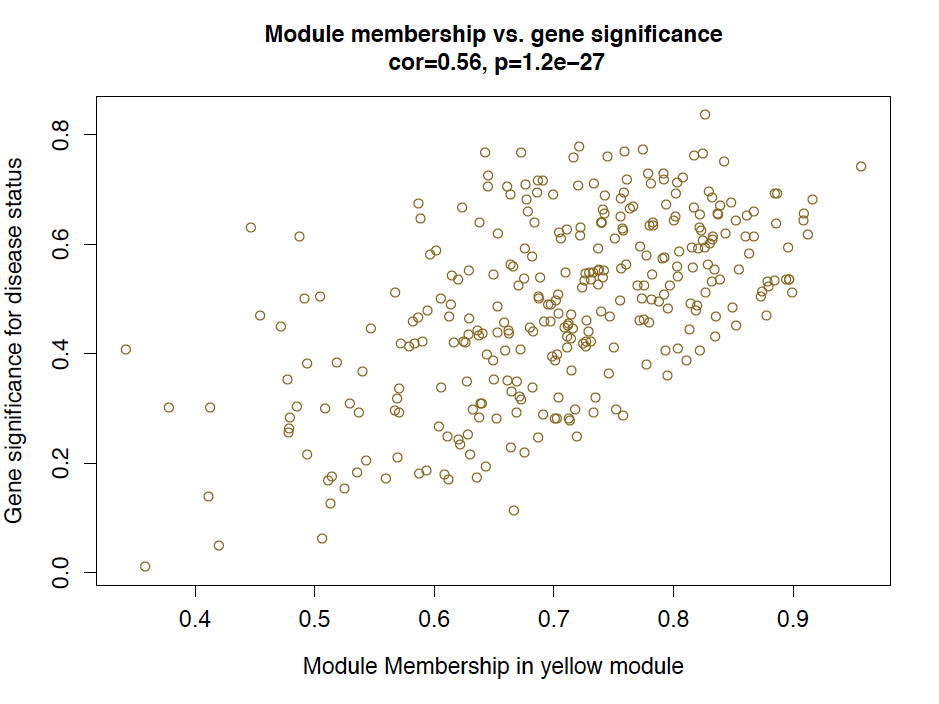
**Figure S18.** Module membership (x-axis) vs Gene-trait significance (y-axis) for the yellow and green modules, significantly associated with MSA-C.

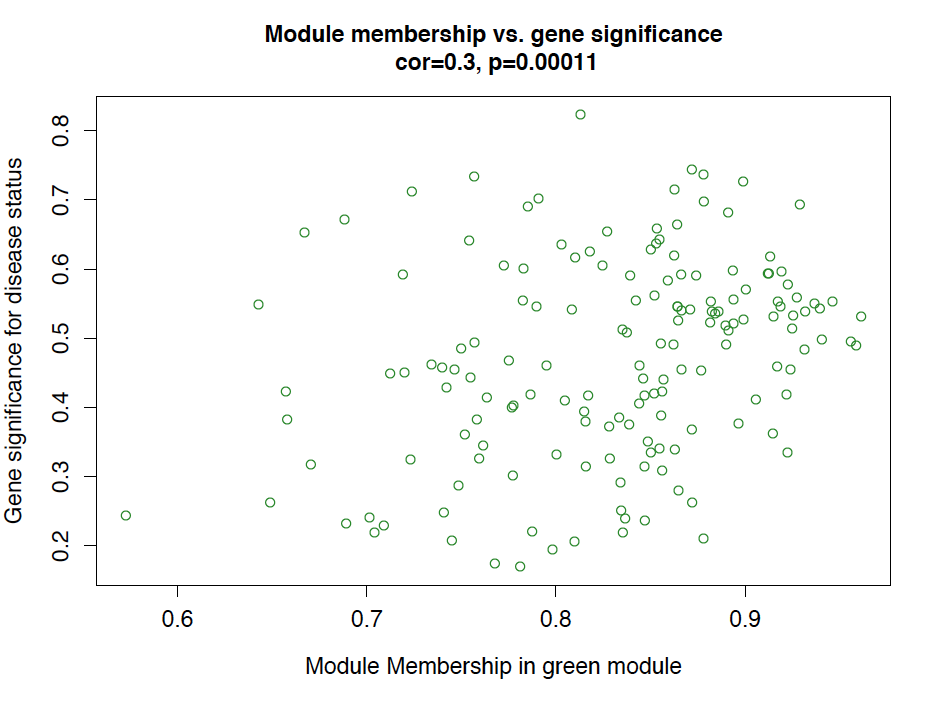

**Figure S19.** Module membership (x-axis) vs Gene-trait significance (y-axis) for the blue and brown modules, significantly associated with MSA-C.

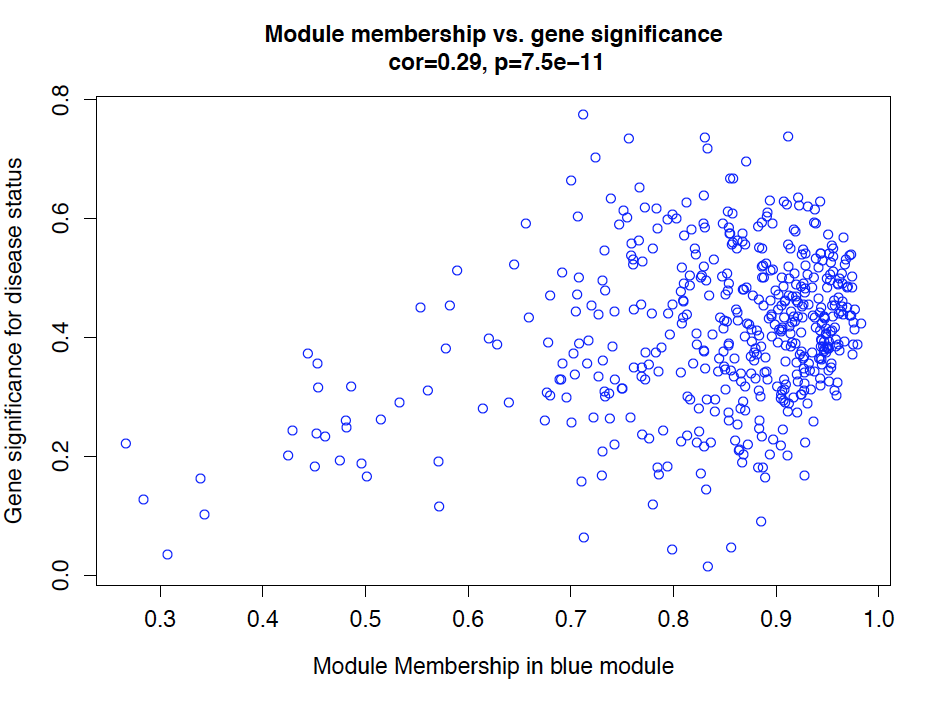

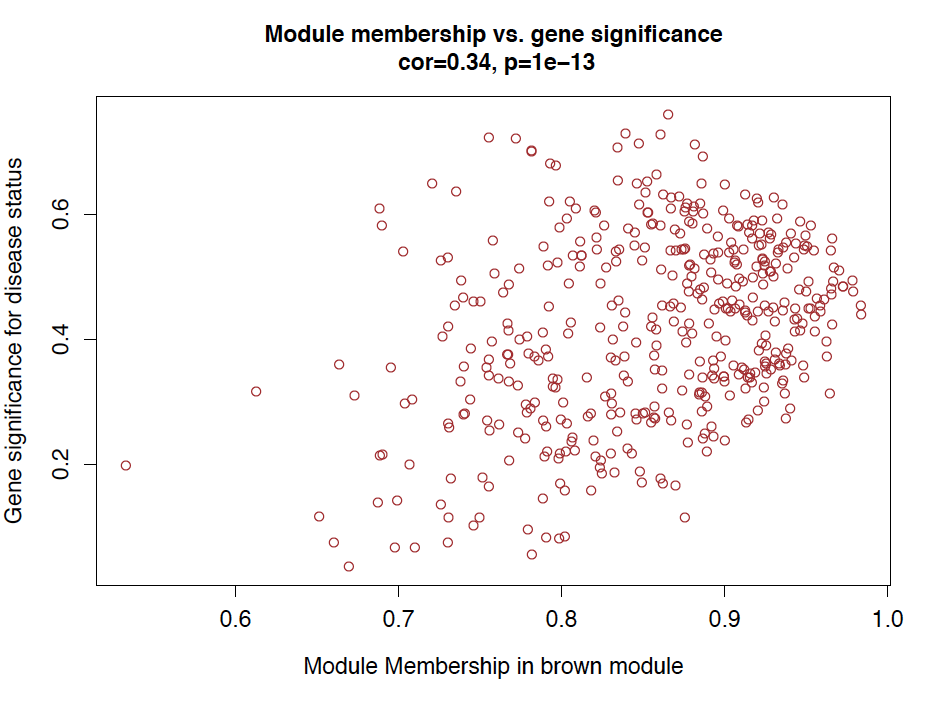

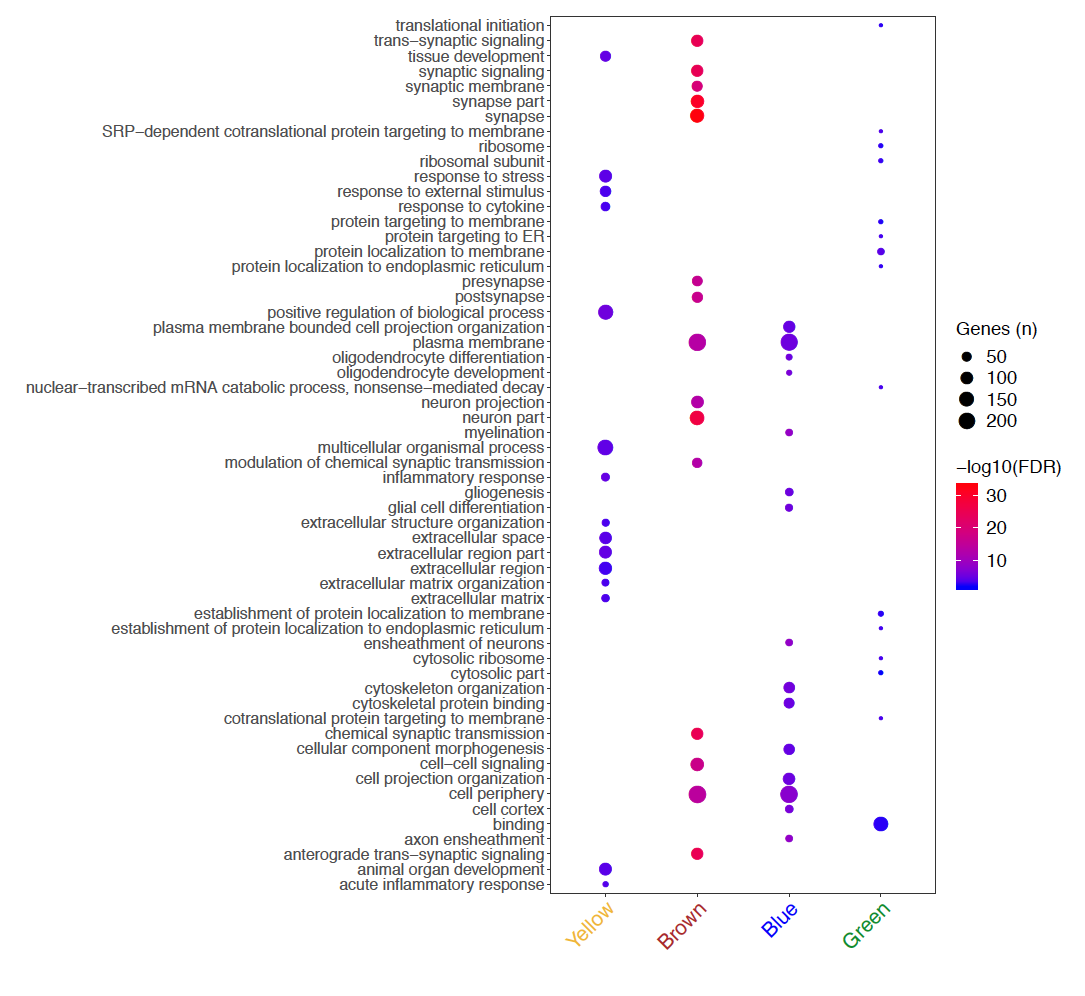
**Figure S20.** Dot plot representing the top 15 GO process enriched in the coexpression modules significantly different between MSA-C and HC.

**Figure S21.** Cell specific enrichment among the WGCNA modules associated with MSA-C in cohort 1. We observed significant enrichment (labeled with a star) of:

1) astrocyte (adj *p* = 3.3E-19) and endothelial genes (adj *p* = 2.8E-04) in the yellow module;
2) neuronal genes in the brown module (adj *p* = 2.5E-60);
3) oligodendrocyte genes in the blue module (adj *p* =7.7E-33).

A = astrocyte; EC = endothelial; M = microglia; N = neuron; O = oligodendrocyte.

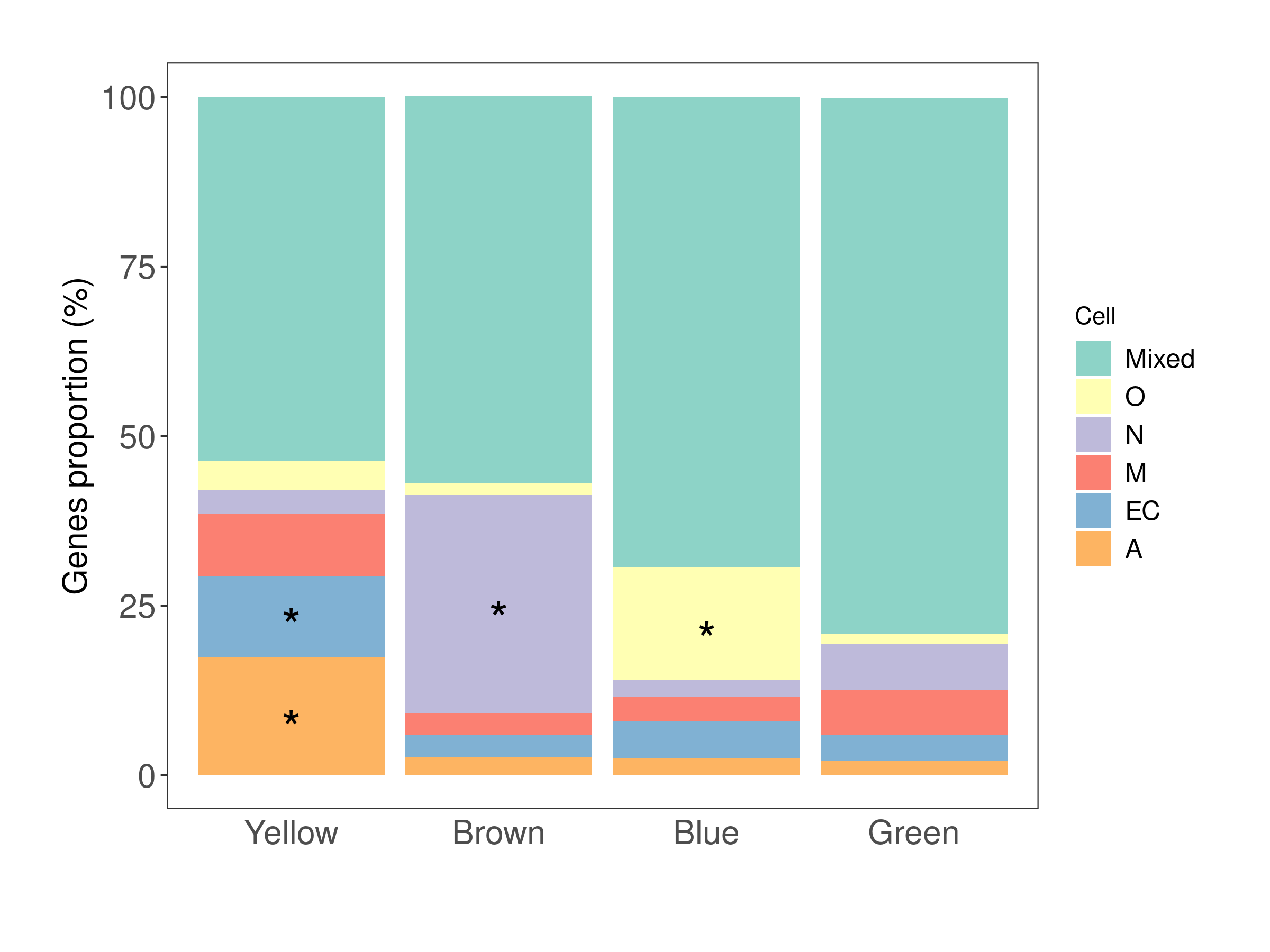

**Figure S22.** Plot representing the *Zsummary* statistics (x-axis) and the module size (y-axis). The *Zsummary* statistics indicates the preservation of a module in a test dataset. In this case, we are investigating whether the modules detected in C1 are preserved in C2. Focusing on the modules associated with the diagnostic status, the *Zsummary* statistics indicates a strong preservation for blue and brown module and a moderate preservation for the green module. No evidence of preservation was found for the yellow module. The gold module represent a random set of genes across the dataset.

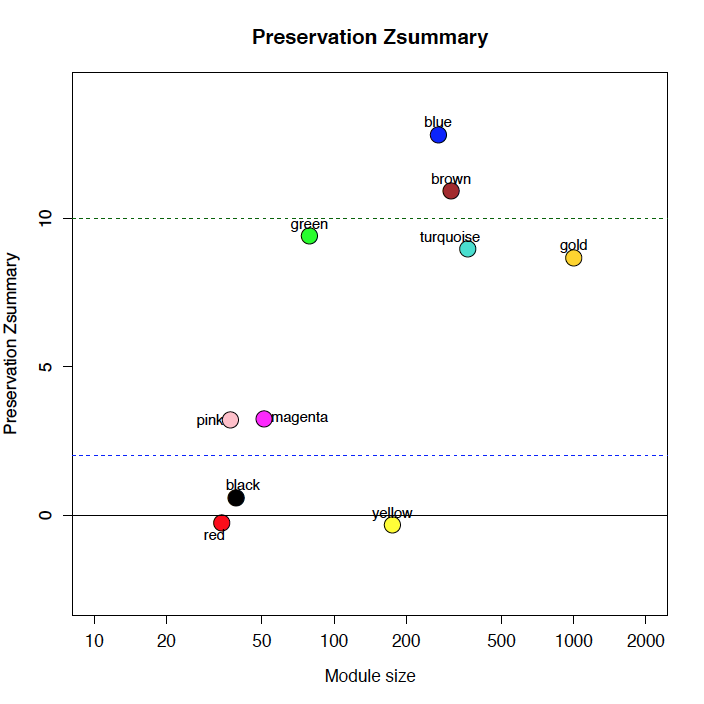

**References**

1.	Bennett DA, Schneider JA, Arvanitakis Z, S. Wilson R (2012) Overview and Findings from the Religious Orders Study. Curr Alzheimer Res 9:628–645. doi: 10.2174/156720512801322573

2.	Allen M, Carrasquillo MM, Funk C, Heavner BD, Zou F, Younkin CS et al. (2016) Human whole genome genotype and transcriptome data for Alzheimer’s and other neurodegenerative diseases. Sci Data 3. doi: 10.1038/sdata.2016.89

3.	Benjamini Y, Hochberg Y (1995) Benjamini Y, Hochberg Y. Controlling the false discovery rate: a practical and powerful approach to multiple testing. J R Stat Soc B 57:289–300. doi: 10.2307/2346101

4.	Cook RD (1977) Detection of Influential Observation in Linear Regression. Technometrics. doi: 10.1080/00401706.1977.10489493

5.	Dobin A, Davis CA, Schlesinger F, Drenkow J, Zaleski C, Jha S, Batut P, Chaisson M, Gingeras TR (2013) STAR: Ultrafast universal RNA-seq aligner. Bioinformatics 29:15–21. doi: 10.1093/bioinformatics/bts635

6.	Durinck S, Moreau Y, Kasprzyk A, Davis S, De Moor B, Brazma A, Huber W (2005) BioMart and Bioconductor: A powerful link between biological databases and microarray data analysis. Bioinformatics 21:3439–3440. doi: 10.1093/bioinformatics/bti525

7.	Ewels P, Magnusson M, Lundin S, K??ller M (2016) MultiQC: Summarize analysis results for multiple tools and samples in a single report. Bioinformatics 32:3047–3048. doi: 10.1093/bioinformatics/btw354

8.	Greene CS, Krishnan A, Wong AK, Ricciotti E, Zelaya RA, Himmelstein DS, Zhang R, Hartmann BM, Zaslavsky E, Sealfon SC, Chasman DI, Fitzgerald GA, Dolinski K, Grosser T, Troyanskaya OG (2015) Understanding multicellular function and disease with human tissue-specific networks. Nat Genet. doi: 10.1038/ng.3259

9.	Langfelder P, Horvath S (2007) Eigengene networks for studying the relationships between co-expression modules. BMC Syst Biol 1:54. doi: 10.1186/1752-0509-1-54

10.	Langfelder P, Horvath S (2008) WGCNA: an R package for weighted correlation network analysis. BMC Bioinformatics 9:559. doi: 10.1186/1471-2105-9-559

11.	Langfelder P, Luo R, Oldham MC, Horvath S (2011) Is my network module preserved and reproducible? PLoS Comput Biol 7. doi: 10.1371/journal.pcbi.1001057

12.	Lee HK, Hsu AK, Sajdak J, Qin J, Pavlidis P (2004) Coexpression analysis of human genes across many microarray data sets. Genome Res 14:1085–94. doi: 10.1101/gr.1910904

13.	Liao Y, Smyth GK, Shi W (2014) FeatureCounts: An efficient general purpose program for assigning sequence reads to genomic features. Bioinformatics 30:923–930. doi: 10.1093/bioinformatics/btt656

14.	Love MI, Huber W, Anders S (2014) Moderated estimation of fold change and dispersion for RNA-seq data with DESeq2. Genome Biol 15:550. doi: 10.1186/s13059-014-0550-8

15.	Okonechnikov K, Conesa A, García-Alcalde F (2015) Qualimap 2: Advanced multi-sample quality control for high-throughput sequencing data. Bioinformatics 32:292–294. doi: 10.1093/bioinformatics/btv566

16.	Ordway GA, Szebeni A, Duffourc MM, Dessus-Babus S, Szebeni K (2009) Gene expression analyses of neurons, astrocytes, and oligodendrocytes isolated by laser capture microdissection from human brain: Detrimental effects of laboratory humidity. J Neurosci Res 87:2430–2438. doi: 10.1002/jnr.22078

17.	R Core Team (2016) R Development Core Team. R A Lang. Environ. Stat. Comput. 55:275–286

18.	Ritchie ME, Phipson B, Wu D, Hu Y, Law CW, Shi W, Smyth GK (2015) limma powers differential expression analyses for RNA-sequencing and microarray studies. Nucleic Acids Res 43:e47. doi: 10.1093/nar/gkv007

19.	Schroder MS, Culhane AC, Quackenbush J, Haibe-Kains B (2011) survcomp: an R/Bioconductor package for performance assessment and comparison of survival models. Bioinformatics 27:3206–3208. doi: 10.1093/bioinformatics/btr511

20.	Wang J, Duncan D, Shi Z, Zhang B (2013) WEB-based GEne SeT AnaLysis Toolkit (WebGestalt): update 2013. Nucleic Acids Res 41. doi: 10.1093/nar/gkt439

21.	Wang M, Beckmann ND, Roussos P, Wang E, Zhou X, Wang Q, Ming C, Neff R, Ma W, Fullard JF, Hauberg ME, Bendl J, Peters MA, Logsdon B, Wang P, Mahajan M, Mangravite LM, Dammer EB, Duong DM, Lah JJ, Seyfried NT, Levey AI, Buxbaum JD, Ehrlich M, Gandy S, Katsel P, Haroutunian V, Schadt E, Zhang B (2018) The Mount Sinai cohort of large-scale genomic, transcriptomic and proteomic data in Alzheimer’s disease. Sci data 5:180185. doi: 10.1038/sdata.2018.185

22.	Zaykin D V. (2011) Optimally weighted Z-test is a powerful method for combining probabilities in meta-analysis. J Evol Biol 24:1836–1841. doi: 10.1111/j.1420-9101.2011.02297.x

23.	Zhang B, Horvath S (2005) A general framework for weighted gene co-expression network analysis. Stat Appl Genet Mol Biol 4:Article17. doi: 10.2202/1544-6115.1128

24.	Zhang Y, Chen K, Sloan SA, Bennett ML, Scholze AR, O’Keeffe S, Phatnani HP, Guarnieri P, Caneda C, Ruderisch N, Deng S, Liddelow SA, Zhang C, Daneman R, Maniatis T, Barres BA, Wu JQ (2014) An RNA-Sequencing Transcriptome and Splicing Database of Glia, Neurons, and Vascular Cells of the Cerebral Cortex. J Neurosci 34:11929–11947. doi: 10.1523/JNEUROSCI.1860-14.2014
